# NEBULA101: an open dataset for the study of language aptitude in behaviour, brain structure and function

**DOI:** 10.1101/2024.08.27.609682

**Authors:** Alessandra Rampinini, Irene Balboni, Olga Kepinska, Raphael Berthele, Narly Golestani

## Abstract

This paper introduces the “NEBULA101 - Neuro-behavioural Understanding of Language Aptitude” dataset, which comprises behavioural and brain imaging data from 101 healthy adults to examine individual differences in language and cognition. Human language, a multifaceted behaviour, varies significantly among individuals, at different processing levels. Recent advances in cognitive science have embraced an integrated approach, combining behavioural and brain studies to explore these differences comprehensively. The NEBULA101 dataset offers brain structural, diffusion-weighted, task-based and resting-state MRI data, alongside extensive linguistic and non-linguistic behavioural measures to explore the complex interaction of language and cognition in a highly multilingual sample. By sharing this multimodal dataset, we hope to promote research on the neuroscience of language, cognition and multilingualism, enabling the field to deepen its understanding of the multivariate panorama of individual differences and ultimately contributing to open science.

## Background & Summary

### Individual differences in language

This paper describes the “NEBULA101 - Neuro-behavioural Understanding of Language Aptitude” dataset. The dataset collects behavioural and brain imaging data of 101 healthy adults for the study of individual differences in language and cognition.

Human language is a complex behaviour, and crucial to understanding its workings is the fact that individuals differ in the way they manifest this and other cognitive skills^1^. The science of individual differences has expanded in recent years, thanks to a more integrated, less modular take on cognition: by combining the study of behaviour and the brain in a deep phenotyping approach, mindful of individual differences, researchers can gain a more comprehensive understanding of complex cognitive functions, language included^2^.

### Language aptitude

One aspect of language shown to display large individual differences is language *aptitude.* Language aptitude was originally proposed to explain why some people display remarkable abilities when learning additional languages ^3–5^. We use the term *additional* language for any language that is not the (or one of the) individual’s *first* language(s), i.e., language(s) to which they were exposed from birth. The term “additional” thus includes *second languages* (e.g. in the context of migration), *foreign languages* (e.g. in classroom learning), or *third/additional languages* of multilinguals who use more than two.

Initially, researchers viewed language aptitude as a stable trait, comprising phonetic coding, grammatical sensitivity, inductive learning, and rote learning abilities. These skills were seen as componential, in the climate of the first cognitive revolution, where the brain-mind-behaviour interaction was seen as the execution of algorithms operating within cognitive modules^6^: in this view, phonetic coding was at the core of production and perception of speech sounds, grammatical sensitivity underlay the capacity to identify structure in language (i.e. morphosyntax), inductive learning supported the generalisation of language rules from the input, and rote learning skills the construction of vocabulary via routinary use. Research has since suggested a more parsimonious structure, combining grammatical sensitivity and inductive learning into a global language analytic ability^7^.

Nowadays, with advances in neuroscience and experimental psychology methods, the language aptitude construct overall still describes a set of skills operating *across* the hierarchy of all language components, from lower to higher levels of complexity. What has changed is the way we conceptualise language itself, which has also changed the way we view language aptitude: in the second wave of the cognitive revolution, thanks to usage-based linguistics and neural-network psychology, language is seen as a complex adaptive system, rather than a set of rigid structures and fixed operations^8^, and one of the interacting components of human cognition, rather than an isolated function. Language aptitude is part of this dynamic: it can vary with age^9–11^ or with multilingual experience (likely increasing meta-linguistic awareness^12–14^, with patterns yet to be clarified^15,16^), and ultimately, with cognition more generally. In this context, we view the brain as a network of interrelated functions giving rise to complex behaviours, including language: to this end, this dataset comprises extensive general and domain-specific cognition tests, together with brain measures. The underlying concept of an integrated mind, arising from an integrated brain, is at the core of our methodological choice and of our first exploration of these data with graph theoretical methods^17^.

Mindful that any quantitative analysis will always require some degree of operationalisation (i.e. to derive scores from tests, we need to identify components of scientific constructs that such tests might tap into), our position is that this view of language as a dynamic system branching out and connecting to more general mechanisms holds promise for better understanding language itself and the human cognitive system more generally.

### The importance of research on the multilingual brain

The views expressed above are pivotal to the study of multilingualism and its behavioural and brain dynamics. In today’s globalised society multilingualism and multicompetence (i.e. using and knowing multiple languages) are becoming normalised^18^. There is, however, a theoretical issue with defining a sociolinguistic construct such as that of any “-lingualism”, due to its multidimensionality^19^, without incurring in stereotypes (e.g. the definition of *native-speakerism*) or perpetrating problematic practices, such as that of selecting certain profiles^20^ or matching “the unmatchable” based on intrinsically tricky parameters (e.g. the number of languages^21^). Nonetheless, we need to better define these concepts, as they constitute important characteristics of our present-day society. Knowing and using multiple languages demands a fundamental cognitive (re)organisation^18^, with several psycho-neurobiological correlates that are somewhat hard to reconcile in a comprehensive view^22^. Therefore, we must seek to imagine language competence and use in newer, more naturalistic, and multidimensional ways to better understand their influence on the brains (and lives) of language users.

To this aim, we believe that multimodal datasets with rich phenotypical information, such as the one presented here, are a step forward in this direction. Language aptitude, viewed as embedded in (and interacting with) cognition might be one of the engines driving multilingualism (in its many forms), and understanding its underpinnings might ultimately influence the way we view, use, and even teach language(s). In this context, Switzerland holds a special status as a country with four official languages and as a destination for expatriates from all over the world, including users of Minority, Indigenous, Non-standard, and Dialect (MIND) varieties. These languages have often been disregarded not only by researchers but by users themselves, who might not even recognise their multilingual status when knowing an additional MIND language^20^. However, these *do* appear in the multicompetent panorama of many of our participants, when asked explicitly (see Figure 2).

These issues have recently been tackled in work calling for a more diverse view of cognitive science in general^23^ and neurolinguistics^24^ in particular, and underlining the contribution *non-English* (Romance^25^, non-Indo-European^26^ as well as MIND ^20,21^) languages to the field. The ongoing discourse on language policies and teaching^27–29^, as well as the thriving field of instructed language learning research, especially in the Swiss context^30^, are just a part of the puzzle. What is still lacking in the psycho- and neurolinguistics of multilingualism is a naturalistic perspective rooted in the brain itself, also likely due to the intrinsic difficulty of quantifying language use in a nonparametric and dynamic way. Humans, however, possess a skill, language, that varies at all levels, challenging us to consider the remarkable plasticity of advanced human abilities by harnessing language diversity as a tool for advancing cognitive science^31^. Thus, while leveraging French knowledge and fluency of our participants - a condition necessary for performance comparability - this dataset also captures and documents their linguistic diversity and multicompetence. This comprehensive documentation has the potential to facilitate investigations into how such diversity relates to both the phenotypical (behavioural) and endophenotypical (brain structural and functional) characteristics of individuals.

### Aptitude for language(s) and individual differences in other domains of cognition

The construct of language aptitude was developed in the domain of foreign language learning, as explained, but the idea that aptitude only manifests in foreign languages has now been surpassed, since individual differences can be observed in first language skills too, even if it is harder to pinpoint them and isolate them from experiential factors^32^. Moreover, such individual differences might co-exist more globally with individual differences in other domains of cognition^17,33^, giving rise to “neurocognitive profiles” of language aptitude involving the mnemonic domain, fluid reasoning, auditory abilities, and even musicality^34^. It is therefore relevant to ask ourselves whether language, in this modernised and dynamic view, is part of a positive manifold ^35,36^ originating from the beneficial interactions between cognitive processes, as we proposed in a recent analysis of part of these data^17^. Both the stable and malleable (or *plastic*) features of the human cognitive system are fundamental to such interactions.

### Stability and malleability, predisposition and experience

This dataset presents cross-sectional data. Nonetheless, to understand the nature of the language (aptitude) construct, it is important to consider it as the result of a complex interaction between stable traits and malleable states, the first ascribed to genetic predisposition and the latter to environmental influences (what we generally define as “experience”)^37–39^. While we provide a *snapshot* of individual profiles at a given moment in time, in order to formulate relevant questions on (and interpretations of) these data, it is important to remember that the observed measures arise from both stable traits and from malleable skills, or states^40^. Cognitive activity^41–44^, including language learning and use^45–48^, and brain *plasticity* are intrinsically related. Changes in regional activity, network connectivity, or morphology, arising from underlying molecular, neurobiochemical and other changes^41^, subtend this state of malleability in brain function and structure^49^, both during development and in skill learning. However, brain functional and structural architecture is also highly polygenic (i.e. controlled by the complex interaction of multiple genes, which in turn are expressed in multiple variants across individuals), and in addition, different cortical loci and tissue features (thickness, surface area) are affected to different degrees by genetics^50^. Further, genetic factors also likely influence the degree to which neuroplasticity manifests^51^.

### The open science of language aptitude

Language aptitude has come a long way since its study was confined to the foreign language classroom, and research restricted to military and government access^52^. The construct now encompasses some of the most promising avenues for research on the human cognitive system more generally: the roles of predisposition and experience, the nature of neuroplasticity and the integrated and multivariate organisation of cognitive domains. Because of its relevance for studying questions on language and cognition more generally (as well as investigating the role of the environment, e.g. the experience of multilingualism), this development in the conceptualisation of language aptitude is of particular interest to the ever-growing world of Findable, Accessible, Interoperable, and Reusable (FAIR) neuroscience data^53^, which has faced a substantial growth in the last 15 years^54^, and possibly as many challenges^55,56^. FAIR principles are accelerating our comprehension of the human brain^57^. Concurrently, the existence of standardised protocols such as the Brain Imaging Data Structure (BIDS)^58^ supports this recent strive towards data and code accessibility, ensuring that data is organized and its analysis reproducible, enabling more efficient and effective use of shared datasets and streamlining data preparation, thus allowing scientists to focus more on discovery.

Mindful that the panorama of shared neuroimaging data is vast and that datasets might be scattered around repositories of the commercial and institutional type, focusing on either a specific population (clinical, paediatric) or modality^59^, we searched for openly available data similar to ours (MRI and behavioural data from healthy adult participants focusing on language) on OpenNeuro^60^, one of the most recent, easy to access and growing neuroimaging databases. At the time of writing, a search on http://openneuro.org for BIDS-valid MRI raw/derivative datasets with more than 50 healthy adults, including the keyword ‘language’ and accompanied by a published or pre-registered data descriptor, yields the following entries (Table 1), with two of the datasets (ds004215, ds000243) being listed but having no relationship with language (and thus not being included in the below table).

**Table 1.** Available neuroimaging datasets on language with more than 50 healthy participants, validated in BIDS. (*)Possibly including or revising ds002382 but no information on dataset version is provided. (**) No data descriptor available.

| OpenNeuro Accession number | Dataset name and reference | Modality | Number of participants | Complete dataset (all participants-measures) | Associated tasks/measures | Main topic |
| --- | --- | --- | --- | --- | --- | --- |
| ds003481<sup>61</sup> | A Dataset to Study Pragmatic Language and Its Underlying Cognitive Processes<sup>62</sup> | sMRI, fMRI, behaviour | 145 | No | Comprehension of metaphorical phrases and proverbs<br>Recognition and emotional categorisation of speech acts<br>Lexical-semantic processing | Language pragmatics |
| ds004765<sup>63</sup> | Relationship between resting state functional connectivity and reading-related behavioural measures in 69 adults<sup>64</sup> | sMRI, fMRI (resting state), DWI | 69 | Yes | Word/nonword reading<br>Spelling<br>Lexical decision<br>Spoonerisms<br>Rapid automatised naming<br>Non-word repetition<br>Vocabulary | Reading |
| ds002382<sup>65</sup> | Age-related differences in auditory cortex activity during spoken word recognition<sup>66</sup> | sMRI, fMRI | 61 | Yes | Word listening<br>Word repetition | Word processing |
| ds004285<sup>67</sup> | Listening task* | sMRI, fMRI | 78 | Yes | Word repetition | Word processing |
| ds004073<sup>68</sup> | Comparing language lateralisation using fMRI and fTCD<sup>69</sup> | fMRI, functional transcranial doppler sonography (fTCD) | 51 | No (minimally missing information) | Word generation<br>Sentence generation<br>Phonological decision<br>Word comprehension<br>Sentence comprehension<br>Syntactic decision | Language lateralisation |
| ds001747<sup>70</sup> | Exploring the Resting State Neural Activity of Monolinguals and Late and Early Bilinguals<sup>71</sup> | sMRI, fMRI (resting state) | 92 | Yes | L1 and L2 (where applicable) proficiency<br>Language background | Bilingualism |
| ds001796<sup>72</sup> | Bilingualism and the brain** | sMRI, fMRI (task-based), fMRI (resting state), DWI | 64 | Yes | Flanker task | Bilingualism |
| ds002345<sup>73</sup> | Narratives: fMRI data for evaluating models of naturalistic language comprehension<sup>74</sup> | sMRI, fMRI | 345 | No | Naturalistic story listening | Language perception |
| ds003643<sup>75</sup> | Le Petit Prince: A multilingual fMRI corpus using ecological stimuli<sup>76</sup> | sMRI, fMRI | 112 | Yes | • Naturalistic story listening | Language perception |

A similar, broad search on Google Scholar for data descriptors associated with openly available language datasets (“open AND mri dataset AND language”) yielded more datasets: the MOUS (Mother Of Unification Studies)^77^, a dataset comprising sMRI, DWI, resting state and task-based fMRI and MEG in 204 participants assigned to either sentence reading or listening; the Alice dataset^78^, where 75 participants listened to the same chapter of Alice in Wonderland during fMRI or during EEG. A third dataset presents a large quantity of data collected in fewer participants with state-of-the-art technology: this is the high-resolution, 7T fMRI Forrest Gump database^79^, during which participants watched the ‘Forrest Gump’ movie. This data can be used to study naturalistic language processing, even though the study did not primarily focus on language. Then, the search yielded the LanA dataset, a probabilistic language atlas derived from brain data in more than 800 individuals^80^. Finally, the search also yielded two speech production datasets of vocal tract MRI^81,82^.

At the behavioural level there is more heterogeneity in the type and quantity of shared data^59^. The origins of such heterogeneity have been discussed for quite some time. In a Nature Neuroscience perspective article from ten years ago, the challenges linked to standardisation and accessibility of behavioural data were already discussed^83^. Importantly, behaviour was defined as a complex, highly dimensional, dynamic, and interconnected phenomenon without distinct separable scales, and this was discussed as being one of the leading causes for lack of standardisation and scarce FAIR compliance. The authors insisted on its foundational and unifying nature, and called for improved standards: “Behaviour (…) is the principal function of the brain. (…) Copious, quantitative and open behavioural data has the potential (…) to solidify the foundations of other [disciplines], including neuroscience” ^83^ (p.1455). However, even in the cited datasets, which represent timely and relevant steps towards accessibility of neuroscience data, the phenotypical (behavioural) information being shared beyond demographics is still relatively less prominent than the endophenotypical (neural) data: oftentimes, only data from the behavioural tasks being performed during fMRI are available. Moreover, even when behaviour was tested outside the scanner, if the original project did not focus on *both* phenotypical and endophenotypical data and on a specific topic (such as, in the cited examples, pragmatics^84^ or reading^64^), the accompanying behavioural information is scarce. Two notable, recent exceptions in the field of individual differences in language are represented by a behavioural dataset including 33 measures from 112 adult Dutch speakers^85^, and by recent work by Berthele and colleagues in children^86^, both representing an important milestone for shared behavioural data.

Given the lack of standardisation in the way we administer behavioural tasks outside the scanner, compared to fMRI task delivery, it seems daunting to force the structure of behavioural paradigms and log files produced by a plethora of software and online platforms to accommodate information within the structure required by the BIDS standard, a difficulty that we encountered in our work, as well. Some initiatives, such as the Behaverse project^85^, are proposing data structures which, they claim, can accommodate phenotypical information better than what BIDS can do. However, it is important to note that any individual differences dataset including both phenotypical and endophenotypical information will have the added strength of multimodality, and to date, BIDS is the only data structure protocol that can accommodate *both* in a relatively straightforward way, even if it requires extra (and sometimes *post-hoc*) work to prepare the materials intended to be shared. Finally, we must note that when sharing mixed raw and derivative datasets, behavioural data can be easily included as derivatives, these having a more liberal structure, lifting from the end-user the load of reprocessing and calculating basic scores starting from item-level data. This is the route we chose for this dataset, including raw (minimally processed) phenotypic and behavioural data with varying underlying structures, coming from questionnaires and tasks respectively, crucially accompanied by their derivate scores.

In sum, the NEBULA101 dataset aims to promote the study of individual differences in language to better understand a multivariate cognitive system, via the sharing of a truly multimodal dataset in an adequately sized participant sample^87^. We provide sMRI, DWI, task-based and resting state fMRI in 101 individuals, together with broad phenotypic and behavioural data on linguistic but also on non-linguistic, domain general and domain specific tasks, including cognitive and perceptuomotor tasks. This includes measures of all language aptitude components from phonetics to syntax, measures of reading and reading mediators (e.g. phonological awareness), domain-general cognitive skills, numerical processing, musicality and musical experience and rich multilingual language experience measures. By providing these data to the public domain, we hope to contribute new discoveries to the over-arching construct of language (aptitude), embracing the components of individual behavioural and neural phenotypes as widely as possible.

### Data types

Behaviour is the phenotype that can be related to, or unify, genetics, neural architecture, neural activity, body structure, physical limitations, and environmental factors^83^. Given the importance of behavioural data in exploring individual differences at the neural level, this dataset includes 28 scores derived from 8 questionnaires, and 74 behavioural measures derived from 21 tasks. Functional neuroimaging data provide information about the neural correlates of cognitive processes, allowing to elucidate how the brain supports specific behaviours and skills. Here, we provide resting-state and task-based functional imaging (fMRI) data, the latter obtained during a language localiser^88^. Finally, NEBULA101 also includes anatomical T1-weighted and diffusion-weighted (DWI) imaging of the brain, which will allow to study brain anatomy and white-matter structural connectivity, to shed light on the brain structural correlates of linguistic behaviours, aptitudes and experiences.

An overview of all measures provided in the NEBULA101 dataset is provided in Table 2. In the Supplementary Information file, Table S1 contains more details specific to the version of the tests used in this dataset, such as any adaptations, modality of administration and derivate scores.

**Table 2.**
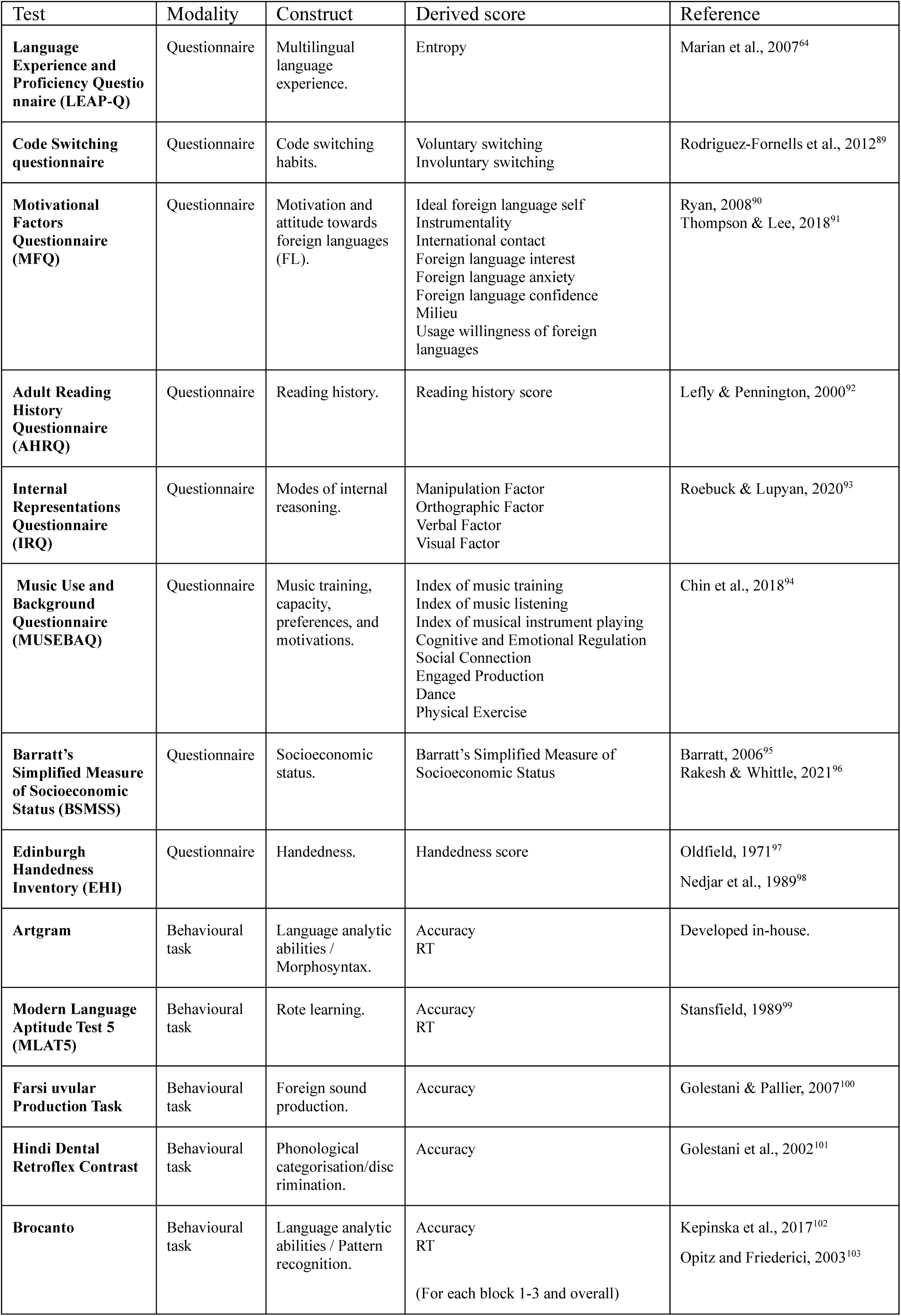
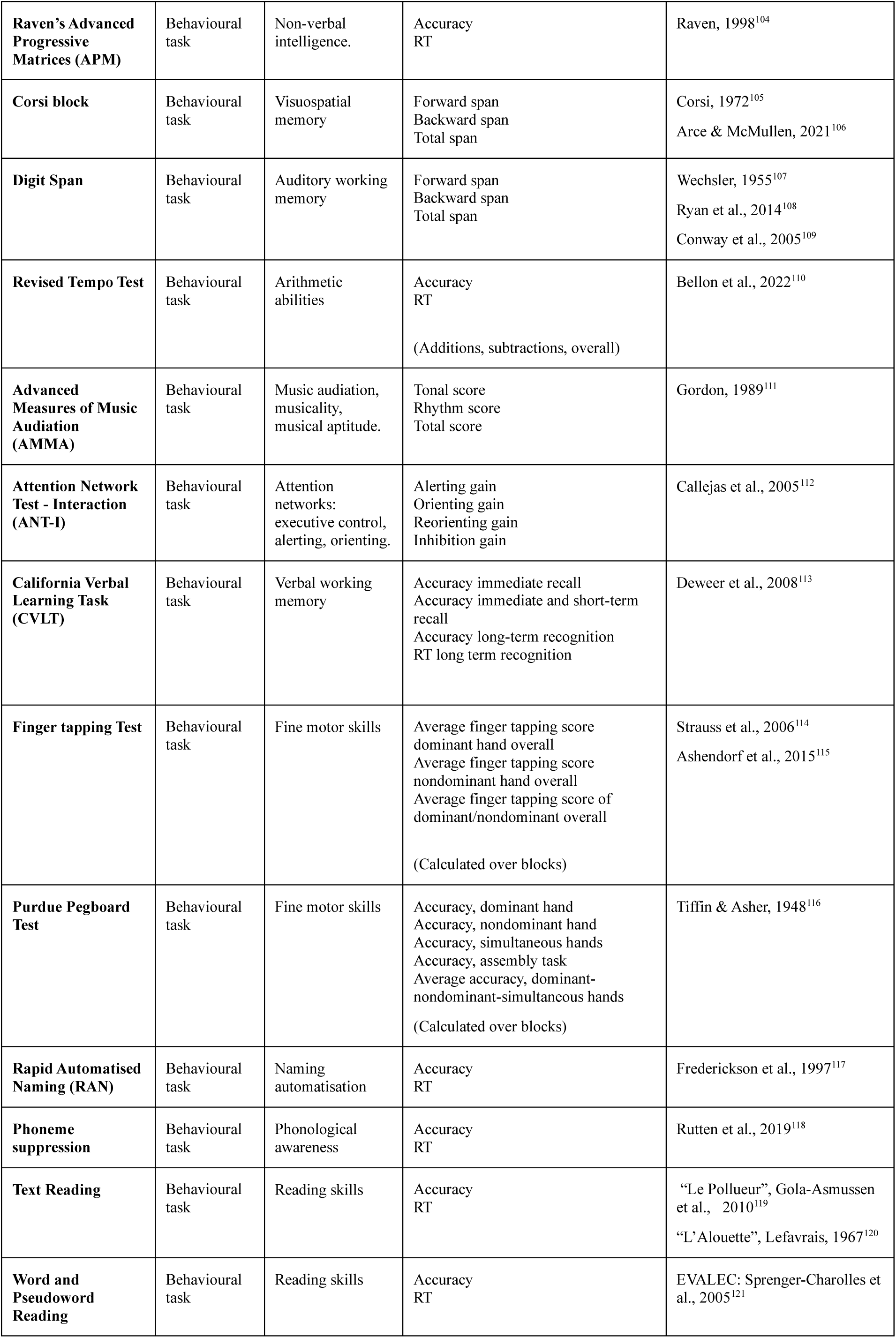
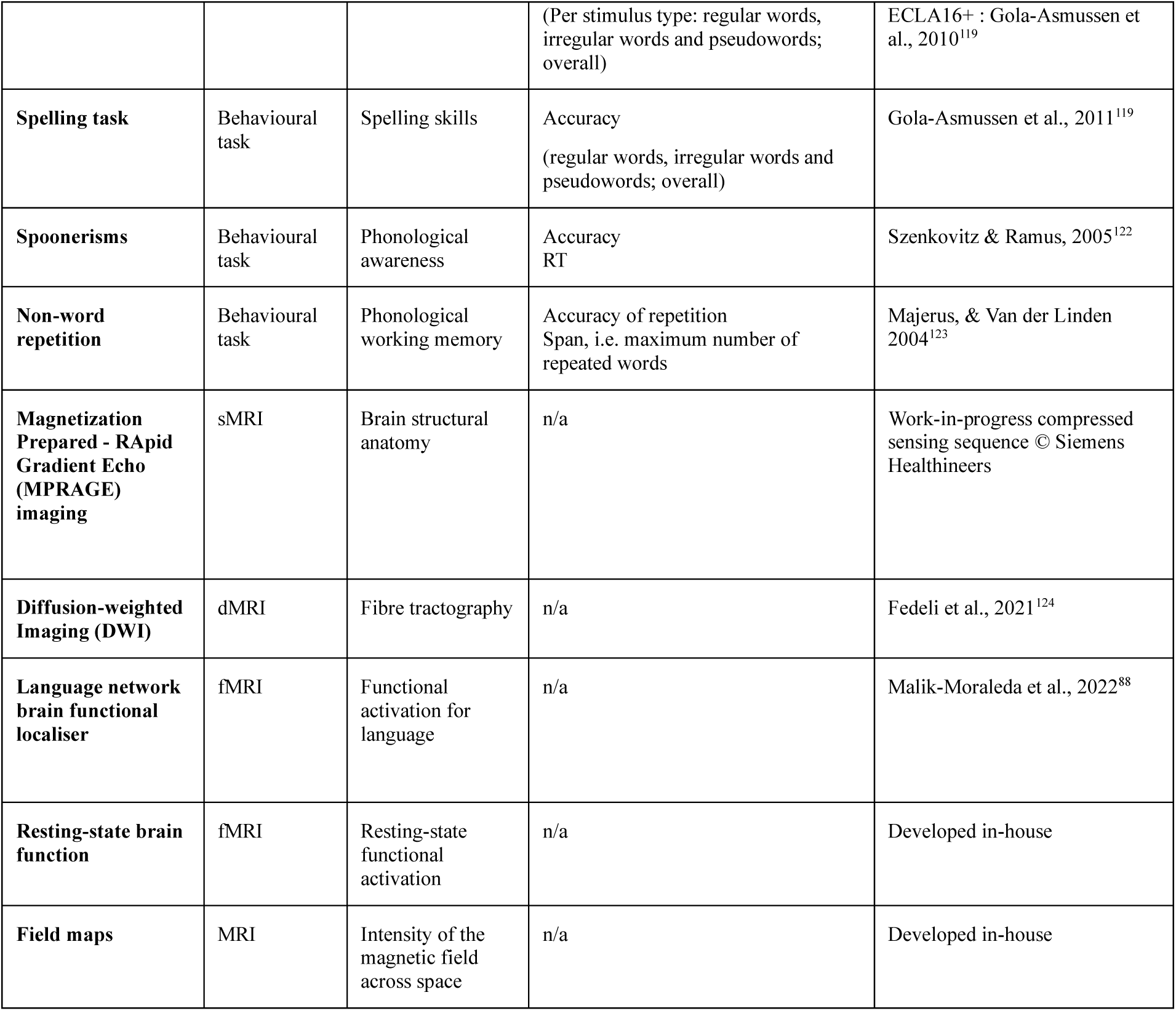
Overview of all tasks and modalities.

| Test | Modality | Construct | Derived score | Reference |
| --- | --- | --- | --- | --- |
| **Language Experience and Proficiency Questionnaire (LEAP-Q)** | Questionnaire | Multilingual language experience. | Entropy | Marian et al., 2007<sup>64</sup> |
| **Code Switching questionnaire** | Questionnaire | Code switching habits. | Voluntary switching<br>Involuntary switching | Rodriguez-Fornells et al., 2012<sup>89</sup> |
| **Motivational Factors Questionnaire (MFQ)** | Questionnaire | Motivation and attitude towards foreign languages (FL). | Ideal foreign language self<br>Instrumentality<br>International contact<br>Foreign language interest<br>Foreign language anxiety<br>Foreign language confidence<br>Milieu<br>Usage willingness of foreign languages | Ryan, 2008<sup>90</sup><br>Thompson & Lee, 2018<sup>91</sup> |
| **Adult Reading History Questionnaire (AHRQ)** | Questionnaire | Reading history. | Reading history score | Lefly & Pennington, 2000<sup>92</sup> |
| **Internal Representations Questionnaire (IRQ)** | Questionnaire | Modes of internal reasoning. | Manipulation Factor<br>Orthographic Factor<br>Verbal Factor<br>Visual Factor | Roebuck & Lupyan, 2020<sup>93</sup> |
| **Music Use and Background Questionnaire (MUSEBAQ)** | Questionnaire | Music training, capacity, preferences, and motivations. | Index of music training<br>Index of music listening<br>Index of musical instrument playing<br>Cognitive and Emotional Regulation<br>Social Connection<br>Engaged Production<br>Dance<br>Physical Exercise | Chin et al., 2018<sup>94</sup> |
| **Barratt's Simplified Measure of Socioeconomic Status (BSMSS)** | Questionnaire | Socioeconomic status. | Barratt's Simplified Measure of Socioeconomic Status | Barratt, 2006<sup>95</sup><br>Rakesh & Whittle, 2021<sup>96</sup> |
| **Edinburgh Handedness Inventory (EHI)** | Questionnaire | Handedness. | Handedness score | Oldfield, 1971<sup>97</sup><br>Nedjar et al., 1989<sup>98</sup> |
| **Artgram** | Behavioural task | Language analytic abilities / Morphosyntax. | Accuracy<br>RT | Developed in-house. |
| **Modern Language Aptitude Test 5 (MLAT5)** | Behavioural task | Rote learning. | Accuracy<br>RT | Stansfield, 1989<sup>99</sup> |
| **Farsi uvular Production Task** | Behavioural task | Foreign sound production. | Accuracy | Golestani & Pallier, 2007<sup>100</sup> |
| **Hindi Dental Retroflex Contrast** | Behavioural task | Phonological categorisation/discrimination. | Accuracy | Golestani et al., 2002<sup>101</sup> |
| **Brocanto** | Behavioural task | Language analytic abilities / Pattern recognition. | Accuracy<br>RT<br>(For each block 1-3 and overall) | Kepinska et al., 2017<sup>102</sup><br>Opitz and Friederici, 2003<sup>103</sup> |
| **Raven's Advanced Progressive Matrices (APM)** | Behavioural task | Non-verbal intelligence. | Accuracy<br>RT | Raven, 1998<sup>104</sup> |
| **Corsi block** | Behavioural task | Visuospatial memory | Forward span<br>Backward span<br>Total span | Corsi, 1972<sup>105</sup><br>Arce & McMullen, 2021<sup>106</sup> |
| **Digit Span** | Behavioural task | Auditory working memory | Forward span<br>Backward span<br>Total span | Wechsler, 1955<sup>107</sup><br>Ryan et al., 2014<sup>108</sup><br>Conway et al., 2005<sup>109</sup> |
| **Revised Tempo Test** | Behavioural task | Arithmetic abilities | Accuracy<br>RT<br>(Additions, subtractions, overall) | Bellon et al., 2022<sup>110</sup> |
| **Advanced Measures of Music Audiation (AMMA)** | Behavioural task | Music audiation, musicality, musical aptitude. | Tonal score<br>Rhythm score<br>Total score | Gordon, 1989<sup>111</sup> |
| **Attention Network Test - Interaction (ANT-I)** | Behavioural task | Attention networks: executive control, alerting, orienting. | Alerting gain<br>Orienting gain<br>Reorienting gain<br>Inhibition gain | Callejas et al., 2005<sup>112</sup> |
| **California Verbal Learning Task (CVLT)** | Behavioural task | Verbal working memory | Accuracy immediate recall<br>Accuracy immediate and short-term recall<br>Accuracy long-term recognition<br>RT long term recognition | Deweert et al., 2008<sup>113</sup> |
| **Finger tapping Test** | Behavioural task | Fine motor skills | Average finger tapping score dominant hand overall<br>Average finger tapping score nondominant hand overall<br>Average finger tapping score of dominant/nondominant overall<br>(Calculated over blocks) | Strauss et al., 2006<sup>114</sup><br>Ashendorf et al., 2015<sup>115</sup> |
| **Purdue Pegboard Test** | Behavioural task | Fine motor skills | Accuracy, dominant hand<br>Accuracy, nondominant hand<br>Accuracy, simultaneous hands<br>Accuracy, assembly task<br>Average accuracy, dominant-nondominant-simultaneous hands<br>(Calculated over blocks) | Tiffin & Asher, 1948<sup>116</sup> |
| **Rapid Automatised Naming (RAN)** | Behavioural task | Naming automatisisation | Accuracy<br>RT | Frederickson et al., 1997<sup>117</sup> |
| **Phoneme suppression** | Behavioural task | Phonological awareness | Accuracy<br>RT | Rutten et al., 2019<sup>118</sup> |
| **Text Reading** | Behavioural task | Reading skills | Accuracy<br>RT | "Le Pollueur", Gola-Asmussen et al., 2010<sup>119</sup><br>"L'Alouette", Lefavrais, 1967<sup>120</sup> |
| **Word and Pseudoword Reading** | Behavioural task | Reading skills | Accuracy<br>RT | EVALEC: Sprenger-Charolles et al., 2005<sup>121</sup> |
|  |  |  | (Per stimulus type: regular words, irregular words and pseudowords; overall) | ECLA16+ : Gola-Asmussen et al., 2010<sup>119</sup> |
| **Spelling task** | Behavioural task | Spelling skills | Accuracy<br>(regular words, irregular words and pseudowords; overall) | Gola-Asmussen et al., 2011<sup>119</sup> |
| **Spoonerisms** | Behavioural task | Phonological awareness | Accuracy<br>RT | Szenkovitz & Ramus, 2005<sup>122</sup> |
| **Non-word repetition** | Behavioural task | Phonological working memory | Accuracy of repetition<br>Span, i.e. maximum number of repeated words | Majerus, & Van der Linden 2004<sup>123</sup> |
| **Magnetization Prepared - RApid Gradient Echo (MPRAGE) imaging** | sMRI | Brain structural anatomy | n/a | Work-in-progress compressed sensing sequence © Siemens Healthineers |
| **Diffusion-weighted Imaging (DWI)** | dMRI | Fibre tractography | n/a | Fedeli et al., 2021<sup>124</sup> |
| **Language network brain functional localiser** | fMRI | Functional activation for language | n/a | Malik-Moraleda et al., 2022<sup>88</sup> |
| **Resting-state brain function** | fMRI | Resting-state functional activation | n/a | Developed in-house |
| **Field maps** | MRI | Intensity of the magnetic field across space | n/a | Developed in-house |

## Methods

### Participants

According to the Organisation for Economic Cooperation and Development (OECD), approximately 40% of Swiss individuals aged 25 to 34 possess upper secondary or post-secondary non-tertiary education, around 50% have attained tertiary education, and about 10% have below-secondary education^125,126^. Additionally, Switzerland is recognized for its linguistic diversity, as we have extensively discussed in the above paragraphs. The Federal Population Census 2022 Structural Survey^127^ indicates that French is spoken by 22.8% of the population. In the survey, 70.1% and 23.4% of the population declared that they speak other national (German, Italian, and/or Romansch) or non-national languages (participants could declare more than one language, therefore there is a degree of overlap in these data). Considering that any study examining the interaction between diverse cognitive domains and language skills will always be somewhat culture-dependent^128^, we chose our target participants with the aim to test at least 100 healthy, relatively multilingual individuals having French as their first or dominant language, which is the primary language of the Canton where the data collection took place.

We recruited 104 adult participants who matched these criteria from the Geneva area, the surrounding French-speaking cantons of Switzerland and neighbouring France through flyers and online advertisements. Participants provided their consent for disclosing their medical history and filled in an online screening survey. To safeguard coherence and avoid confounding factors within the sample, prospective participants were excluded *a priori* if they had musical or simultaneous interpreting professional qualifications (known to interact with language); vision defects that could not be corrected; body implants incompatible with MRI or known claustrophobia; neurological or psychiatric conditions; traumatic head injuries with loss of consciousness; ongoing pregnancies; and past or present serious illnesses that required invasive and/or continued medical treatment (such as cancer, chronic and/or autoimmune diseases). Participants with MRI-conditional implants were evaluated on an individual basis upon providing further documentation. Participants with diagnosed developmental dyslexia, as well as those who reported knowing more than 10 languages with a self-reported proficiency equal to or higher than 4 out of 10 in reading, speaking and listening, are not included in this dataset. Figure 1 reports the number of declared languages for the final sample.

**Figure 1.**
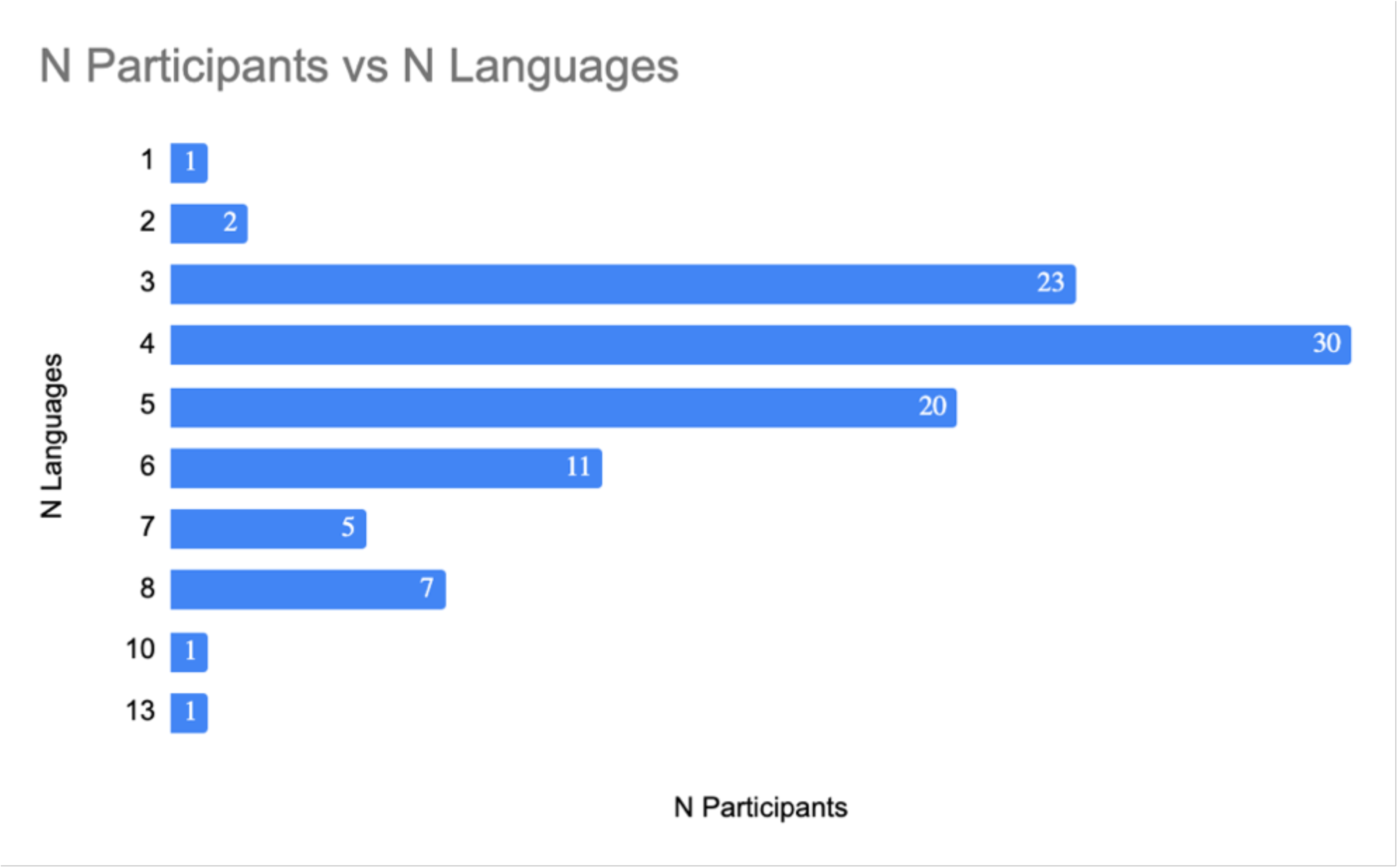
Frequency distribution of the number of reported languages in the final sample (N = 101).

Once eligibility was established, all participants provided a signed informed consent to all subsequent experimental procedures, including anonymised data reuse for open science. One participant subsequently withdrew their consent to data sharing and two were not able to undergo brain imaging due to claustrophobia. Therefore, the final sample includes 101 individuals (*M*_age_= 23.35 years, *SD* = 4.08, 68F, *M_education_*= 15.34 years, *SD* = 2.35). At study completion, all participants received monetary compensation, an image of their brain and a simplified report on their performance on the behavioural tests. Language background and social status show that our participants were quite representative of the well educated and economically stable Swiss society. This information is shown in Figure 2, with social status represented as a cumulative measure derived from the education and job category of the participant, as well as those of their family of origin and of their partner, if present, calculated with Barratt’s Simplified Measure of Social Status (BSMSS)^95^.

**Figure 2.**
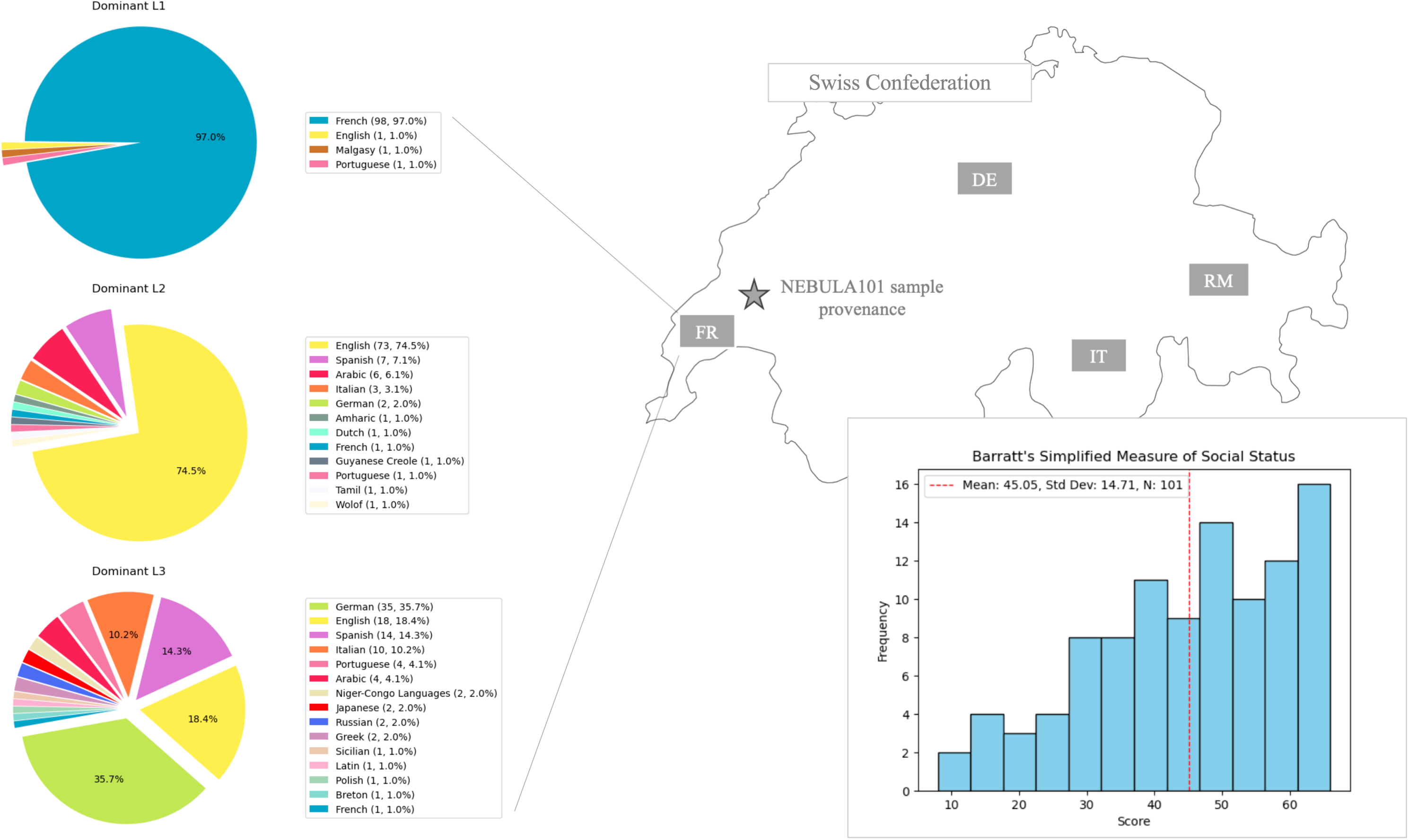
Participants’ language background up until their third language (left, languages coded by colour) and Socioeconomic status (bottom right) measured via Barratt’s simplified measure of social status^95^.

### Data Collection

#### Data collection

All interactions and documents provided to participants were in French. Data were collected by six individuals who were either first-language French users, or who had learned French as their second/additional language at an advanced level. All procedures were approved by the Geneva Cantonal Ethical Commission (CCER Protocol N. 2021-01004) prior to the beginning of the study. All participants signed a waiver for anonymised data release in the public domain, and no identifying information was retained, to the best of our knowledge.

Data were collected in two online and two in-person sessions, hereafter named Session 1, 2, 3 and 4, which always occurred on different days but in the same order (1-4) between July 2022 and June 2023. Session 1 was unsupervised, and required participants to fill out online questionnaires. Session 2 involved online behavioural data collection, and was supervised by an experimenter. During Session 3 we collected more behavioural data, this time in person. Finally, in Session 4 we collected neuroimaging (MRI) data.

#### Behavioural testing

Using established and published tests in cognitive psychology is crucial for addressing the reproducibility crisis and curbing the proliferation of one-off tests, for ensuring that findings are accurate and that they can be compared and replicated by other researchers^129^. Therefore, where possible, we chose existing questionnaires and behavioural tasks, with the exception of an explicit morphosyntax learning test, which we developed and piloted ourselves due to the lack of such a test in the field (see section “Pilot study”). All instructions and tests were delivered in French. Where the French version of a test was not available, it was *adapted* and checked by at least one first-language French user in the team. Questionnaire instructions (Session 1) were presented in written form. Behavioural task instructions (Sessions 2 and 3) were read by a commercial natural reader (https://www.naturalreaders.com/commercial/read), using the voice “Renee France” (if not otherwise specified), at speed −1. fMRI task instructions were given by the experimenter via a microphone. All technical alterations and task adaptations were related to 1) language of delivery, or 2) adapting a paper & pencil test for computer-based delivery. All the above information is thoroughly documented for each test in the Supplementary Information file, Table S1. An overview of the data collection structure is given in Figure 3 and an extensive theoretical explanation of the tests included in sessions 1-3 can be found in our recent exploratory work on the behavioural correlates of language aptitude, which included this sample^17^.

**Figure 3.**
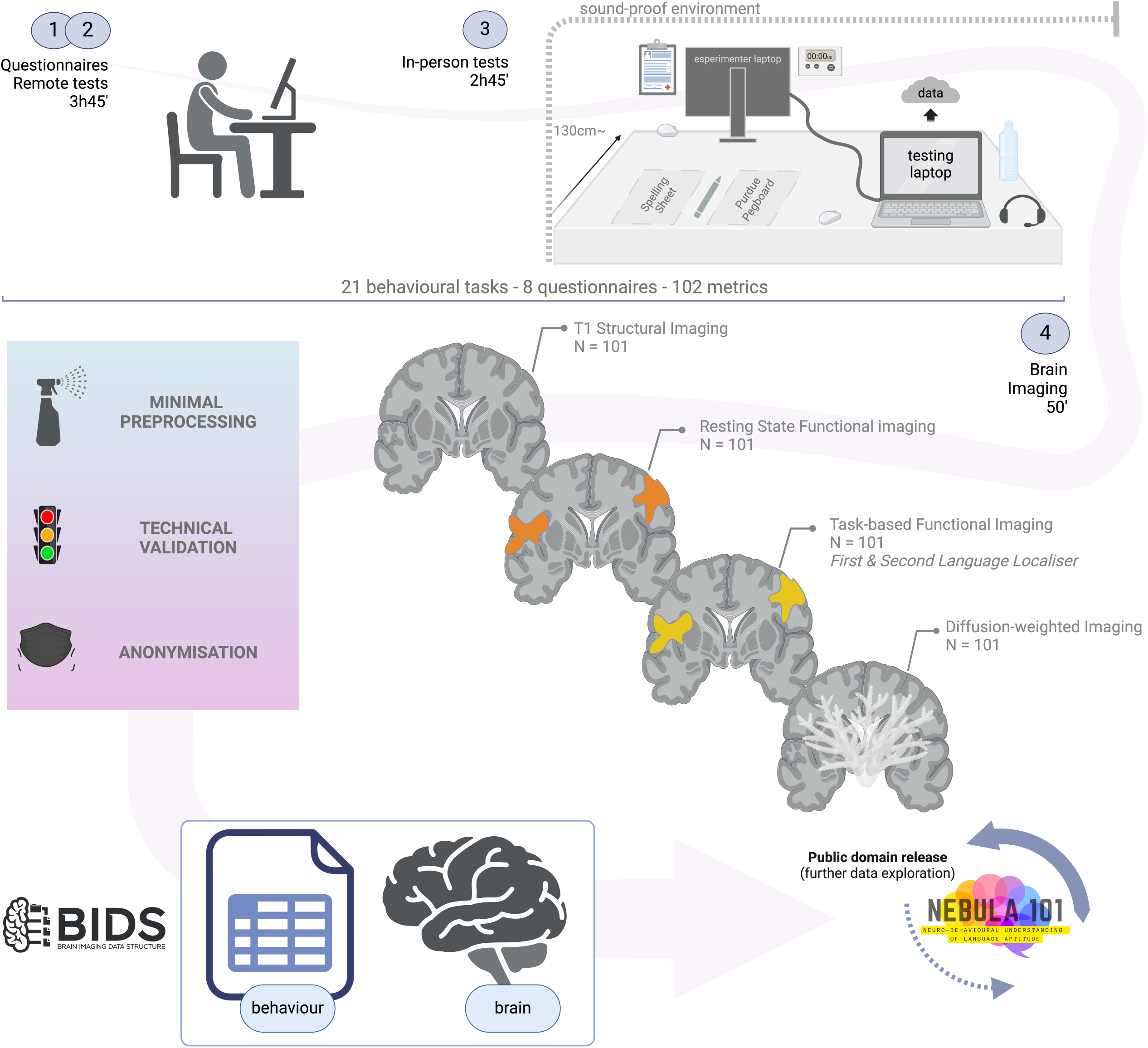
Data acquisition and processing structure (session duration is approximate for behavioural data acquisition due to individual variability in task completion times). [Illustration created with BioRender.com].

#### Pilot study

Before data collection, an explicit grammar learning task called ArtGram was developed and pilot-tested. ArtGram, designed for adults, extends the PLAB4 task used for language aptitude testing in children and adolescents, for which we could not find an equivalent in adults^130,131^. The test involves learning an artificial, declensional language lexicon with inflected sample sentences, followed by a self-paced multiple-choice, speeded translation task with novel sentences, as described in Table 2 and more extensively in Rampinini et al., (2024)^17^. The pilot study aimed to assess: (1) the task’s feasibility, (2) the reliability of the online platform, (3) redundancy with an implicit grammar learning task (Brocanto). Twenty first-language French speakers (11F, *M*_age_ = 27.65, *SD* = 8.9) without language or reading disorders participated via video conference. Results showed an unsignificant correlation of *r*(18) = .44 between the two grammar tasks.

#### Session 1: questionnaires

Participants initially filled out a series of questionnaires online using Qualtrics XM©. The questionnaire sequence was fixed: first came the Edinburgh Handedness Test and the BSMSS questionnaire, completed upon recruitment. Then came the demographics, MFQ, IRQ, AHRQ, MUSEBAQ, Code Switching and LEAPQ (see Table 2). When available from the questionnaire manual, automatic scoring was implemented in Qualtrics via the “Formula Field” and resulted in one column per score in the derivative questionnaire dataset, for each participant.

#### Session 2: online behavioural testing

During session 2, supervised behavioural data collection was conducted via Zoom©. Participants were guided through a demonstration video on how to share both their sound and entire screen, use wired headphones for reliable online measurements^132,133^, and ensure their microphone was functioning properly. Tasks were delivered through the Gorilla web interface^134^. Headphone and microphone tests from the Gorilla open materials section were mandatory before starting the task sequence. Researchers supervised the session, intervening only if technical issues arose. The system prevented participants from logging into the session from mobile phones or tablets, and only allowed the use of Mozilla Firefox© or Google Chrome©.

Participants navigated through tasks autonomously in a predetermined order among 15 possible pseudo-randomizations, one of which was automatically assigned by the system at the start of the testing sequence. Before each task, participants received written instructions in French on screen, and could not proceed until the natural reader finished delivering the same instructions orally. Fixed-length breaks (3 or 5 minutes) were included after the most intensive tasks to optimize concentration and compliance. Participants could end the break early if ready to continue, with a 60-second timer appearing in the last minute of the break if they had not yet proceeded to the next task.

#### Session 3: in-person behavioural testing

In this session, participants were tested in person for tasks that required closer supervision due to their length (e.g. the ANT-I), or that required manual measurements of reaction times (RT), (e.g. the literacy and literacy mediator tests). Testing was conducted at the Human Neuroscience Platform of “Campus Biotech” in Geneva, in a dedicated, sound-protected room, using the same laptop, mouse, and headset (headphone with microphone) for all participants (see Figure 3). Tasks were organized and delivered via the Gorilla interface using the previously described pseudo-randomization strategy. The same microphone check was included to ensure the safe recording of tasks requiring vocal responses for later assessment.

An experimenter closely supervised the session through a connected screen, mouse, and keyboard while facing the participant, intervening only when necessary due to task requirements or technical issues. For tasks requiring a vocal response, the experimenter manually recorded accuracy and/or RT (assessed via a chronometer) in the session booklet. To prevent data loss, these tasks were also audio-recorded, and the responses were later verified for accuracy by a team member with French as their first language. Task events not requiring manual measurement or verification were recorded directly in Gorilla. During this session, participants also performed three extra tasks that were part of another project^17^.

#### Session 4: Magnetic Resonance Imaging

Following behavioural testing, participants were invited to a brain imaging session on a Siemens 3T Magnetom-Prisma scanner equipped with a 64-channel head coil, again at Campus Biotech. Prior to scanning, they filled in and signed an MRI safety questionnaire to again verify their eligibility for the procedure. When required, participants were fitted with MRI-compatible goggles to correct their vision. The language localiser task was administered in Matlab r2021b with Psychtoolbox3, through a computer connected to a screen in the back of the scanner room, that participants could see through a mirror placed on top of the RF head coil. During the whole session, we could observe their eyes through an eye-tracking camera, to check that they were awake. During the imaging session, the field map, resting state sequence and the language localiser were administered in random order, but the resting state sequence was always preceded by the T1-weighted anatomical scan to avoid spurious activations due to carrying out an active task just before. After a short break outside the scanner, participants were repositioned and underwent the DWI sequence and its corresponding field map acquisition. During this session, participants also performed three fMRI tasks and one anatomical scan that were part of different projects. An overview of the imaging session parameters is provided in Table S1 of the Supplementary Information file.

#### fMRI language localiser

Participants were instructed to keep their eyes open and look at a black fixation cross on a white background while listening to intact and degraded snippets of the story ‘Alice in Wonderland’, in their first and second most dominant language of choice (L1, L2)^88,135^. This localiser can be used to inspect individual differences in language activation during quasi-naturalistic listening, and their relationship with behavioural measures of language and cognition^136^. The original localiser paradigm is <u>publicly available</u> from the authors’ webpage. As described in Table S1 in the Supplementary Information file, and the Data Records section, we modified this paradigm to include a degraded L2 condition.

#### Resting-state fMRI sequence

Brain connectivity at rest can be linked to individual differences in language^137,138^, reading^139,140^, and other domains of cognition such as executive function^141,142^. To collect information on resting-state brain activity, we asked participants to lie down with their eyes open, instructing them to relax their body and mind as best as they could, while projecting a white fixation cross on a black background.

#### BIDS conversion

This dataset conforms to BIDS v1.9.0 and was validated using the command line version of `bids-validator v1.14.14` (http://bids-standard.github.io/bids-validator/). A LINUX Terminal print of the output is provided in the Supplementary Information file (Figure S1). The dataset has been annotated using the Neurobagel annotation tool (https://neurobagel.org/) for enhanced findability^143^, the annotations have been saved in `/neurobagel` at root level, and all JSON sidecars have been validated with the online version of JSONLint. The BIDS data format conversion occurred in several steps, part of which could be planned before data collection (such as naming of the MRI sequences, folder structure, and participant codes), while others were performed *post hoc* to adapt data generated by environments not optimised for BIDS. In general, the procedure aimed at having a BIDS-coherent data structure and file names, (re)organising the content of tabular files, adding sidecar files to accompany data and customising code to work in a BIDS folder structure. These steps were performed on all data collected during the same testing session, and the NEBULA101 data were subsequently imported into the /nebula101 data space. Nonetheless, for clarity, we provide the specific heuristic for the construction of this dataset after DICOM to NiFTI conversion in `/code/heudiconv/heuristic.py`. Considering this procedure, all code described is specific to the NEBULA101 dataset but is not meant to be rerun, and is given with paths relative to the dataset BIDS root folder, but might reference to folders outside this structure, for example to source data. Outside of the code performing data import or behavioural data cleaning, no further reference is generally made to external/unavailable files. We describe the steps in Table 3.

**Table 3.**
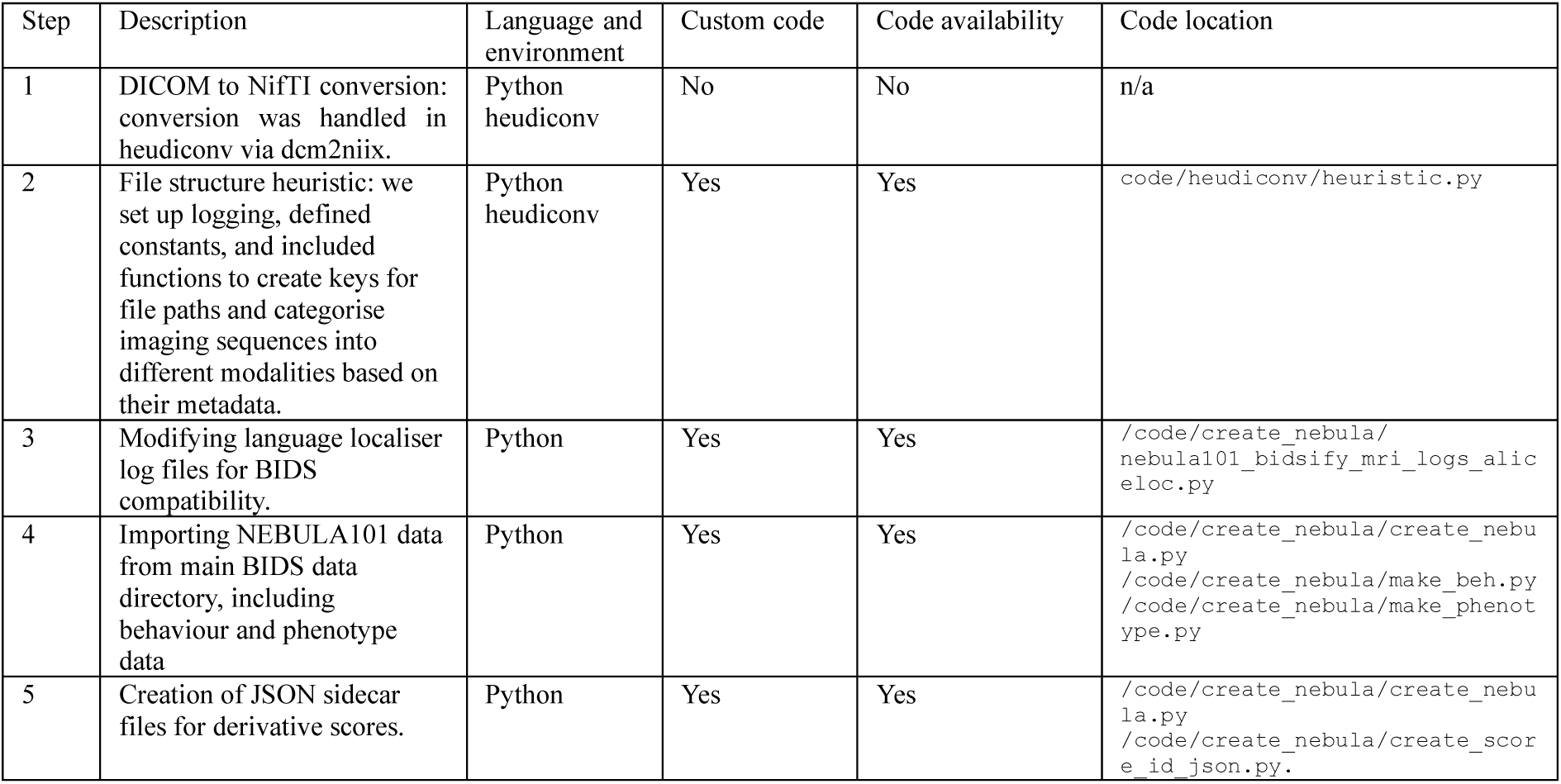
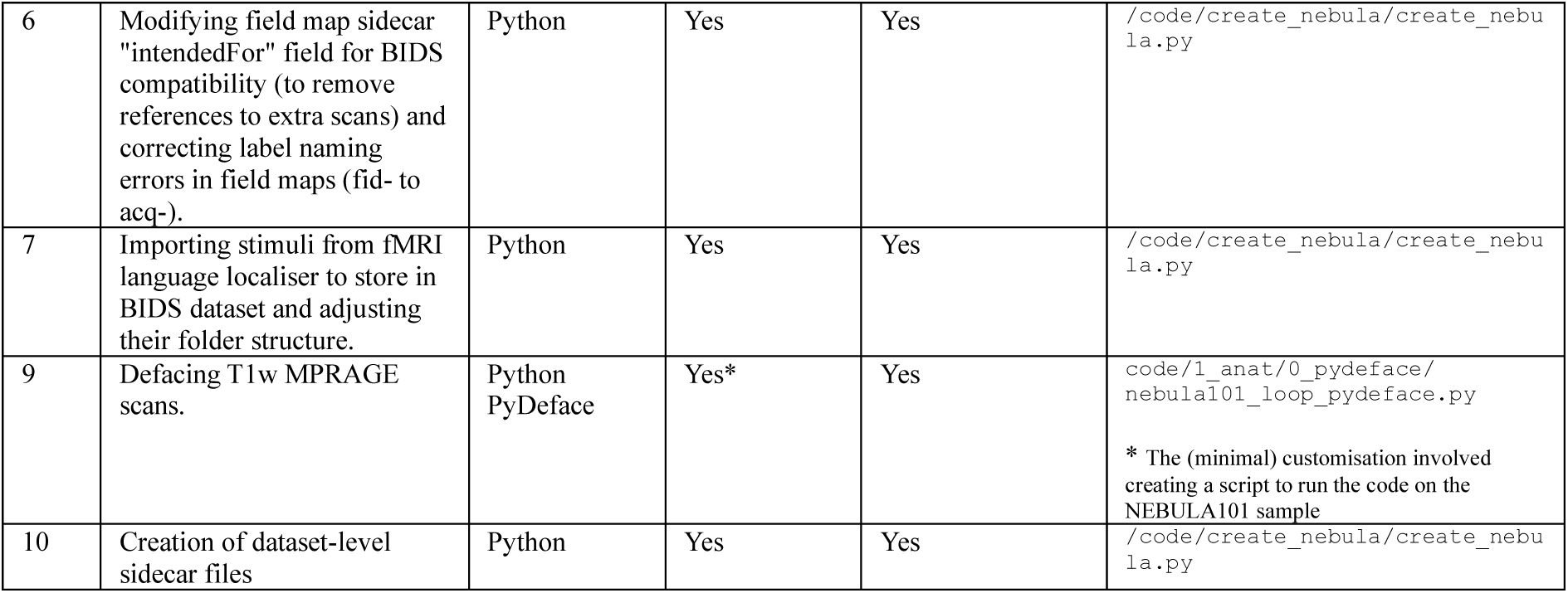
List of the steps taken to make the dataset BIDS-compliant.

### Data Records

The dataset is publicly accessible and downloadable from OpenNeuro under a Creative Commons CC0 1.0 (Universal Public Domain Dedication) license with accession number ds005613 at the address doi:10.18112/openneuro.ds005613.v1.0.1.

In this section we further describe the data structure and its contents. The folder `/nebula101` (Figure 4) constitutes the root level of the dataset. It contains 101 participant folders with raw imaging data denoted by the code `sub-pp` followed by three digits, the folders /`phenotype`, /`code`, /`derivatives`, /`stimuli, /neurobagel`, and the mandatory files required by BIDS (README, participant and dataset description files). The /`phenotype` folder contains tabular data from questionnaires, while `/neurobagel` contains subject-level annotations (harmonized phenotypic properties and imaging metadata) that can be encoded in a knowledge graph (see Technical Validation). As concerns the subject folders, inconsistent numbering is due to non-included participants (see Participants section), the participant who denied consent to share their data, and the two participants who could not undergo brain imaging). We describe the other contents of /`nebula101` in detail here below.

**Figure 4.**
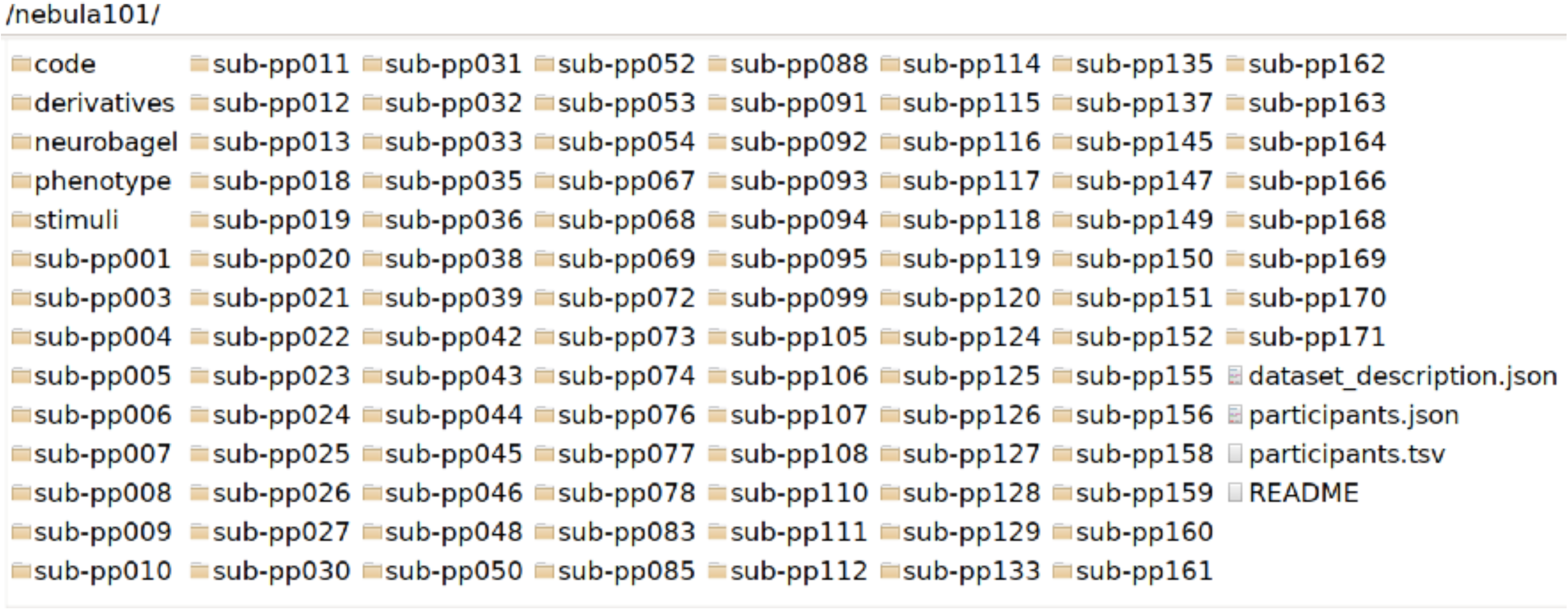
Root dataset folder.

The /code folder in Figure 5 contains subfolders with 1) fMRI paradigm data for the language localiser; 2) code to import, modify or generate the BIDS-compliant files as described in Table 3; 3) the BIDS conversion heuristic; 4) the preprocessing information and code folder; 5) the validation materials folder.

**Figure 5.**
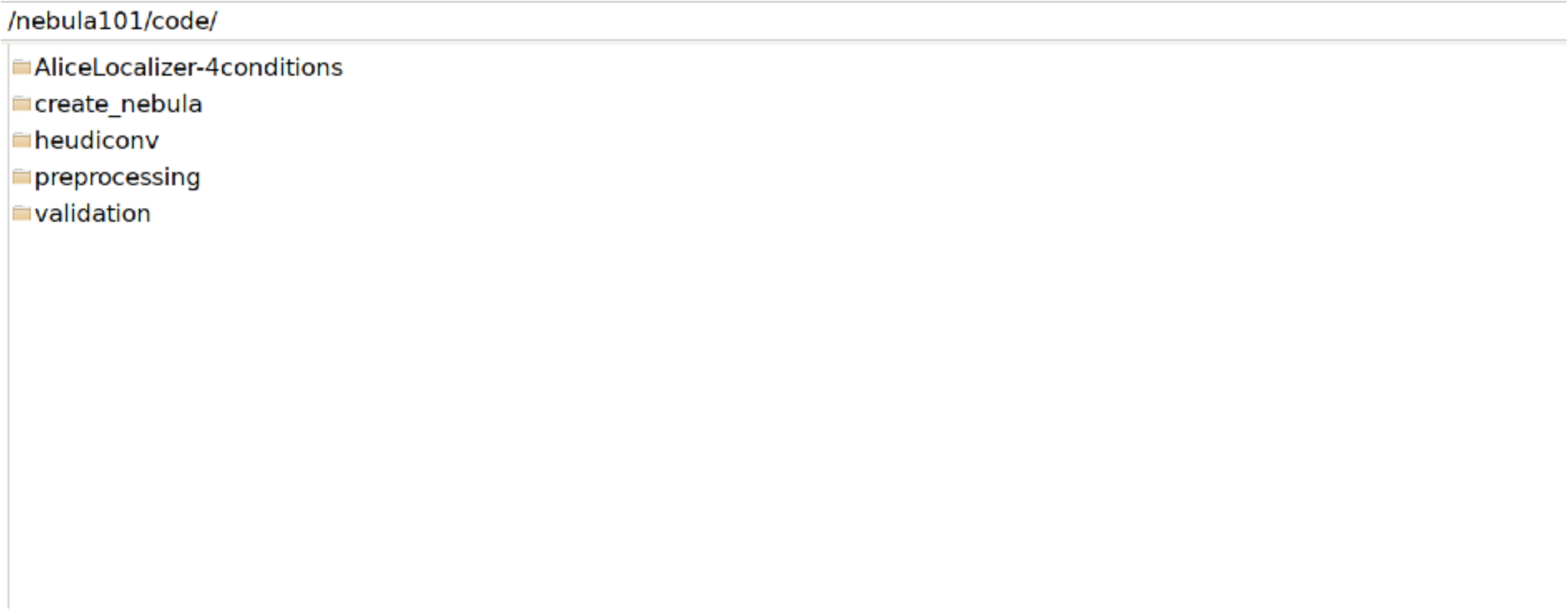
Code folder.

The /derivatives folder in Figure 6 contains the preprocessed behavioural data and their sidecar files, as well as the results of all validation pipelines.

- `cumulative_farsi_rater*` = derivative scores from the Farsi task ratings and their sidecar files.
- `nebula_101_all_questionnaire_scores*` = scores from questionnaires and their sidecar file.
- `nebula_101_all_task_scores*` = scores from tasks and their sidecar file.
- `nebula_101_leapq_annotation_iso_glottolog*` = mapping of language names to ISO an Glottolog and its sidecar file.
- `nebula_101_leapq_data*` = LEAPQ data and their sidecar file.
- `nebula_101_leapq_langname_order*` = LEAPQ languages in order of acquisition and their sidecar file.

**Figure 6.**
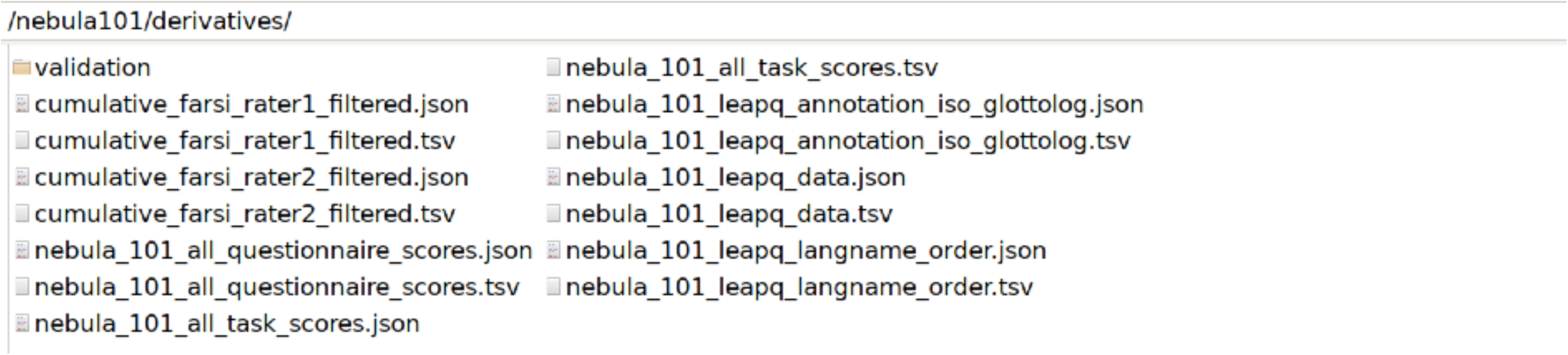
Derivatives folder.

The /stimuli folder contains a subfolder where the language localiser materials are stored and can be referenced by the BIDS validator.

The participant folders, named `sub-pp*/ses-01/` contain `/anat, /dwi, /fmap`, `/func` and `/beh` folders. These host the raw imaging and behavioural data of each participant, the latter having been minimally preprocessed to remove metadata and information unrelated to the scoring, as described in Technical Validation. An example with `sub-pp001/ses-01/` is shown in Figure 7, Figure 8, Figure 9, Figure 10 and Figure 11.

**Figure 7.**
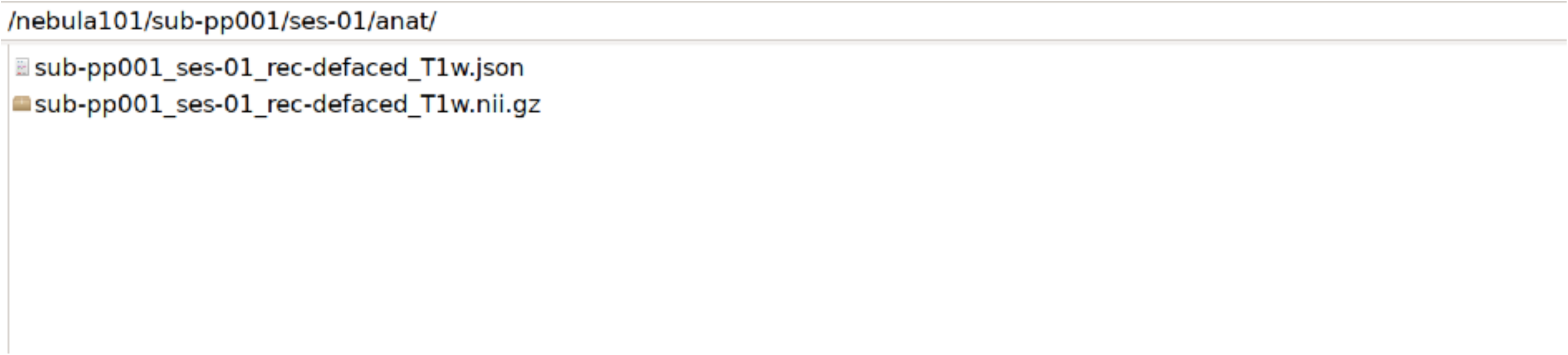
Defaced T1 MPRAGE data and their sidecar file.

**Figure 8.**
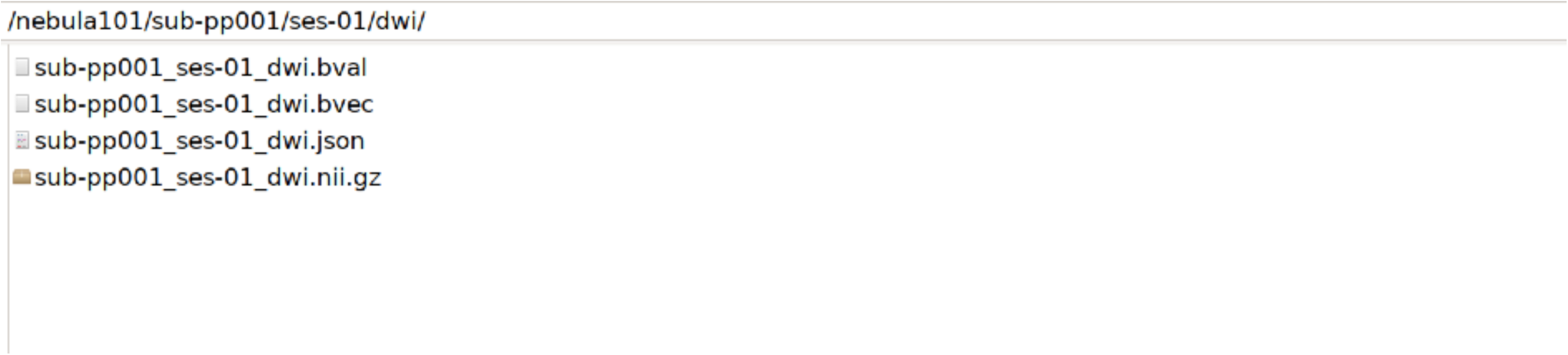
DWI data: beta values, vectors, and sequence data, with its sidecar file.

**Figure 9.**
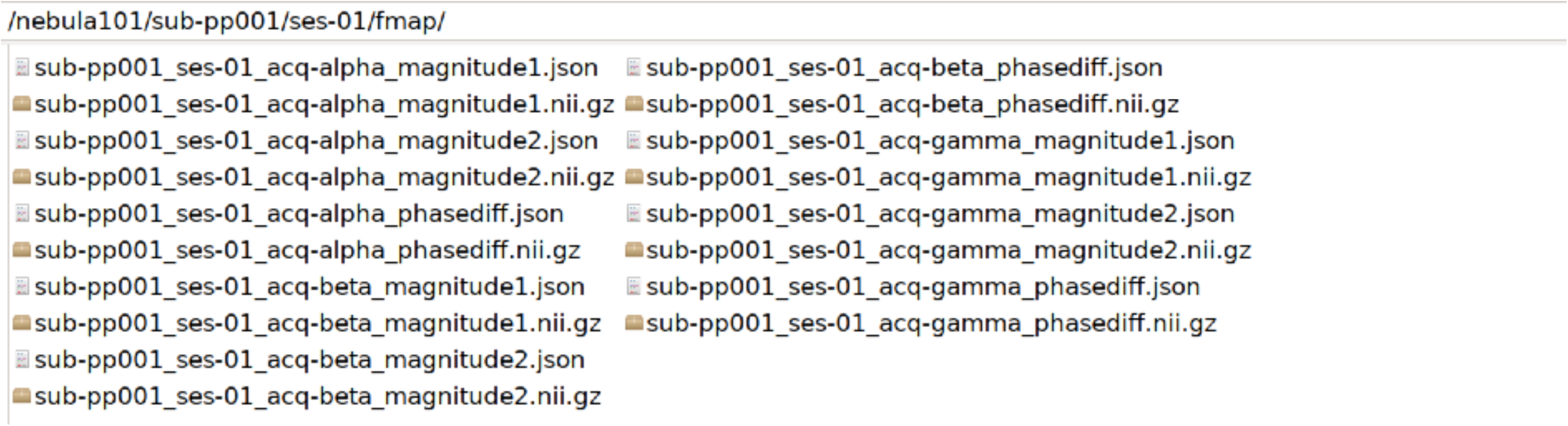
Field maps: phase and magnitude images with their sidecars. This participant has an extra field map (gamma), as explained in the Usage Notes.

**Figure 10.**
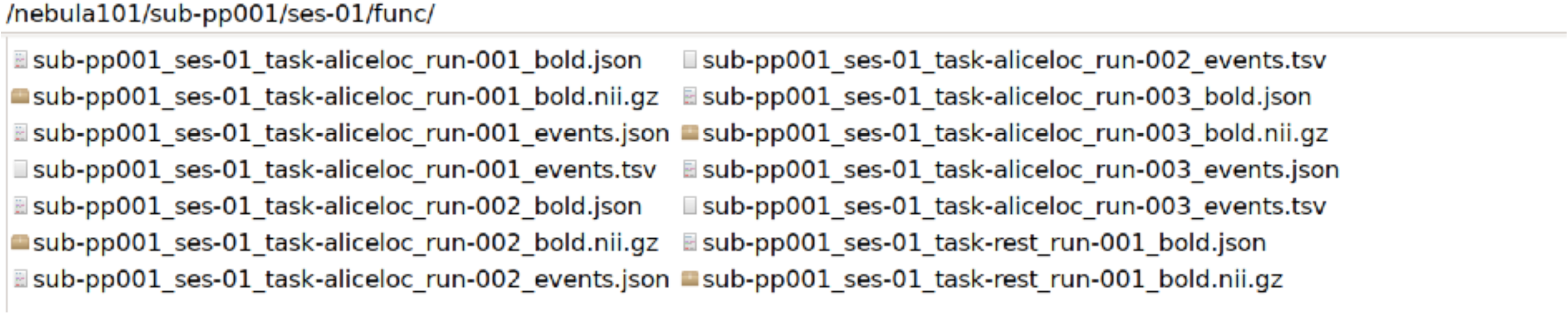
Functional MRI data with their sidecar files and event files. Rest refers to resting state fMRI, aliceloc refers to the language localiser.

**Figure 11.**
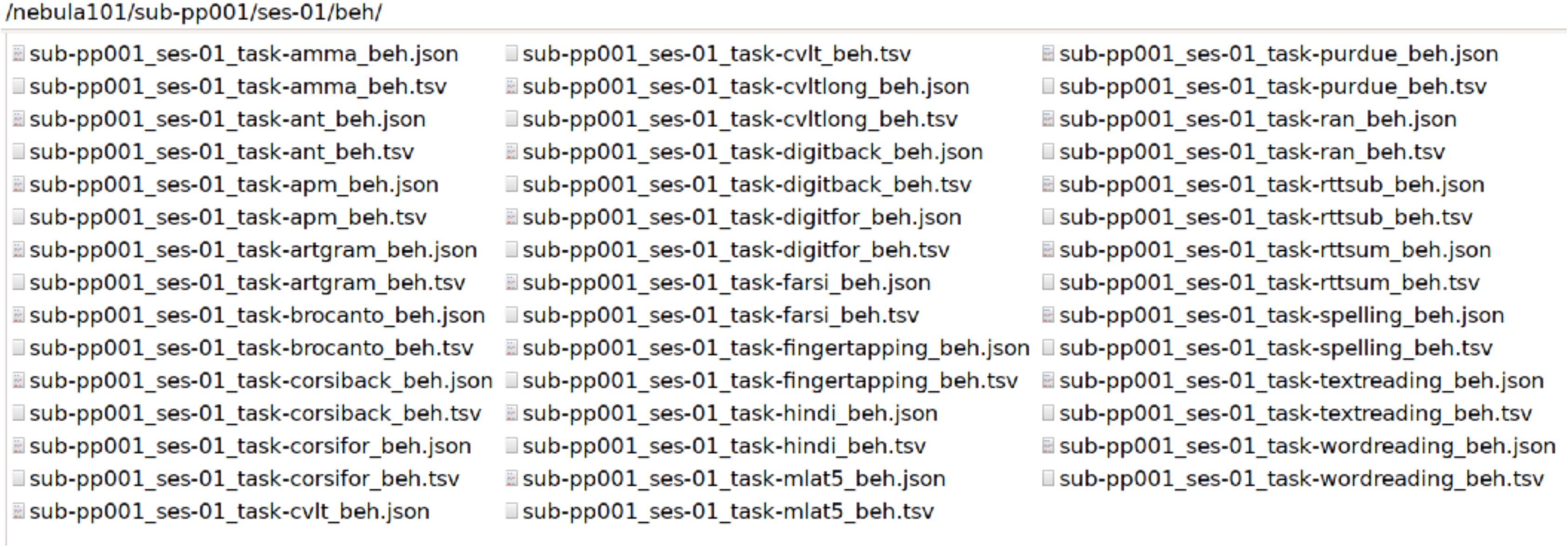
Raw behavioural data files with their sidecar files.

The `/code/AliceLocalizer-4conditions/` folder contains our own version of the materials provided in the public domain by the creators of the task^88,135^, which we modified to suit our needs, as described in the fMRI language localiser description (see: Data collection). This version of the code can read the stimuli from the structure of the BIDS folder. It will still require manual intervention to create the `events.tsv` files from the Matlab log files, as Matlab currently provides a different tabular structure. We remind the reader that we altered the structure of the log files to create simplified BIDS events with no redundant or unneeded information in `/code/create_nebula/bidsify_mri_logs_aliceloc.py`.

The `/code/preprocessing` folder includes numerically ordered subfolders containing code to reprocess the anatomical, field map, functional and diffusion data (see Technical Validation), and the reference code for the raw-to-derivative conversion of behavioural data. Any steps contained in these subfolders are meant to be run sequentially to recreate the materials needed for technical validation, or if the user decides to process our raw brain data using the same pipelines.

- `0_beh` contains behavioural data preprocessing code.
- `1_anat` contains the brain extraction code.
- `2_fmap` contains code for field map preparation.
- `3_func` contains code for minimal fMRI preprocessing, to get the displacement parameters used in the technical validation.
- `4_dwi` contains the preprocessing code for the diffusion data necessary to obtain technical validation reports.

The /code/validation folder contains the following subfolders (see Technical Validation for details on the operations performed):

- `/anat`: segmentation and sample homogeneity plotting code for the anatomical scans (auto-generated by Matlab): /`cat12/cat_stat_homogeneity.m` and `/cat12/cat_stat_homogeneity_autoplot.m`
- `/beh` contains the following items:

- The code to run reliability analysis on behavioural data: `calculate_cronbach_alpha.py`
- Three additional Python notebooks for generating the correlation data presented here and additional tables and matrices contained in the Supplementary Information file (Table S2, Table S3 and Figure S2):

- correlate_matrix.py
- calculate_descriptives.py
- farsi_inter_rater.py
- `/data_checks`: various Python scripts to create data lists of behavioural and imaging data, and missing data heatmaps, as explained in Technical Validation.
- `/dwi`: individual participant pdf QC reports, the code for generating them, and the group report folder.
- `/func`: two Python scripts, one for importing displacement `.rms` data from FSL, and the other for violin plots of average absolute and relative displacement during fMRI (resting state and task based).

- 1_copy_motion_params.py
- 2_mean_abs_rel_disp_violin.py

For each of the described folders in `/code/validation/`, there is a mirror /derivatives/validation/ folder where the results of the validation pipelines are stored, and specifically:

- `/derivatives/validation/anat/cat12/` contains the single subject CAT12 reports in PDF form.
- `/derivatives/validation/beh/` contains the item-level reliability data used to calculate Cronbach alpha and all correlation measures.
- `/derivatives/validation/data_checks/` contains data presence checks generated by the respective code for brain imaging, phenotype and behaviour.
- `/derivatives/validation/dwi/fsl/` contains the reports from subject- and group-level quality assessment of DWI data.
- `/derivatives/validation/func/fsl/aliceloc/` contains contains the reports from subject- and group-level quality assessment of task fMRI data.
- `/derivatives/validation/func/fsl/rest/` contains contains the reports from subject- and group-level quality assessment of resting state fMRI data.

All Python notebooks and executables are heavily commented for user-friendliness.

### Technical Validation

In compliance with the BIDS indications for mixed raw and derivative datasets, to improve user experience and reduce redundancy, we chose to include pre-processed behavioural data as derivative tables. We also provide quality control (QC) reports for all imaging modalities, together with the code for reproducing them. Below we specify the details of technical validation for each data type.

#### Questionnaires

Questionnaire source data were retrieved from Qualtrics XM© and preprocessed in Python. Each questionnaire was cleaned with its own script and then merged with the others via another script. The overall process aimed at data cleaning, as scoring was generally performed via the Qualtrics graphical user interface.

1. Log file data collection: for each questionnaire, from the source tabular file containing all participants, we eliminated irrelevant metadata. In rare cases, when duplicates were identified we eliminated them at source level by keeping the first non-empty instance of the questionnaire, blind to the contents, to avoid selection bias.
2. Data cleaning and handling of missing data: in all questionnaires, string responses were recoded as numbers. Missing data (NaN) was handled as follows: as a general rule, within-questionnaire, all responses were forced (i.e. progression was halted if a response was missing or on-screen warnings appeared to highlight unfilled responses). When information was indeed unavailable for a participant, we set up options to input specific strings, signalling to us that the information was missing. When missing information was nonetheless identified in a questionnaire with *optional* responses, but without the string signalling unavailability, we evaluated on a case-by-case basis whether it was possible to obtain it from the participant. In such rare cases, the questionnaire was retaken by the participant in session 4, assisted by an experimenter (to avoid data alteration), and only missing responses were filled in. In even rarer cases, when an *entirely* missing questionnaire was identified, participants were invited to complete it through an individual link to that specific questionnaire. Any remaining entries with unavailable information were converted to NaN during data preprocessing. With the strategies we put in place, all questionnaires have been completed by all participants in this dataset, except for one monolingual participant not having taken the code-switching questionnaire due to only speaking one language.
3. LEAPQ cleaning: the LEAPQ, due to its length and structure, required additional, *ad-hoc* handling. In Qualtrics, participants reported the number of languages they knew, including extinct ones and dialects, in order of dominance, that is, how much they used each of them over the others, and in order of acquisition, that is, the temporal order in which they learned their languages. They then provided general information for each language in order of dominance (e.g., names, exposure) and specific information (e.g., learning modality, use context). The resulting data file was restructured by language number and question type, filling gaps with NaN to create a ‘staircase-like’ structure where participants (rows) and questions (columns) are listed by language number, in ascending order. The resulting file is available in `/derivatives/nebula101_leapq_data.tsv` with a corresponding sidecar file explaining the column contents. The list of languages in order of acquisition was extracted into a separate file to avoid confusion, as it was not always the case that the dominance and acquisition lists coincided. Language order information can be accessed at /derivatives/nebula_101_leapq_langname_order.tsv.
4. Raw language names as entered by the participants were conformed to ISO 639-3 and Glottolog codes^144^ for standardised reference. The conversion map can be found in `/derivatives/nebula_101_leapq_annotation_iso_glottolog.tsv` and its sidecar file.
5. Multilingual language experience entropy. The LEAP-Q is a widely used instrument for assessing multilingual language experience, encompassing contexts of use, learning, use choices, history with the language, and native-likeness. Given that composite measures of multilingualism derived from the LEAP-Q would be beneficial for quantitative analyses, we calculated four different continuous ‘multilingualism scores’ for each participant, to reflect their multilingual experience cumulatively^64^. Three scores were based on their self-reported proficiency in speaking, reading, and comprehension across all reported languages, while the fourth score was based on current exposure to all reported languages. Following previous research^17,145,146^, each participant’s multilingualism score combined proficiency or exposure across their different languages using Shannon’s entropy equation^147^ within the R entropy package^148^.

Item-level questionnaire data are stored in `/nebula101/phenotype/` as individual tabular files, each containing data for one questionnaire, where each subject is represented in a single row as a `participant_id` and their responses in the subsequent columns. Each phenotypic tabular file is accompanied by a JSON sidecar file describing each column:

- `ahrq.tsv` and `ahrq.json` contain data for the Adult Reading History Questionnaire.
- `bsmss.tsv` and `bsmss.json` contain data for the Barratt Simplified Measure of Social Status.
- `code_swt.tsv` and `code_swt.json` contain data for the Code Switching Questionnaire.
- `handedness.tsv` and `handedness.json` contain data for the 10-item French version of the Edinburgh Handedness Inventory.
- `irq.tsv` and `irq.json` contain data for the Internal Reasoning Questionnaire.
- `mfq.tsv` and `mfq.json` contain data for the Motivation Factors Questionnaire.
- `musebaq.tsv` and `musebaq.json` contain data for the Music Experience Use and Engagement Questionnaire.

Given the higher level of processing that the LEAP-Q data went through, its tabular files (responses, language annotations and language order information) reside in /derivatives, together with the comprehensive derivate score file of all the questionnaires, nebula_101_all_questionnaire_scores.tsv.

### Behavioural tasks

Behavioural tasks were pre-processed in Python for data cleaning, derivate score calculation and BIDS conversion. The process happened in steps:

1. Log file collection: data were downloaded from Gorilla. Given that participants entered 1 out of 15 possible randomised task sequences, automatically assigned to people by Gorilla, the empty log files from the 14 unused randomisations for each participant had to be deleted. Microphone and headphone check files were identified and renamed.
2. Log file cleaning: irrelevant metadata were eliminated. Task log files were renamed to reflect the actual task (as Gorilla provides them in encrypted form) and to contain BIDS-compliant strings by adding the relevant labels, then converted to tab-separated values. Within each task’s log file, column names were renamed to more easily interpretable strings where necessary. Audio files from voice-recorded tasks were renamed as well, for easier reference.
3. Derivate score calculation and tabular file merging: for each task, one or more derivate scores were calculated, based on the task’s protocol. See Table 2. and Table S1 for the references and scores that were selected.

The behavioural data preprocessing code is provided for reference in /nebula101/code/preprocessing/0_beh, and specifically:

a. `2DataCleaning_gorilla.py`: this script cleans the source logfiles (not provided in this dataset) from metadata, as explained.
b. `3Scoring_gorilla.py`: this script scores the files that were generated by Gorilla and cleaned, based on an automated pipeline by using the ‘correct’ and ‘incorrect’ columns and/or reaction time information (depending on the task’s specific scoring protocol).
c. `4Scoring_manual.py`: this script scores logfiles that were created from tasks that required human intervention (for example, all tasks requiring manual RT measurements of voice responses via a chronometer, and/or live pencil scoring by the experimenter, such as the Text or Word and Pseudoword reading tasks, and/or the subject to write down answers like in the Spelling task). Here, Gorilla was used to run the task within the pseudo-randomisation pipeline but scoring necessitated manual intervention.
d. `5Scores-merging.py`: this script merged the scores into a tabular file, from which they were later imported to the `/nebula101` data space via the script `create_nebula.py`.
e. `7farsi_raters_assessment.py`: The script calculates cumulative and average scores for the Farsi uvular sound production task, based on assessments made by two independent raters. It produces the derivative file provided in `/nebula101/derivatives/`.

These actions were performed in subsequent steps by Python scripts acting on source data. Users will not be required to rerun these scripts since we provide cleaned tabular files containing the minimally processed raw accuracy and RT scores rather than the source data, as well as, crucially, the derivative scores calculated and ready for use.

For tasks whose data were recorded automatically in Gorilla (when a simple correct/incorrect input was sufficient, and RT could be measured by key press or mouse click), the raw data of each subject will contain their `participant_id`, as minimally mandated by BIDS, accuracy and RT columns for each trial, if these are sufficient to derivate a score provided in /`derivatives` via the code provided in `/code/preprocessing/0_beh/`. For the manually scored tasks, as described in steps c) and e) above, single subject raw data resulting from the digitisation of paper-and-pencil materials are provided: such might include block-level data (e.g. each iteration and type of recall trials in the case of the CVLT) or condition-level data (e.g. word types in the case of the Word and Pseudoword reading task, story types in the case of the Text reading task), whose structure will be inherent and specific to the task itself. In the case of the Spelling, Nonword repetition and Spoonerisms tests, scoring was digitised at task level (as per protocol). We therefore included these only in the derivative table. Despite any differences in structure, all behavioural files conform to the minimal BIDS requirements for their data type and are accompanied by extensive JSON descriptions. All materials used to process source data can be made available upon request, as well as source data that can be anonymised.

A comprehensive file containing all derivate scores for all behavioural tasks, obtained with steps b) and c) lives in /derivatives and is called nebula_101_all_task_scores.tsv.

In this location the Farsi task derivate scores obtained with step e) can also be found: these are called `cumulative_farsi_rater1_filtered.tsv` and `cumulative_farsi_rater1_filtered.tsv`. All derivate files are accompanied by JSON sidecars.

#### Behavioural data: correlations and reliability

Behavioural scores, whether resulting from tasks or questionnaires (here we generally refer to them together as “behavioural data”, and we use “scores” or “measures” when referring specifically to their outcomes), derive from item-level data. To validate these measures, we first explored them via correlations, and then ran reliability analyses in the form of internal consistency coefficients, in a conceptually similar approach to our recent exploratory analysis of behavioural data with a larger cohort^17^. Intercorrelation of the scores obtained from behavioural data (tasks and questionnaires) was minimal, and reliability was acceptable to good in most cases (please refer to the Supplementary Information file for com. The code for calculating Pearson’s correlations on the z-scored data as well as Cronbach Alpha, along with item-level reliability data and summary output (figures, tables) are available in `/code/validation/beh/` and in the Supplementary Information (Figures S2 and S3, Tables S2, S3, S4, S5).

To better understand the correlation patterns and reliability of these data, some considerations must be made about correlations and about reliability measured by internal consistency. Regarding correlations, on the one hand, high intercorrelations can be informative about shared variance between tests, but on the other hand, having an excessively intercorrelated dataset can lead to problems linked to multicollinearity^149–152^. On a conceptual level, the separability of latent variables facilitates the interpretation of the overall construct. Our choices, when planning this data collection and selecting the tests, aimed at isolating the best measures to represent each variable that we wanted to test in the context of this exploratory project on language aptitude. All things considered, a relative degree of independence of our measures was not only expected, but desirable. In the Supplementary Information (Table S2), we report all Pearson pairwise linear correlations having a coefficient of at least |*r*(100)| > .5 and significant at *p* < .05. In `/code/validation/beh/` code is provided to generate Table S2 and a matrix (Supplementary Figure S2) to visualise these data (`correlate_matrix.py`). Keeping in mind that correlations are just a preliminary way to explore data, most significant correlations were between metrics within the same test, and between these and their cumulative score, when present. As concerns correlations across tasks (marked with * and ** for positive and negative relationships respectively, in Table S2), the most evident patterns emerging are the correlations between reading measures and those typically related to or predictive of reading skill or deficit (all sub-scores from the RAN, text reading, word and pseudoword reading, spelling, spoonerisms, phoneme suppression and non-word repetition); between the Brocanto and CVLT long-term recognition score; between AMMA (total and tonal scores) and the musical training score from the MUSEBAQ (time, intensity and level of practice reached). No other measures were highly and significantly correlated.

Regarding reliability, the main advantage of internal consistency measured via Cronbach alpha is that, as long as covariance is inspected and there are minimal to no *nonignorable nonresponses* (MNAR data, i.e. missing not at random), it can be applied quite flexibly, unlike other methods^153^. Minimal reliability requirements are a matter of debate, as the α value can be affected by the number of items and improved by the independence of the construct these tap into^154–156^. This being said, it is harder to obtain a reliable measurement in tasks that measure *spans* (e.g. Corsi blocks, Digit span) and tasks where participants stop after reaching a maximum of answers with high variability (e.g. Revised tempo test): these will generate a MNAR pattern, where each row ends with a certain number of NaN because the span has been reached, and thus will be less suited to internal consistency measures, providing hardly interpretable coefficients^157^. This can explain the somewhat unreliable Revised tempo test RT, reflected in a pattern where negative or null covariances outweigh positive ones in a MNAR fashion (Figure 12), (`/code/validation/beh/cronbach_alpha_results.tsv`).

**Figure 12.**
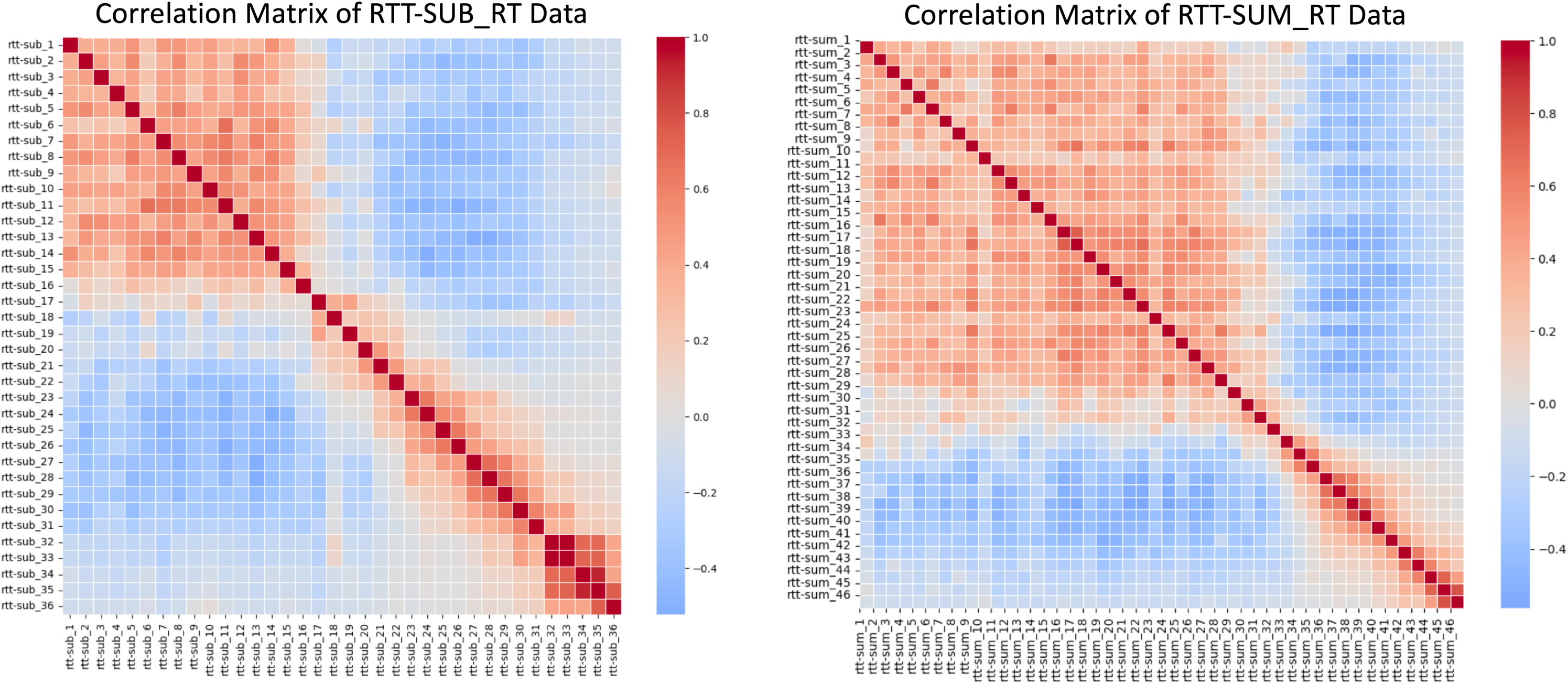
Revised tempo test correlation patterns for subtractions (left) and sums (right) RT data. The different number of items across the 2 figures is due to the fact that 1) participants reached a different number of completed operations across the two conditions and 2) no one reached the end of either condition (60 operations). Knowing that for this task, there is data missing for a reason, this can be interpreted as a MNAR pattern.

Rapid automatised naming accuracy (α = .21) as well as accuracy on the first run of Brocanto (α = .25) also had low reliability. We ascribe the former to the difficulty of the task for the participant, and therefore reaction times may provide more reliable measures than accuracy: RAN measures naming latency^158^, whose motor component may show more consistent RTs than accuracy (and this indeed shows in our data, as the two had α coefficients of .93 and .21, respectively). The lower reliability of the first run of Brocanto is conceptually interesting if compared to the subsequent runs and to the RT data: it shows that upon learning an artificial language completely inductively, responses are more variable at the beginning of the learning process, where people tend to guess more frequently, while the time to make a grammaticality judgment is overall always consistent. We provide the correlation matrices for accuracy and RT across the 3 blocks of Brocanto in Figure 13. In a few cases, data were recorded at the item level but digitised at the task level (spoonerisms, non-word repetition and phoneme suppression): item-level data are available upon request. To facilitate readers, in the Supplementary Information file we show the internal consistency values for tasks where α > .5, and tasks with α > .6 are additionally highlighted in bold (Table S3). Complete reliability data are available in `/code/validation/beh/` and can be regenerated with `calculate_cronbach_alpha.py`. Descriptive statistics and plots of the test metrics are available in the Supplementary Information file: Table S4, S5; Figure S4, S5, S6).

**Figure 13.**
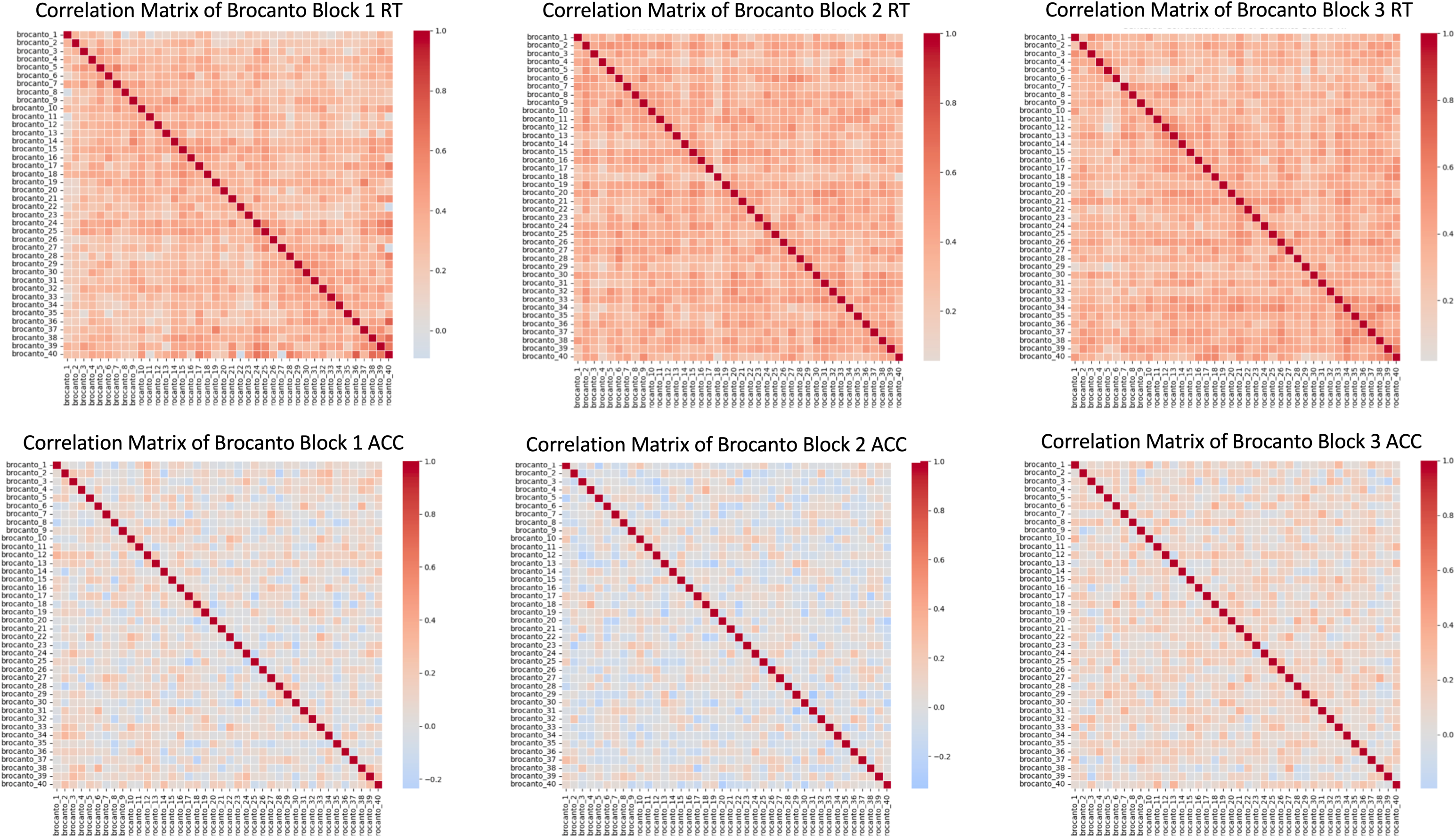
Top row: correlation matrix for the RT data of Brocanto across 3 blocks. Bottom row: accuracy data for Brocanto across 3 blocks. Plots show that overall, RTs were more reliable than was accuracy.

The Farsi uvular sound production task was rated by two first-language Farsi speakers who heard each recorded utterance from participants and gave it a ‘native-like production score’: therefore, any internal consistency metric would reflect the way the raters scored the task, more than the way the participant performed it (even though these are clearly related). Thus, for this task, we chose to calculate inter-rater reliability between the first and second rater, in a procedure identical to a previous study where this task was used^159^. For the ANT-I task, given its structure (each trial reflecting more than one possible condition from which a score can be derived), we chose to measure the maximal split-half coefficient to assess reliability^160^ (the Supplementary Information file contains the code for this procedure).

### Brain imaging

#### Anatomical imaging anonymisation, QC and brain extraction

Facial features in anatomical MRI scans violate the principle of anonymity in open data. Therefore, T1-weighted MPRAGE scans were defaced with PyDeface (https://pypi.org/project/pydeface/) and their quality was assessed in the Computational Anatomy Toolbox for SPM (hereon, CAT12)^161^. Specifically, we used the CAT12 segmentation and sample homogeneity toolboxes, providing easily interpretable quality measures at the participant and group level. The weighted overall image quality (IQR) and the quartic mean Z-score are the two key indicators of image quality. IQR combines noise and spatial resolution measurements before pre-processing, while the mean quartic Z-score assesses the homogeneity of data after pre-processing, with deviations increasing variance and reducing statistical power. The *product* of IQR and quartic mean Z-scores combines these quality measures, with a low number indicating high quality. For each participant, we provide a PDF with the CAT12 report, as well as information on sample homogeneity, in `/code/validation/anat/cat12/`. The CAT12 toolbox can be easily run via a graphical user interface (GUI) in Matlab and requires no custom code. We provide the group distribution of the described measures in Figure 14.

**Figure 14.**
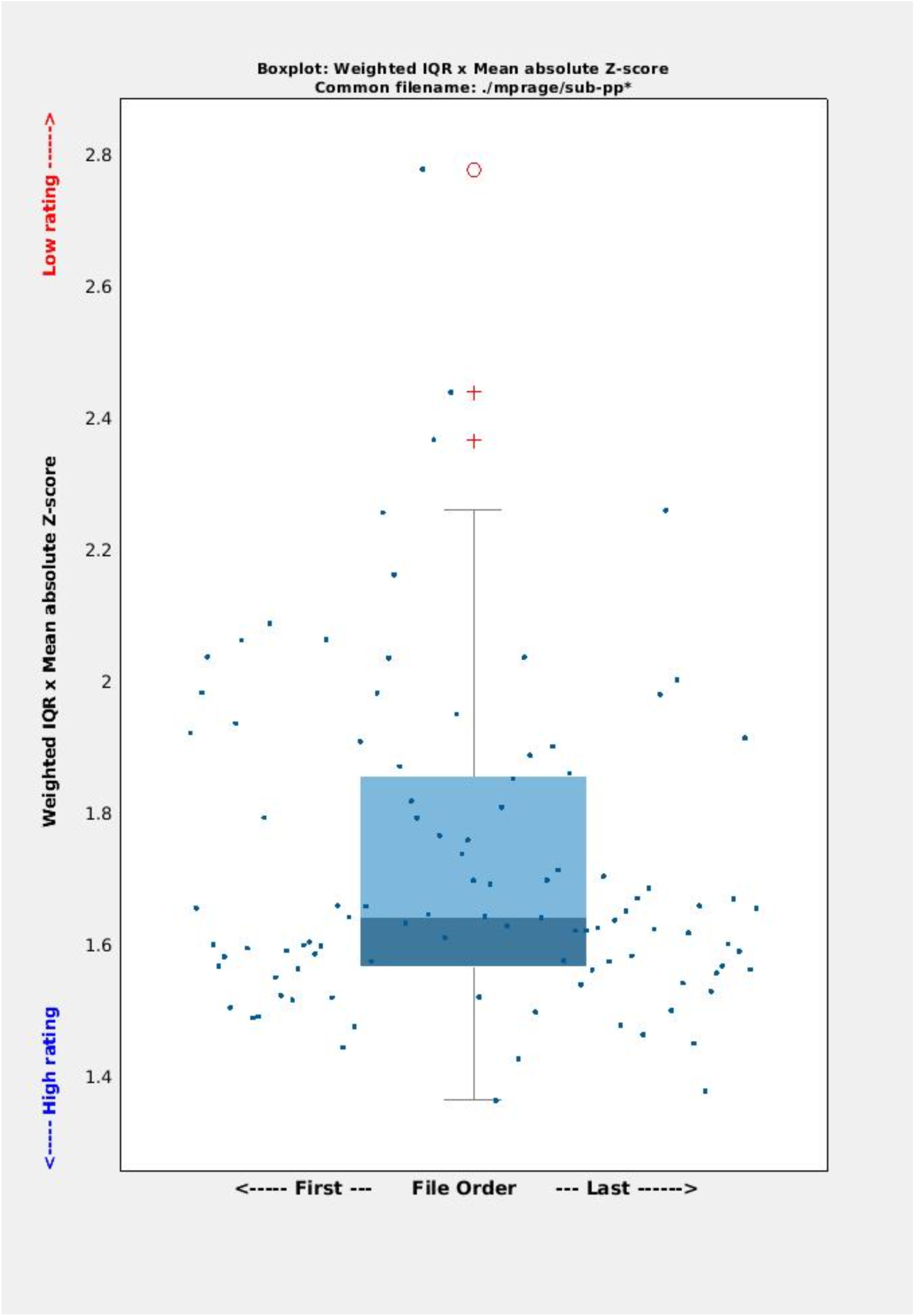
CAT12 output. Boxplot showing the weighted IQR by mean absolute z-score scaled by a factor of 4 (i.e. quartic) to emphasize outliers. As the plot shows, all but 1 participant in this distribution lie in the good to optimal range of the quality measure. Of note, even when outliers are detected, these are not necessarily data to discard on an absolute basis, as the score is calculated relatively to the specific sample being analysed.

To prepare T1 MPRAGE anatomical scans to be fed to FSL^162^ (see *Functional neuroimaging* section), we provide code for running an improved skull-stripping as a loop in the NEBULA101 dataset in `/code/preprocessing/1_anat/1_optiBET/nebula101_run_optiBET.sh.` This code will process T1-weighted MRI images stored in a BIDS directory to perform improved brain extraction with optiBET^136^ via the following actions:

1. Iterating through all participant directories in the BIDS directory.
2. For each participant, creating an output directory in the derivatives folder if it doesn’t exist.
3. Copying the T1-weighted images (`sub-pp*_ses-01_T1w.nii.gz`) to the corresponding output directory.
4. Checking if each T1-weighted image has already been processed.
5. If an image has not been processed, running the `nebula101_optiBET.sh` master script on the image, which performs actual brain extraction.

The script referenced in step 5 is the master optiBET script, which needs to be downloaded from https://montilab.psych.ucla.edu/fmri-wiki/optibet/ and stored in the location where the loop script is stored. *Field maps.* We acquired field maps to accompany the DWI and fMRI sequences. Field map preparation consists of a few steps that are partly scanner-specific, and in the case of Siemens Prisma, consisted in recomposing the phase and magnitude images. We provide field map preprocessing code in `/code/preprocessing/2_fmap/fsl/1_nebula101_run_fmap_prep_all.sh`

The script will perform the following steps:

1. Transformation Calculation: calculating the transformation matrix from the anatomical T1 image to the magnitude image and applying the calculated transformation to the T1 brain mask to align it with the magnitude image (`FLIRT`).
2. Magnitude Image Masking: using the transformed brain mask to mask the magnitude image, creating an untrimmed brain image (`fslmaths`).
3. Brain Mask Trimming (`fslmaths`):

a. Binarising the untrimmed mask.
b. Smoothing the mask with an 8 mm kernel.
c. Thresholding the smoothed mask at 75%.
d. Binarising the thresholded mask.
e. Trimming the original brain image using the final binary mask.
f. Removing intermediate untrimmed files.
4. Preparing the final field map (`fsl_prepare_fieldmap`): preparing the final field map using the processed magnitude and phase difference images, with a specified echo spacing of 2.46 ms.

#### Diffusion-weighted imaging

To validate our diffusion data, we run the `QUAD` (participant) and `SQUAD` (group) programs within the `FSL-FDT`^162,163^ suite as part of the diffusion pre-processing pipeline. The DWI data and prepared field map of each participant were fed to `EDDY` to correct for susceptibility, eddy currents, inter- and intra-volume participant movement and signal dropout, using the prepared field map via the `-- field` flag. Finally, we ran `eddyQC`^164^ for quality assessment. We provide a pdf with QC statistics for each NEBULA101 participant and for the group, as well as the code to reproduce this procedure, in `/code/validation/dwi/fsl/dwi_qc_quad_squad.py.` The code is configured to automatically grab the acquisition parameters and the correct number of shells from the DWI sidecars to generate the `acqparam.txt` and `index.txt` files required by `EDDY,` and will raise flags if the expected numbers do not match with the data. An `eddy_quad_qc_paths.txt` file has also been set up to this aim in the same location, which the user can alter for their needs, should they decide to preprocess DWI data (which we provide in raw form) with the same pipeline (`/code/preprocessing/dwi/fsl/1_dwi_prep.py`). Figure 15 reports the signal- and contrast-to-noise ratios from the group QC metrics generated using `SQUAD`. The complete PDF report is available in `code/validation/dwi/fsl/squad/group_qc.pdf.`

**Figure 15.**
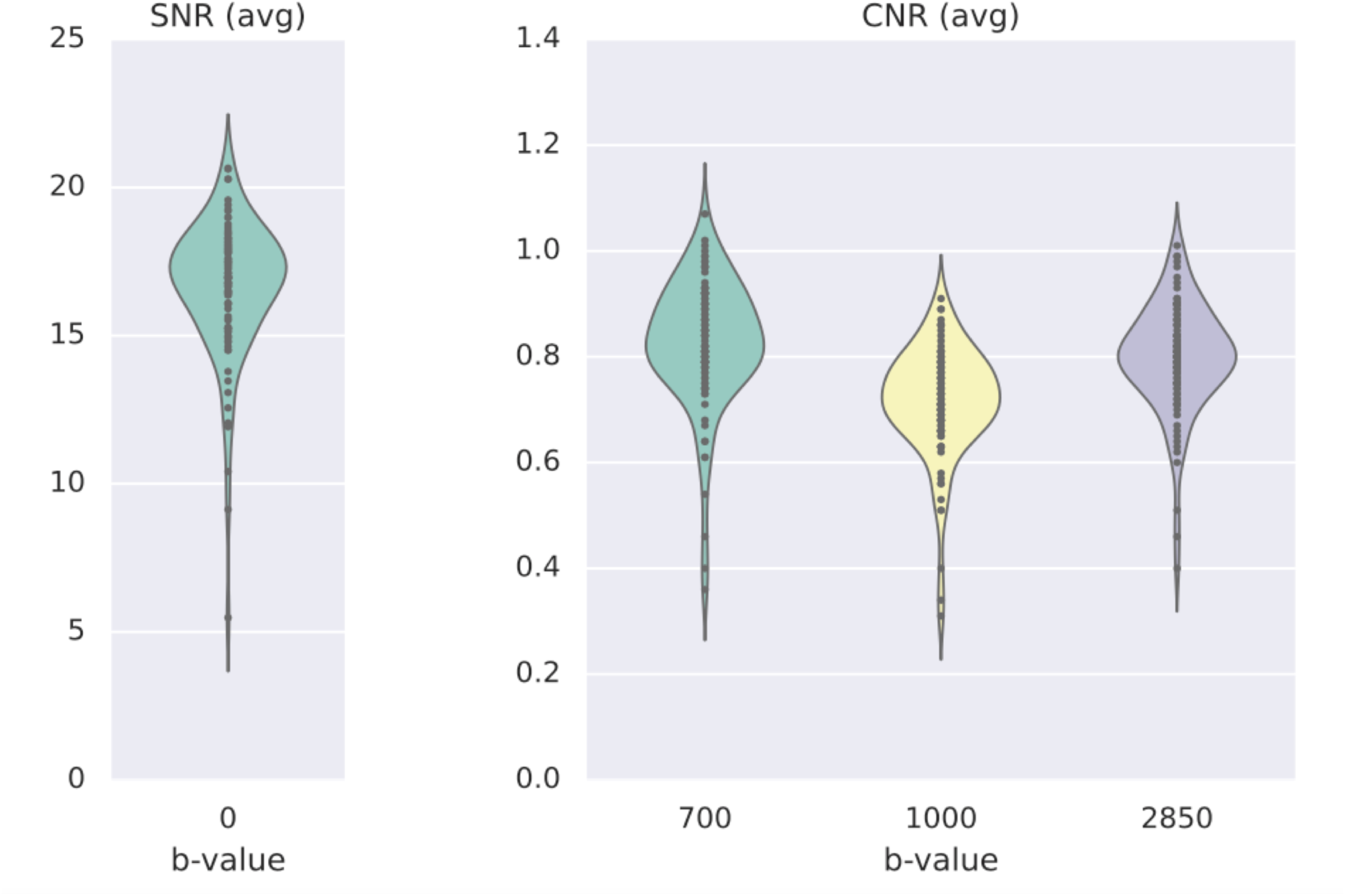
Signal-to-noise and contrast-to-noise ratios from the QC-ed DWI data.

#### Functional imaging

As concerns fMRI, we provide QC metrics obtained from preprocessing of the whole dataset in FSL, and the code to reproduce the procedure on the NEBULA101 sample.

Figure 16 and Figure 17 report violin plots of the absolute and relative displacement by run for the language localiser and for the resting state fMRI sequence, respectively. Absolute displacement reflects the movement with respect to the reference volume, while relative displacement reports volume-to-volume information.

**Figure 16.**
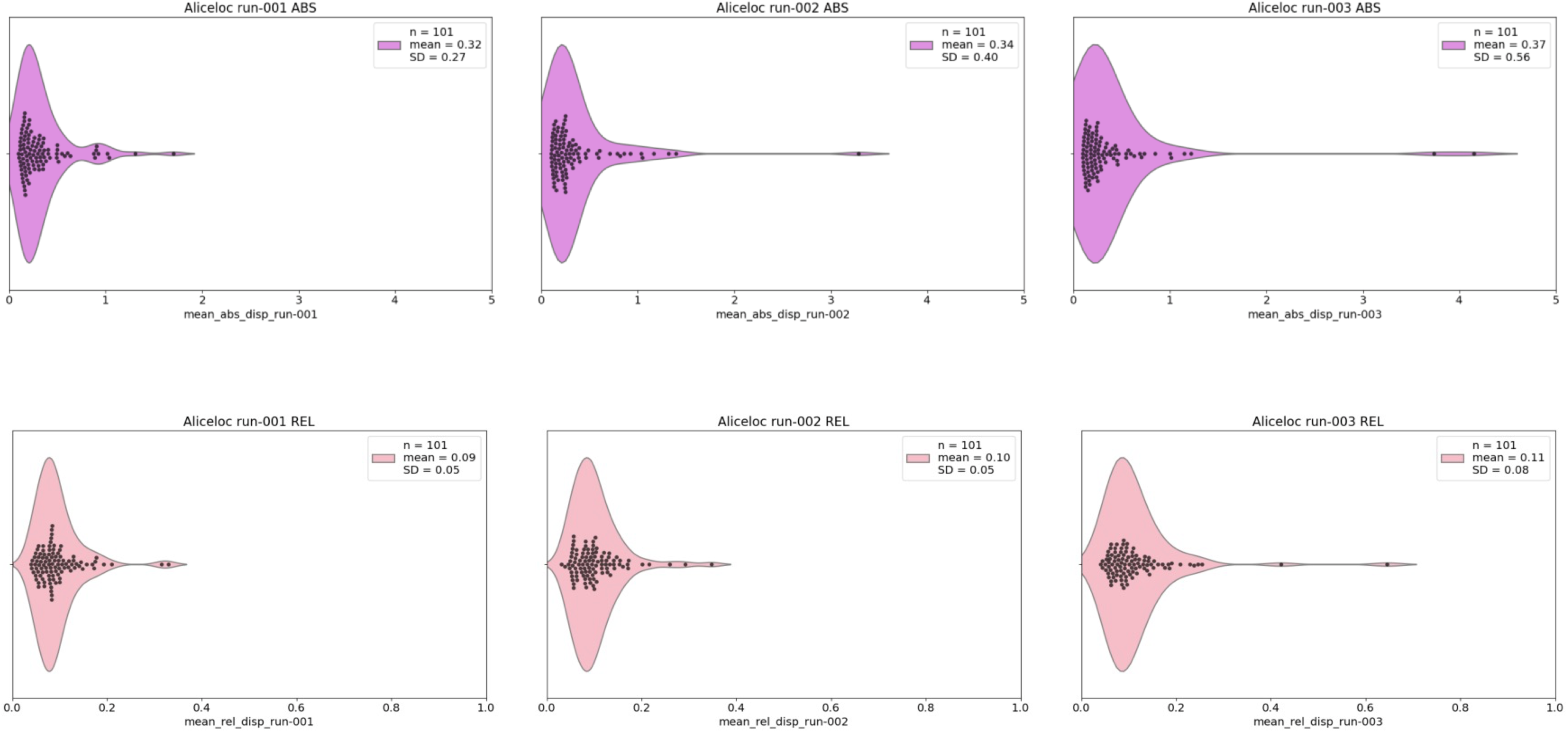
Absolute (ABS) and relative (REL) displacement (mm) across runs of the language localiser.

**Figure 17.**
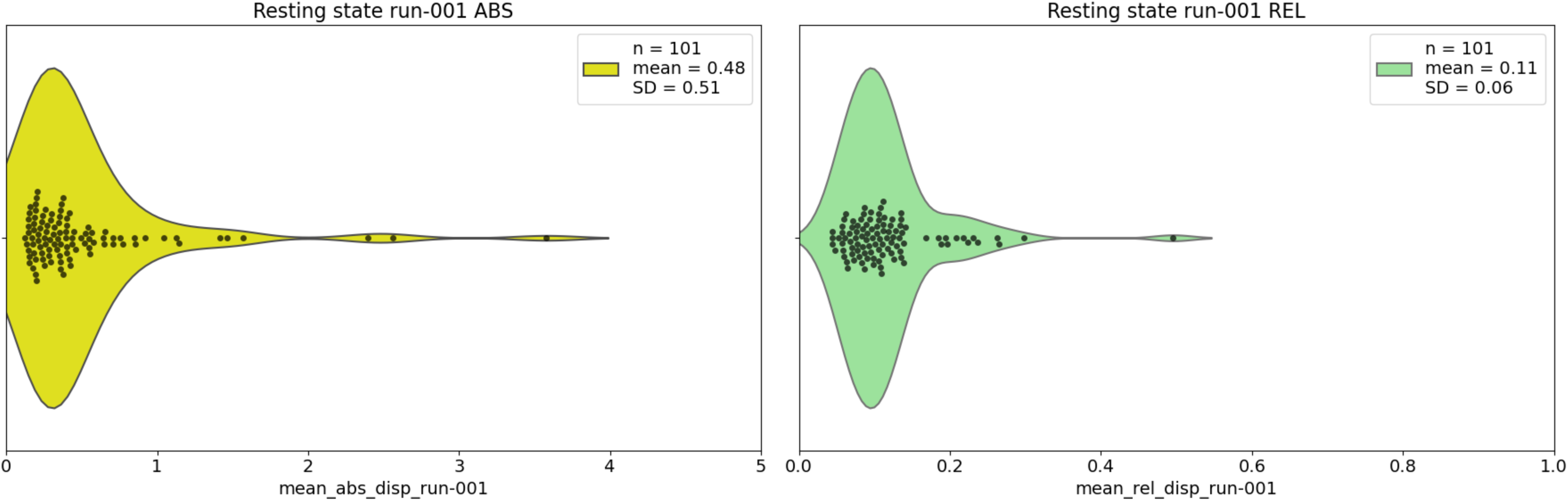
Absolute (ABS) and relative (REL) displacement (mm) during the single resting state sequence run.

Both can impact SNR but while absolute displacement can be counteracted, relative displacement is usually more harmful, and both must be checked when assessing the quality of fMRI data^165^. Displacement information and plots (Figure 16 and Figure 17) can be obtained by running `code/validation/func/fsl/1_copy_motion_params.py` and `/code/validation/func/fsl/2_mean_abs_rel_disp_violin_py.` In principle, if the data are not being reprocessed, `1_copy_motion_params` does not need to be rerun as we provide `.rms` output from FSL, containing absolute and relative displacement data for each participant, in `/code/validation/func/fsl/aliceloc/` and `/rest`. FIGURE 16 HERE

A folder containing the code for performing the basic preprocessing steps of fMRI for the language localiser and resting state sequences has been set up in `/code/preprocessing/3_func/fsl/1_preproc1/1_loop_fsl_preproc1.py`. This code performs the first step of fMRI preprocessing with FSL FEAT, consisting of:

1. Motion correction
2. 4D mean intensity normalization
3. Spatial smoothing (5 mm FWHM)
4. B0 unwarping (includes `BBR reg`)
5. Registration (`BBR, FNIRT`)

This code will run using /code/preprocessing/3_func/fsl/1_preproc1/design_pp.fsf, a design file required by FSL which has been set up to contain information on the preprocessing loop, should the user wish to preprocess our data with the same pipeline.

*Overall validation of data structure.* This dataset is mostly complete, with minimal missing information. However, to provide the user with information that is traceable without necessarily having this data descriptor at hand, we have created a series of scripts within the `/code/data_checks/` folder aimed at providing report logs on the presence of behavioural and brain imaging data. Such logs are saved to `/derivatives/validation/data_checks/`.

- `nebula101_bids_report_beh.py`: this script plots behavioural data presence checks on the raw and derivate dataset and compiles the findings to a report log called `nebula101_beh_report_[timestamp].txt`. It will also create:

- A tabular file `nebula101_datalist_[timestamp].tsv` listing data as 1 and 0 if present or absent, for easy reference.
- A heatmap with the above information, called `data_presence_heatmap_[timestamp].png`.
- A report specific to the LEAPQ data, to have missing and present information at a glance, called `nebula101_leapq_data_report_[timestamp].txt`
- `nebula101_bids_report_mri.py`: this script plots imaging data presence checks and compiles the findings to a report log called `nebula101_mri_report_[timestamp].txt`. This can be used for easy referencing to participants having undergone different conditions (e.g. different field map, see: Usage Notes).

### Usage Notes

#### Behavioural data

This dataset is mostly complete, but a few technical malfunctions during testing have caused some minimal missing data, as described in Table 4. This information and a heatmap for visualising missing data at a glance can easily be regenerated using the scripts in `/code/validation/data_checks/` called `nebula101_bids_report_beh.py` and `nebula101_missing_data_log.py.`

**Table 4.**
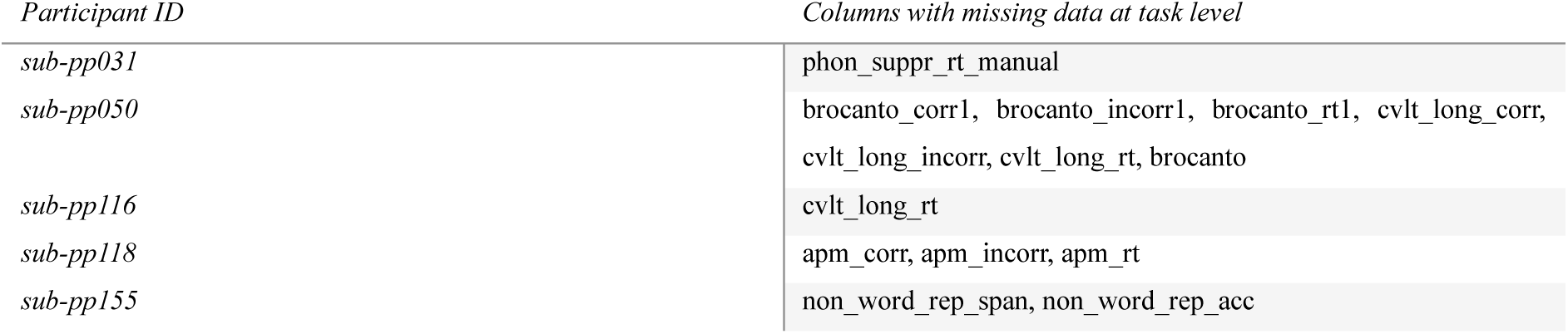
Missing behavioural data.

| Participant ID | Columns with missing data at task level |
| --- | --- |
| `sub-pp031` | `phon_suppr_rt_manual` |
| `sub-pp050` | `brocanto_corr1`, `brocanto_incorr1`, `brocanto_rt1`, `cvlt_long_corr`, `cvlt_long_incorr`, `cvlt_long_rt`, `brocanto` |
| `sub-pp116` | `cvlt_long_rt` |
| `sub-pp118` | `apm_corr`, `apm_incorr`, `apm_rt` |
| `sub-pp155` | `non_word_rep_span`, `non_word_rep_acc` |

#### Imaging data

Here we provide some details to facilitate easier navigation of the imaging data. This information can be generated by `/code/validation/data_checks/nebula101_bids_report_mri.py`. JSON sidecar files have already been adjusted to report the below nuances:

- `sub-pp161`: for this participant, the `beta` field map must be used for resting state fMRI processing, and the `delta` field map must be used for DWI processing.
- `sub-pp001, sub-pp010, sub-pp013, sub-pp023, sub-pp043, sub-pp053`: for these participants, the `gamma` field map must be used for resting state fMRI processing.
- `sub-pp032`: the voxel dimensions of `/sub-pp032/ses-01/anat/sub-pp032_ses- 01_rec-defaced_T1w.nii.gz` slightly differ in the y and z directions (pixdim1 = 1.000000, pixdim2 = 1.039060, pixdim3 = 1.039060).

When running the BIDS validator, this information is reported as *warnings* and saved in the log, unless the `--ignoreWarnings` flag is raised. Warnings to not impede BIDS validation and the dataset will pass it every time. When processing the data, this information will be read automatically by software that can work primarily in BIDS (such as fMRIprep^166^ in the case of functional imaging). When using software that lacks this functionality, such as FSL, we recommend processing these participants manually. The `.bidsignore` file contains instructions for skipping the validation of `/neurobagel` as it contains extra files.

## Code Availability

All code that can be rerun on these data is provided within the `/code` folder, as described in the Technical Validation section. Raw-to-derivative code is provided for all raw data. Given that we do not provide *source* behavioural data, source-to-raw preprocessing code for tasks and questionnaires is not included, but can be shared upon request.

## Author Contributions

CRediT (Contributor Roles Taxonomy) is provided, as follows: Conceptualization: AR IB OK RB NG; Investigation: AR IB; Methodology and Formal Analysis: AR IB OK; Data Curation: AR IB; Technical Validation: AR; Writing - Original Draft AR; Writing - Review & Editing: AR IB OK RB NG; Visualization AR; Resources: RB NG; Supervision: RB NG; Funding acquisition: RB NG.

AR and IB (*) contributed equally to this work.

## Supporting information

main supplementary material file

## Acknowledgments

We would like to express our sincere gratitude to several collaborators who contributed their valuable resources and support throughout this large and intensive project.

We gratefully acknowledge the following people: Roberto Martuzzi, Loan Mattera and Nathalie Philippe from the Human Neuroscience Platform at Campus Biotech for their invaluable and continued assistance with MRI setup and data collection; we thank our student assistants Vanessa Gottofrey, Melody Cascioli, and Aspasia Sfakaki for their contribution to data collection, and Rixa Gruhnert for helping with the LEAPQ annotations.

We are thankful to Michael Dayan for his help with setting up the BIDS heuristic; to Priscilla Borges, Jutta Mueller and Hettie Roebuck for discussions on the Modes of Internal Reasoning Questionnaire use and its scoring; to Mark Eckert and Davide Fedeli for their helpful hints on technical validation of anatomical and diffusion-weighted MRI; to Gwendoline Mahé for providing us with the French version of the Adult Reading History Questionnaire; to Peter Schneider for consultations on the musical experience and musicality tests; to Evelina Fedorenko and her team for discussing with us the modifications to the Alice in Wonderland Localiser.

We also thank Isabelle Udry for translating the Motivation Factor Questionnaire to French; Neiloufar Family and our colleague Sevil Maghsadagh for rating the Farsi task.

Finally, we thankfully acknowledge the collaborators who gave us access to and information on materials they have developed themselves: Tobias Kober and Siemens Healthineers© for the WIP-Compressed Sensing MPRAGE sequence, which crucially shortened MRI time; Will Barratt for sharing his Socioeconomic Status Measure; Franck Ramus and Steve Majerus for sharing their literacy and phonological awareness tasks; Charles Stansfield for granting permission to use the MLAT5; Elsje Van Bergen for providing the Tempo Test Revised.

Lastly, our heartfelt gratitude goes to two anonymous reviewers whose suggestions greatly improved this dataset, and to the 101 participants who took part in our study, for their commitment and generosity with their time.

