## Supplementary material for "NEBULA101: an open dataset for the study of language aptitude in behaviour, brain structure and function": main supplementary material file

### Supplementary Information File

**Supplementary information v2.0 OCT-24: NEBULA101: an open dataset for the study of language aptitude in behaviour, brain structure and function**

A. Rampinini, I. Balboni, O. Kepinska, R. Berthele, N. Golestani

This file contains supplementary information for the paper **NEBULA101: an open dataset for the study of language aptitude in behaviour, brain structure and function**

For details, please refer to the main article.

Figure S1. Output of BIDS validation (screen-print from the LINUX Terminal):

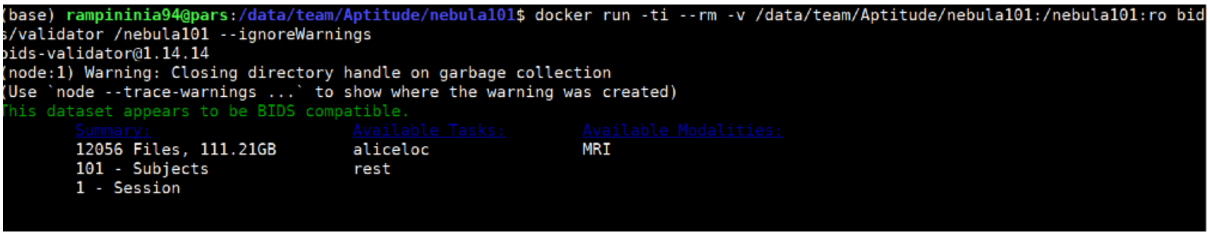

A. Rampinini, I. Balboni, O. Kepinska, R. Berthele, N. Golestani

Table S1: overview of all tests and modalities, with details specific to this dataset. For the complete bibliography, please refer to the main article, Table 2.

| *Test* | *Modality* | *Construct* | *Delivery* | *Instructions* | *Derived score* | *Specifications* |
|---|---|---|---|---|---|---|
| ***Language Experience and Proficiency Questionnaire (LEAP-Q)*** | Questionnaire | Multilingual language experience. | Online, unsupervised | Fill in the provided form. | Entropy | Extended to accommodate up to 50 languages.<br>Added questions on:<br>*Time spent* in contexts such as online communities, fandoms and subcultures where a language is spoken.<br>*Contributors* to language learning: social media, apps and everyday life in the country where a language is spoken.<br>Removed question on cultural identification.<br>French adaptation. |
| ***Code Switching questionnaire*** | Questionnaire | Code switching habits. | Online, unsupervised | Fill in the provided form. | Voluntary switching<br>Involuntary switching | Selected only voluntary and involuntary switching scores.<br>French adaptation. |
| ***Motivational Factors Questionnaire (MFQ)*** | Questionnaire | Motivation and attitude towards foreign languages (FL). | Online, unsupervised | Fill in the provided form. | Ideal foreign language self<br>Instrumentality<br>International contact<br>Foreign language interest<br>Foreign language anxiety<br>Foreign language confidence<br>Milieu<br>Usage willingness of foreign languages | French adaptation.<br>5-point Likert scale instead of 6-point.<br>Constructs/items assuming that participants are actively involved in studying foreign languages (academically) were not included.<br>Items that refer specifically to the English language, anglophone world and culture reworded to fit a more global context where possible or excluded.<br>Included constructs are shown in the Derived Score column. |
| ***Adult Reading History Questionnaire (AHRQ)*** | Questionnaire | Reading history. | Online, unsupervised | Fill in the provided form. | Reading history score | French version of the questionnaire. |

| | | | | | | |
|---|---|---|---|---|---|---|
| ***Internal Representations Questionnaire (IRQ)*** | Questionnaire | Modes of internal reasoning. | Online, unsupervised | Fill in the provided form. | Manipulation Factor<br>Orthographic Factor<br>Verbal Factor<br>Visual Factor | French adaptation. |
| ***Music Use and Background Questionnaire (MUSEBAQ)*** | Questionnaire | Music training, capacity, preferences, and motivations. | Online, unsupervised | Fill in the provided form. | Index of music training<br>Index of music listening<br>Index of musical instrument playing<br>Cognitive and Emotional Regulation<br>Social Connection<br>Engaged Production<br>Dance<br>Physical Exercise | French adaptation. |
| ***Barratt's Simplified Measure of Socioeconomic Status (BSMSS)*** | Questionnaire | Socioeconomic status. | Online, unsupervised | Fill in the provided form. | Barratt's Simplified Measure of Socioeconomic Status | French adaptation. |
| ***Edinburgh Handedness Inventory (EHI)*** | Questionnaire | Handedness. | Online, unsupervised | Fill in the provided form. | Handedness score | 10-item version. |
| ***Artgram*** | Behavioural task | Language analytic abilities / Morphosyntax. | Online, supervised | Study the provided dictionary and choose the appropriate adaptation for given sentences by recognising the use of morphological cues. | Accuracy<br>RT | 7-word dictionary to study for 3 minutes.<br>12 sentences to translate, with 4 possible choices, only 1 correct choice.<br>Self-advancement.<br>15-minute time limit. |
| ***Modern Language Aptitude Test 5 (MLAT5)*** | Behavioural task | Rote learning. | Online, supervised | Study the provided dictionary and choose the appropriate adaptation for given words. | Accuracy<br>RT | French adaptation. |
| ***Farsi uvular Production Task*** | Behavioural task | Foreign sound production. | Online, supervised | Listen to and reproduce words containing a foreign | Accuracy | n/a |

| | | | | | | |
|---|---|---|---|---|---|---|
| | | | | language sound not present in French. | | |
| ***Hindi Dental Retroflex Contrast*** | Behavioural task | Phonological categorisation/discrimination. | Online, supervised | Categorise a foreign language sound not present in French. Training and testing blocks. | Accuracy | 200 trials were presented. |
| ***Brocanto*** | Behavioural task | Language analytic abilities / Pattern recognition. | Online, supervised | Recognise grammatical and ungrammatical sentences with different structures and violation types in an artificial language, inductively.<br>Training (reading-only) and testing (judge grammaticality by button press) blocks. | Accuracy<br>RT<br><br>(For each block 1-3 and overall) | Version from Kepinska et al., 2017 with minor modifications related to timing and condition counterbalancing, as follows:<br>A practice screen was added at the beginning of each testing block to test button press and rule comprehension.<br>Button press instructions: 1 for "grammatical" and 0 for "ungrammatical".<br>We chose a 2-second fixation cross without jittering.<br>We chose a 6-second sentence presentation in both testing and training block types.<br>We maintained 40 sentences per testing block, but we included 8-word sentences and obtained a 1/2 ratio of grammatical and ungrammatical sentences **overall** (N=120, of which the 60 ungrammatical sentences were new and roughly 1/3 of the 60 grammatical ones [N=22] had been presented during training). |
| ***Raven's Advanced Progressive Matrices (APM)*** | Behavioural task | Non-verbal intelligence. | Online, supervised | Select the missing block to complete a picture from a set of proposed choices. | Accuracy<br>RT | Advanced and abridged French version.<br>Time limit of 20 minutes for 23 trials.<br>Programmed for computer-based presentation on the Gorilla platform. |
| ***Corsi block*** | Behavioural task | Visuospatial memory | Online, supervised | Watch a sequence of blocks being highlighted, and repeat the same sequence by clicking on them in the same order and then backwards. | Forward span<br>Backward span<br>Total span | Programmed for computer-based presentation on the Gorilla platform in place of in-person.<br>The blocks were presented as 9 2-dimensional black squares arranged on a white background.<br>Blocks were black.<br>Sample block sequences were shown in |

| | | | | | | |
|---|---|---|---|---|---|---|
| | | | | | | yellow.<br>Participants clicked on the blocks instead of tapping them with their hand.<br>Selected blocks were shown in green upon clicking.<br>Block placement on the screen followed Kessels et al. 2000[89] |
| ***Digit Span*** | Behavioural task | Auditory working memory | Online, supervised | Recollect the digits immediately after the presentation in the same order, in the 'forward' section of the task, or in the inverse order, in the 'backward' section. | Forward span<br>Backward span<br>Total span | Auditory modality.<br>Response collected through upper numeric keypad. |
| ***Revised Tempo Test*** | Behavioural task | Arithmetic abilities | Online, supervised | Solve 60 additions and 60 subtractions in 1 minute each. | Accuracy<br>RT<br><br>(Additions, subtractions, overall) | Programmed for computer-based presentation on the Gorilla platform in place of paper and pencil, otherwise unchanged.<br>Response collected through upper numeric keypad. |
| ***Advanced Measures of Music Audiation (AMMA)*** | Behavioural task | Music audiation, musicality, musical aptitude. | Online, supervised | Judge difference or identity in melody or rhythm between pairs of musical excerpts. | Tonal score<br>Rhythm score<br>Total score | Programmed for computer-based presentation on the Gorilla platform. |
| ***Attention Network Test-Interaction (ANT-I)*** | Behavioural task | Attention networks: executive control, alerting, orienting. | In person | Detect arrow orientation in presence of flankers and sound cues. | Alerting gain<br>Orienting gain<br>Reorienting gain<br>Inhibition gain | French adaptation (instructions)<br>18 conditions and 432 trials. |
| ***California Verbal Learning Task (CVLT)*** | Behavioural task | Verbal working memory | In person | Remember lists of words short-term and long-term.<br>Recognise previously presented words, long-term. | Accuracy immediate recall<br>Accuracy immediate and short-term recall<br>Accuracy long-term recognition<br>RT long term recognition | Programmed for computer-based presentation on the Gorilla platform.<br>Removed cued recall (short and long-term). |

| | | | | | | |
|---|---|---|---|---|---|---|
| ***Finger tapping Test*** | Behavioural task | Fine motor skills | In person | Tap with the index finger of the dominant and then non-dominant hand on the spacebar as fast as possible. | Average finger tapping score dominant hand overall<br>Average finger tapping score nondominant hand overall<br>Average finger tapping score of dominant/nondominant overall<br><br>(Calculated over blocks) | Abbreviated version of the Finger Tapping test as proposed by Ashendorf et al., 2015.<br><br>5 trials per hand.<br>3 dominant hand trials – 1-minute break – 2 dominant hand trials.<br>3 non-dominant hand trials – 1-minute break – 2 non-dominant hand trials. |
| ***Purdue Pegboard Test*** | Behavioural task | Fine motor skills | In person | Insert pegs in holes, first with the dominant and and then with the non-dominant hand.<br><br>Build assembly of pegs alternating hands as instructed. | Accuracy, dominant hand<br>Accuracy, nondominant hand<br>Accuracy, simultaneous hands<br>Accuracy, assembly task<br>Average accuracy, dominant-nondominant-simultaneous hands<br><br>(Calculated over blocks) | French adaptation (instructions) |
| ***Rapid Automatised Naming (RAN)*** | Behavioural task | Naming automatisation | In person | Rapidly denominating objects, digits, colours | Accuracy<br>RT | French adaptation.<br>Programmed for computer-based presentation on the Gorilla platform. |
| ***Phoneme suppression*** | Behavioural task | Phonological awareness | In person | Repeat words by omitting the first phoneme. | Accuracy<br>RT | Programmed for computer-based presentation on the Gorilla platform.<br><br>Pre-recorded female voice. |
| ***Text Reading*** | Behavioural task | Reading skills | In person | Read two texts of increasing difficulty. | Accuracy<br>RT | Programmed for computer-based presentation on the Gorilla platform. |
| ***Word and Pseudoword Reading*** | Behavioural task | Reading skills | In person | Reading lists of words and pseudowords. | Accuracy<br>RT<br><br>(Per stimulus type: regular words, irregular words and pseudowords; overall) | With the target population in mind, this task was made more difficult by merging two standardized dyslexia assessment tests in French, the ECLA16+ and the EVALEC:<br>- There were 56 regular words, 52 irregular words, and 56 pseudowords overall. |

| | | | | | | |
|---|---|---|---|---|---|---|
| | | | | | | - For each category 20 words were taken from the Ecla16+ and the remaining from the EVALEC.<br>- The words from the EVALEC were randomized within-list rather than mixing across lists, to be able to time and score the lists per word-type (regular, irregular and pseudowords). |
| ***Spelling task*** | Behavioural task | Spelling skills | In person | Write down words, pseudowords and irregular words after hearing them. | Accuracy<br>(regular words, irregular words and pseudowords; overall) | Programmed for computer-based presentation on the Gorilla platform. |
| ***Spoonerisms*** | Behavioural task | Phonological awareness | In person | Swap the first phoneme of word pairs. | Accuracy<br>RT | Programmed for computer-based presentation on the Gorilla platform.<br>Pre-recorded male voice |
| ***Non-word repetition*** | Behavioural task | Phonological working memory | In person | Repeat non-word lists of increasing length. | Accuracy of repetition<br>Span, i.e. maximum number of repeated words | Programmed for computer-based presentation on the Gorilla platform. |
| ***Magnetization Prepared - RApid Gradient Echo (MPRAGE) imaging*** | sMRI | Brain structural anatomy | In person | Lie still in scanner. | n/a | whole-brain coverage<br>1mm isotropic voxel<br>FOV read = 256mm, FOV phase 93.8%<br>TR = 2300ms, TE = 3.26ms<br>Flip Angle: 9°<br>Distance Factor 50 %<br>Orientation: Sagittal<br>Phase Encoding Direction: A >> P |
| ***Diffusion-weighted Imaging (DWI)*** | dMRI | Fiber tractography | In person | Lie still in scanner. | n/a | Multishell sequence:<br>1.5mm isotropic voxel<br>FOV read = 225mm, FOV phase 100%<br>whole-brain coverage<br>TR 6700.0 ms, TE 74.00 ms<br>Distance Factor 0 %<br>Acceleration factor SMS = 2 GRAPPA = 3<br>Orientation: Transversal<br>Phase Encoding Direction: A >> P<br>Diffusion-encoding gradient directions: 117 |

| | | | | | | |
|---|---|---|---|---|---|---|
| | | | | | | $12\ B_0$ volumes distributed along the sequence<br>7 volumes at 700 s/mm2<br>30 volumes at 1000 s/mm2<br>68 volumes at 2850 s/mm2 |
| ***Language network brain functional localiser*** | fMRI | Functional activation for language | In person | Lie still in scanner, eyes open, fixate cross, listen to story | n/a | Added a condition for degraded second language.<br>Added a fixation at the beginning and at the end of each run. |
| ***Resting-state brain function*** | fMRI | Resting-state functional activation | In person | Lie still in scanner, eyes open | n/a | 2mm isotropic voxel<br>FOV read = 224mm, FOV phase 100%<br>whole-brain coverage<br>72 slices<br>TR = 2000ms, TE = 32ms<br>Flip Angle: 75°<br>Distance Factor 0 %<br>Acceleration factor SMS = 3<br>Orientation: Transversal<br>Phase Encoding Direction: A >> P |
| ***Field map*** | MRI | Intensity of the magnetic field across space | In person | Lie still in scanner. | n/a | Intended to correct B0 distortion in fMRI:<br>$2.4x2.4x2mm^3$ voxel<br>FOV read = 225mm, FOV phase 100%<br>whole-brain coverage<br>72 slices (fMRI), 66 slices (DWI)<br>TR = 700ms, TE1 = 4.92ms, TE2=7.38ms<br>Flip Angle: 60°<br>Distance Factor 0 %<br>Orientation: Transversal<br>Phase Encoding Direction: R >> L |

A. Rampinini, I. Balboni, O. Kepinska, R. Berthele, N. Golestani

Table S2. Pairwise linear Pearson correlations among variables that reached significance and were above |r(100)| > .5. For brevity, between scores having a 'corr' and 'incorr' (correct ,incorrect) columns, only 'corr' was selected. Age and education were not considered. Correlations across tasks are marked with * and ** for direct and inverse, respectively.

| Construct(s) | Test(s) | Variable 1 | Variable 2 | Correlation ($r_{Pearson}$) |
|---|---|---|---|---|
| Verbal working memory: immediate and total recall | CVLT | cvlt_tot_imm | cvlt_tot_recall | 0.99 |
| Music audiation, musicality, musical aptitude: tonal accuracy and total accuracy | AMMA | amma_tonal | amma_total | 0.99 |
| Music audiation, musicality, musical aptitude: rhythm accuracy and total accuracy | AMMA | amma_rhythm | amma_total | 0.99 |
| Reading skills: regular words and total accuracy | Word and Pseudoword Reading | regular_acc | wordreading | 0.97 |
| Arithmetic abilities: subtractions accuracy and total accuracy | RTT | rtt_sub_corr | arith | 0.96 |
| Fine motor skills: dominant hand taps and total accuracy | Finger Tapping Test | finger_tapping_dominant | fingertap | 0.96 |
| Music audiation, musicality, musical aptitude: rhythm accuracy and tonal accuracy | AMMA | amma_tonal | amma_rhythm | 0.96 |
| Reading skills: pseudowords and total accuracy | Word and Pseudoword Reading | pseudo_acc | wordreading | 0.95 |
| Fine motor skills: non-dominant hand taps and total accuracy | Finger Tapping Test | finger_tapping_nondominant | fingertap | 0.95 |
| Arithmetic abilities: additions accuracy and total accuracy | RTT | rtt_sum_corr | arith | 0.95 |
| Fine motor skills: both hands accuracy and accuracy across all single-tool trials | Purdue Pegboard Test | purdue_both_avg | purdue_dh_ndh_both_avg | 0.91 |
| Reading skills: irregular words and total accuracy | Word and Pseudoword Reading | irregular_acc | wordreading | 0.91 |
| Fine motor skills: non-dominant hand and accuracy across all single-tool trials | Purdue Pegboard Test | purdue_ndh_avg | purdue_dh_ndh_both_avg | 0.9 |
| Reading skills: regular and irregular words RT | Word and Pseudoword Reading | regular_rt | irregular_rt | 0.9 |
| Reading skills: regular and pseudowords accuracy | Word and Pseudoword Reading | regular_acc | pseudo_acc | 0.89 |
| Fine motor skills: dominant hand and accuracy across all single-tool trials | Purdue Pegboard Test | purdue_dh_avg | purdue_dh_ndh_both_avg | 0.88 |
| Reading skills: regular and irregular words accuracy | Word and Pseudoword Reading | regular_acc | irregular_acc | 0.88 |
| Arithmetic abilities: additions and subtractions RT | RTT | rtt_sub_rt | rtt_sum_rt | 0.88 |
| Auditory/verbal working memory: backward span and total span | Digit span | digit_back_span | span_verbal | 0.87 |
| Reading skills: regular and pseudowords RT | Word and Pseudoword Reading | regular_rt | pseudo_rt | 0.85 |

| | | | | |
|---|---|---|---|---|
| Language analytic abilities / Pattern recognition: block 2 and block 3 RT | Brocanto | brocanto_rt2 | brocanto_rt3 | 0.85 |
| Spelling skills: irregular words and total accuracy | Spelling Task | spelling_irregular_acc | spelling_tot_acc | 0.83 |
| Fine motor skills: dominant and non-dominant hand taps | Finger Tapping Test | finger_tapping_dominant | finger_tapping_nondominant | 0.83 |
| Arithmetic abilities: additions and subtractions accuracy | RTT | rtt_sub_corr | rtt_sum_corr | 0.82 |
| Auditory/verbal working memory: forward span and total span | Digit span | digit_for_span | span_verbal | 0.82 |
| Fine motor skills: non-dominant hand and both hands accuracy | Purdue Pegboard Test | purdue_ndh_avg | purdue_both_avg | 0.78 |
| Visuospatial memory: forward span and total span | Corsi Block | corsi_for_span | span_visual | 0.78 |
| Reading skills: pseudowords and irregular RT | Word and Pseudoword Reading | irregular_rt | pseudo_rt | 0.77 |
| Fine motor skills: accuracy across all single-tool trials and assembly | Purdue Pegboard Test | purdue_dh_ndh_both_avg | purdue_assembly_avg | 0.76 |
| Visuospatial memory: backward span and total span | Corsi Block | corsi_back_span | span_visual | 0.76 |
| Fine motor skills: both hands accuracy and assembly accuracy | Purdue Pegboard Test | purdue_both_avg | purdue_assembly_avg | 0.75 |
| Reading skills: pseudowords and irregular accuracy | Word and Pseudoword Reading | irregular_acc | pseudo_acc | 0.75 |
| Language analytic abilities / Pattern recognition: block 2 and overall accuracy | Brocanto | brocanto_corr2 | brocanto | 0.74 |
| Language analytic abilities / Pattern recognition: block 3 and overall accuracy | Brocanto | brocanto_corr3 | brocanto | 0.74 |
| Spelling skills: regular words and total accuracy | Spelling Task | spelling_regular_acc | spelling_tot_acc | 0.73 |
| Language analytic abilities / Pattern recognition: block 1 and overall accuracy | Brocanto | brocanto_corr1 | brocanto | 0.73 |
| Reading skills: test reading and regular word reading RT* | Text reading & Word and Pseudoword Reading | reading_text_rt | regular_rt | 0.72 |
| Spelling skills: pseudowords and total accuracy | Spelling Task | spelling_pseudo_acc | spelling_tot_acc | 0.72 |
| Language analytic abilities / Pattern recognition: block 1 and block 2 RT | Brocanto | brocanto_rt1 | brocanto_rt2 | 0.72 |
| Phonological working memory: span and accuracy | Non-word repetition | non_word_rep_span | non_word_rep_acc | 0.71 |
| Fine motor skills: dominant hand and both hands accuracy | Purdue Pegboard Test | purdue_dh_avg | purdue_both_avg | 0.7 |
| Reading skills: test reading and irregular word reading RT* | Text reading & Word and Pseudoword Reading | reading_text_rt | irregular_rt | 0.69 |
| Reading skills: test reading and pseudoword reading RT* | Text reading & Word and Pseudoword Reading | reading_text_rt | pseudo_rt | 0.68 |
| Fine motor skills: dominant hand and assembly accuracy | Purdue Pegboard Test | purdue_dh_avg | purdue_assembly_avg | 0.67 |

| | | | | |
|---|---|---|---|---|
| Naming automatisation and Reading skills: total RT and regular word reading RT* | Rapid Automatised Naming & Word and Pseudoword reading | ran_tot_rt | regular_rt | 0.67 |
| Fine motor skills: dominant hand and non-dominant hand accuracy | Purdue Pegboard Test | purdue_dh_avg | purdue_ndh_avg | 0.66 |
| Language analytic abilities / Pattern recognition: block 1 and block 3 RT | Brocanto | brocanto_rt1 | brocanto_rt3 | 0.66 |
| Naming automatisation and Reading skills: automatic naming total accuracy and irregular words RT* | Rapid Automatised Naming & Word and Pseudoword reading | ran_tot_acc | irregular_acc | 0.64 |
| Language analytic abilities / Pattern recognition: block 2 and block 3 accuracy | Brocanto | brocanto_corr2 | brocanto_corr3 | 0.64 |
| Naming automatisation and Reading skills: automatic naming total RT and irregular words RT* | Rapid Automatised Naming & Word and Pseudoword reading | ran_tot_rt | irregular_rt | 0.63 |
| Fine motor skills: non-dominant hand and assembly accuracy | Purdue Pegboard Test | purdue_ndh_avg | purdue_assembly_avg | 0.62 |
| Naming automatisation and Reading skills: automatic naming total and irregular words accuracy* | Rapid Automatised Naming & Word and Pseudoword reading | ran_tot_acc | regular_acc | 0.62 |
| Motivation and attitude towards foreign languages: ideal L2 self and instrumentality scores | MFQ | ideal_l2_self | instrumentality | 0.62 |
| Naming automatisation and Reading skills: automatic naming and word reading total accuracies* | Rapid Automatised Naming & Word and Pseudoword reading | ran_tot_acc | wordreading | 0.59 |
| Naming automatisation and Reading skills: automatic naming and text reading total RT* | Rapid Automatised Naming & Text reading | ran_tot_rt | reading_text_rt | 0.59 |
| Naming automatisation and Reading skills: automatic naming and pseudowords RT* | Rapid Automatised Naming & Word and Pseudoword reading | ran_tot_rt | pseudo_rt | 0.59 |
| Motivation and attitude towards foreign languages: ideal L2 self and international contact scores | MFQ | ideal_l2_self | intl_contact | 0.57 |
| Phonological awareness RT * | Spoonerisms & Phoneme suppression | spoon_rt_manual | phon_suppr_rt_manual | 0.56 |
| Motivation and attitude towards foreign languages: ideal L2 self and interest in foreign languages scores | MFQ | ideal_l2_self | l2_interest | 0.56 |
| Spelling skills: regular and irregular words accuracy | Spelling Task | spelling_regular_acc | spelling_irregular_acc | 0.55 |
| Code switching habits: contextual and involuntary switching | Code Switching | swt_score_cs | swt_score_us | 0.55 |
| Naming automatisation and Reading skills: automatic naming and nonword repetition accuracy* | Rapid Automatised Naming & Nonword Repetition | ran_tot_acc | non_word_rep_acc | 0.52 |
| Language analytic abilities / Pattern recognition block 1 accuracy & Verbal working memory (long term recall)* | Brocanto & CVLT | brocanto_corr1 | cvlt_long_corr | 0.52 |
| Language analytic abilities / Pattern recognition total accuracy & Verbal working memory (long term recall)* | Brocanto & CVLT | cvlt_long_corr | brocanto | 0.51 |
| Music audiation, musicality, musical aptitude and Musical experience: tonal accuracy and musical training* | AMMA & MUSEBAQ | amma_tonal | ind_mus_train_imt | 0.5 |
| Music audiation, musicality, musical aptitude and Musical experience: total accuracy and musical training* | AMMA & MUSEBAQ | amma_total | ind_mus_train_imt | 0.5 |

| | | | | |
|---|---|---|---|---|
| Reading skills: test reading accuracy and irregular word reading RT** | Text reading & Word and Pseudoword Reading | reading_text_acc | irregular_rt | -0.51 |
| Reading skills: test reading RT and Spelling skills total accuracy** | Text reading & Spelling test | reading_text_rt | spelling_tot_acc | -0.51 |
| Reading skills: test reading RT and Spelling skills irregular words accuracy** | Text reading & Spelling test | reading_text_rt | spelling_irregular_acc | -0.52 |
| Rote learning: accuracy and RT | MLAT5 | mlat5_corr | mlat5_rt | -0.55 |
| Reading skills: test reading accuracy and regular word reading RT** | Text reading & Word and Pseudoword Reading | reading_text_acc | regular_rt | -0.59 |
| Reading skills: test reading accuracy and pseudoword reading RT** | Text reading & Word and Pseudoword Reading | reading_text_acc | pseudo_rt | -0.61 |
| Motivation and attitude towards foreign languages: anxiety and confidence scores | MFQ | l2_anxiety | l2_confidence | -0.72 |
| Arithmetic abilities: subtractions accuracy and additions RT | RTT | rtt_sub_corr | rtt_sum_rt | -0.78 |
| Arithmetic abilities: subtractions RT and additions accuracy | RTT | rtt_sub_rt | rtt_sum_corr | -0.81 |
| Arithmetic abilities: additions accuracy and RT | RTT | rtt_sum_corr | rtt_sum_rt | -0.86 |
| Arithmetic abilities: additions RT and total accuracy | RTT | rtt_sum_rt | arith | -0.86 |
| Arithmetic abilities: subtractions RT and total accuracy | RTT | rtt_sub_rt | arith | -0.9 |
| Arithmetic abilities: subtractions accuracy and RT | RTT | rtt_sub_corr | rtt_sub_rt | -0.91 |

Table S3: Internal consistency values measured via Cronbach alpha, where a >= .5. Lower and upper bounds are reported at a 95% confidence interval. For clarity, values where a = > .6 are in bold.

| Test | Metric | Cronbach alpha | 95% CI lower bound | 95% CI upper bound |
|---|---|---|---|---|
| Finger tapping: non-dominant hand | **finger-tapping_nondominant** | **0.98** | 0.97 | 0.99 |
| Finger tapping: dominant hand | **finger-tapping_dominant** | **0.97** | 0.95 | 0.98 |
| Brocanto: RT block 3 | **brocanto_rt_3** | **0.96** | 0.94 | 0.97 |
| Brocanto: RT block 2 | **brocanto_rt_2** | **0.96** | 0.95 | 0.97 |
| Brocanto: RT block 1 | **brocanto_rt_1** | **0.95** | 0.93 | 0.96 |
| Purdue pegboard: assembly score | **purdue_assembly** | **0.93** | 0.90 | 0.95 |
| Text reading: RT | **reading_rt** | **0.93** | 0.89 | 0.95 |
| Rapid automatised naming: RT | **ran_rt** | **0.93** | 0.91 | 0.95 |
| MUSEBAQ: engaged (music) production score | **engaged_production** | **0.92** | 0.89 | 0.94 |
| MUSEBAQ: dance factor | **dance** | **0.90** | 0.85 | 0.93 |
| CVLT: total recall score | **cvlt_tot-recall** | **0.90** | 0.87 | 0.93 |
| Code switching: contextual | **swt_CS** | **0.89** | 0.84 | 0.92 |
| Purdue pegboard: both hands score | **purdue_both** | **0.89** | 0.84 | 0.92 |
| CVLT: total immediate recall score | **cvlt_tot-imm** | **0.88** | 0.84 | 0.91 |
| Raven's Advanced Progressive Matrices: RT | **apm_rt** | **0.88** | 0.85 | 0.91 |
| RTT: subtractions accuracy | **rtt-sub_corr** | **0.86** | 0.82 | 0.90 |
| MFQ: foreign language usage willingness | **usage_willingness** | **0.85** | 0.80 | 0.89 |
| MLAT5: RT | **mlat5_rt** | **0.85** | 0.81 | 0.89 |
| Purdue pegboard: dominant hand | **purdue_DH** | **0.85** | 0.80 | 0.90 |
| Artgram: RT | **artgram_rt** | **0.84** | 0.79 | 0.88 |
| MUSEBAQ: social connection factor | **social_connection** | **0.84** | 0.77 | 0.89 |
| RTT: additions accuracy | **rtt-sum_corr** | **0.84** | 0.80 | 0.89 |
| Purdue pegboard: non-dominant hand | **purdue_NDH** | **0.83** | 0.77 | 0.88 |
| CVLT: recognition RT | **cvlt_reco_rt** | **0.82** | 0.76 | 0.87 |
| MLAT5: accuracy | **mlat5_corr** | **0.82** | 0.77 | 0.87 |
| MLAT5: errors | **mlat5_incorr** | **0.82** | 0.77 | 0.87 |
| IRQ: visual factor | **irq_visual** | **0.81** | 0.75 | 0.86 |
| Adult Reading History score | **ahrq** | **0.81** | 0.75 | 0.86 |
| CVLT: recognition accuracy | **cvlt_reco_corr** | **0.81** | 0.76 | 0.86 |
| Brocanto accuracy: block 3 | **brocanto_corr_3** | **0.80** | 0.74 | 0.85 |
| IRQ: verbal factor | **irq_verbal** | **0.78** | 0.71 | 0.84 |
| MFQ: international contact | **intl_contact** | **0.78** | 0.70 | 0.84 |
| MFQ: confidence using foreign languages | **l2_confidence** | **0.77** | 0.69 | 0.84 |
| MFQ: foreign language environment | **milieu** | **0.76** | 0.68 | 0.83 |
| MUSEBAQ: cognitive and emotional regulation factor | **cognitive_emotional_regulation** | **0.72** | 0.63 | 0.80 |

| | | | | |
|---|---|---|---|---|
| Raven's Advanced Progressive Matrices: accuracy | **apm_corr** | **0.72** | 0.64 | 0.79 |
| Text reading: accuracy | **reading_acc** | **0.72** | 0.59 | 0.81 |
| MFQ: self idealisation as foreign language speaker | **ideal_l2_self** | **0.71** | 0.61 | 0.79 |
| Brocanto accuracy: block 2 | **brocanto_corr_2** | **0.70** | 0.61 | 0.78 |
| Hindi dental-retroflex task | **hindi** | **0.70** | 0.61 | 0.78 |
| MFQ: anxiety towards foreign languages | **l2_anxiety** | **0.69** | 0.57 | 0.78 |
| Code switching: unwilling/involuntary | **swt_US** | **0.68** | 0.56 | 0.78 |
| MUSEBAQ: physical exercise factor | **physical_exercise** | **0.67** | 0.53 | 0.76 |
| MFQ: interest for foreign languages | **l2_interest** | **0.67** | 0.55 | 0.76 |
| IRQ: orthographic factor | **irq_orto** | **0.66** | 0.55 | 0.75 |
| MUSEBAQ: music listening factor | **ind_mus_listening_iml** | **0.66** | 0.50 | 0.77 |
| AMMA: total score | **amma** | **0.65** | 0.54 | 0.74 |
| MFQ: instrumentality of foreign languages | **instrumentality** | **0.65** | 0.54 | 0.75 |
| IRQ: manipulation factor | **irq_manip** | **0.65** | 0.54 | 0.75 |
| Spelling test: accuracy | **spelling** | **0.62** | 0.46 | 0.73 |
| Digit span: backward score | **digit-back** | **0.61** | 0.49 | 0.71 |
| Digit span: forward score | digit-for | 0.58 | 0.44 | 0.69 |
| MUSEBAQ: index of musical training | ind_mus_train_imt | 0.57 | 0.40 | 0.70 |
| Corsi Blocks: forward score | corsi-for | 0.55 | 0.42 | 0.67 |
| Artgram: accuracy | artgram_corr | 0.54 | 0.39 | 0.66 |
| ANT-I inhibition gain | **ANT-I inhibition** | **0.67*, CI 95%, 0.55, 0.77** | | |
| ANT-I re-orienting gain | **ANT-I re-orienting** | **0.65*, CI 95% 0.51, 0.76** | | |
| ANT-I orienting gain | **ANT-I orienting** | **0.61*, CI 95% 0.46, 0.73** | | |
| ANT-I alerting gain | ANT-I alerting | 0.51*, CI 95% 0.32, 0.66 | | |
| Farsi uvular production: native likeness score (accuracy) | **Farsi uvular production** | ***Inter-rater reliability:*** $r_{pearson}$**(95): 0.68** $p < .0001$ | | |

A. Rampinini, I. Balboni, O. Kepinska, R. Berthele, N. Golestani

Table S4
Descriptives of the z-scored Questionnaire data.

| | *Range* | *Skewness* | *Kurtosis* |
|---|---|---|---|
| *entropy_competence_speak* | 6.57 | -0.20 | 1.31 |
| *entropy_competence_read* | 6.53 | -0.20 | 1.43 |
| *entropy_competence_compr* | 6.67 | -0.11 | 1.84 |
| *entropy_curr_tot_exp* | 3.92 | -0.07 | -0.71 |
| *ahrq_score* | 5.60 | 0.68 | 0.69 |
| *manipulationfactor* | 5.39 | -0.51 | 0.35 |
| *orthohraphicfactor* | 4.59 | 0.29 | -0.15 |
| *verbalfactor* | 4.35 | -0.32 | -0.31 |
| *visualfactor* | 5.48 | -0.86 | 1.16 |
| *ideal_l2_self* | 4.38 | -0.84 | 0.20 |
| *instrumentality* | 4.47 | -0.55 | 0.49 |
| *intl_contact* | 4.52 | -1.25 | 1.42 |
| *l2_interest* | 4.91 | -0.89 | 0.75 |
| *l2_anxiety* | 4.31 | 0.10 | -0.63 |
| *l2_confidence* | 4.21 | -0.27 | -0.32 |
| *milieu* | 4.72 | -1.67 | 3.39 |
| *usage_willingness* | 4.95 | -0.52 | 0.13 |
| *ind_mus_train_imt* | 3.85 | 0.46 | -0.78 |
| *ind_mus_listening_iml* | 3.91 | 0.24 | -0.52 |
| *mus_instr_play_imip* | 9.74 | 8.70 | 79.20 |
| *cognitive and emotional regulation* | 4.38 | -0.35 | -0.31 |
| *social connection* | 4.48 | -0.29 | -0.42 |
| *engaged production* | 3.52 | 1.20 | 0.18 |
| *dance* | 2.96 | 0.35 | -1.36 |
| *physical exercise* | 4.88 | -1.01 | 1.62 |
| *hand_index* | 4.42 | -1.94 | 3.51 |
| *bsmss* | 3.96 | -0.45 | -0.58 |

Table S5
Descriptives of the z-scored Task data.

| | *Range* | *Skewness* | *Kurtosis* |
|---|---|---|---|
| *purdue_dh_avg* | 4.22 | -0.16 | -0.75 |
| *purdue_ndh_avg* | 4.66 | 0.05 | -0.34 |
| *purdue_both_avg* | 5.39 | -0.53 | 0.38 |
| *purdue_dh_ndh_both_avg* | 4.58 | -0.39 | -0.53 |
| *purdue_assembly_avg* | 5.25 | -0.53 | 0.28 |
| *ran_tot_acc* | 8.07 | -4.71 | 28.72 |
| *ran_tot_rt* | 6.47 | 1.99 | 6.45 |
| *reading_text_acc* | 5.92 | -2.09 | 5.54 |
| *reading_text_rt* | 5.46 | 1.19 | 2.03 |

| | | | |
|---|---|---|---|
| *cvlt_tot_imm* | 4.91 | -0.61 | 0.42 |
| *cvlt_tot_recall* | 4.98 | -0.74 | 0.70 |
| *spelling_regular_acc* | 4.21 | -0.92 | 0.32 |
| *spelling_irregular_acc* | 4.52 | -0.38 | -0.48 |
| *spelling_pseudo_acc* | 5.02 | -0.41 | -0.13 |
| *spelling_tot_acc* | 4.19 | -0.62 | -0.24 |
| *spoon_acc* | 4.94 | -2.61 | 6.76 |
| *spoon_rt_manual* | 4.91 | 1.78 | 3.49 |
| *regular_acc* | 8.80 | -7.38 | 56.13 |
| *irregular_acc* | 10.50 | -7.06 | 58.74 |
| *pseudo_acc* | 7.11 | -5.41 | 32.13 |
| *regular_rt* | 6.59 | 1.72 | 5.24 |
| *irregular_rt* | 5.46 | 1.49 | 2.69 |
| *pseudo_rt* | 5.55 | 1.28 | 2.59 |
| *phon_suppr_acc* | 5.29 | -2.21 | 5.78 |
| *wordreading* | 8.74 | -6.85 | 49.85 |
| *alerting* | 4.33 | 0.33 | -0.52 |
| *orienting* | 5.64 | 0.00 | 0.17 |
| *reorienting* | 5.35 | 0.65 | 0.48 |
| *inhibition* | 5.10 | 0.47 | 0.18 |
| *rtt_sub_corr* | 4.54 | -0.17 | -0.40 |
| *rtt_sub_incorr* | 5.06 | 1.40 | 2.21 |
| *rtt_sub_rt* | 5.50 | 1.05 | 1.32 |
| *finger_tapping_dominant* | 5.34 | 0.42 | 0.85 |
| *finger_tapping_nondominant* | 5.85 | 0.63 | 0.76 |
| *digit_for_span* | 4.09 | 0.19 | -0.56 |
| *digit_back_span* | 4.93 | 0.54 | -0.05 |
| *rtt_sum_corr* | 4.34 | -0.30 | -0.74 |
| *rtt_sum_incorr* | 5.28 | 1.43 | 2.80 |
| *rtt_sum_rt* | 6.13 | 1.48 | 3.57 |
| *corsi_back_span* | 3.35 | 0.10 | -0.90 |
| *brocanto_corr2* | 4.70 | 0.20 | -0.54 |
| *brocanto_incorr2* | 4.77 | -0.12 | -0.46 |
| *brocanto_rt2* | 5.11 | -0.54 | 0.39 |
| *brocanto_corr3* | 4.09 | 0.29 | -0.68 |
| *brocanto_incorr3* | 4.18 | -0.25 | -0.72 |
| *brocanto_rt3* | 5.23 | -0.45 | 0.25 |
| *mlat5_corr* | 4.01 | -0.64 | -0.26 |
| *mlat5_incorr* | 4.02 | 0.61 | -0.37 |
| *mlat5_rt* | 5.46 | 1.12 | 1.64 |
| *artgram_corr* | 5.14 | -0.57 | 0.06 |
| *artgram_incorr* | 5.29 | 0.74 | 0.59 |
| *artgram_rt* | 6.59 | 1.60 | 4.12 |

| | | | |
|---|---|---|---|
| *corsi_for_span* | 4.02 | -0.32 | -0.45 |
| *amma_tonal* | 4.46 | 0.15 | -0.44 |
| *amma_rhythm* | 4.84 | -0.01 | -0.10 |
| *amma_total* | 4.69 | 0.09 | -0.29 |
| *hindi_score_weighted* | 4.06 | 0.84 | -0.41 |
| *arith* | 4.54 | -0.19 | -0.68 |
| *fingertap* | 5.26 | 0.43 | 0.41 |
| *span_visual* | 4.78 | -0.04 | -0.41 |
| *span_verbal* | 4.92 | 0.64 | -0.09 |

A. Rampinini, I. Balboni, O. Kepinska, R. Berthele, N. Golestani

Figure S2

Correlation matrix of the z-scored behavioural data with only significant correlations highlighted.

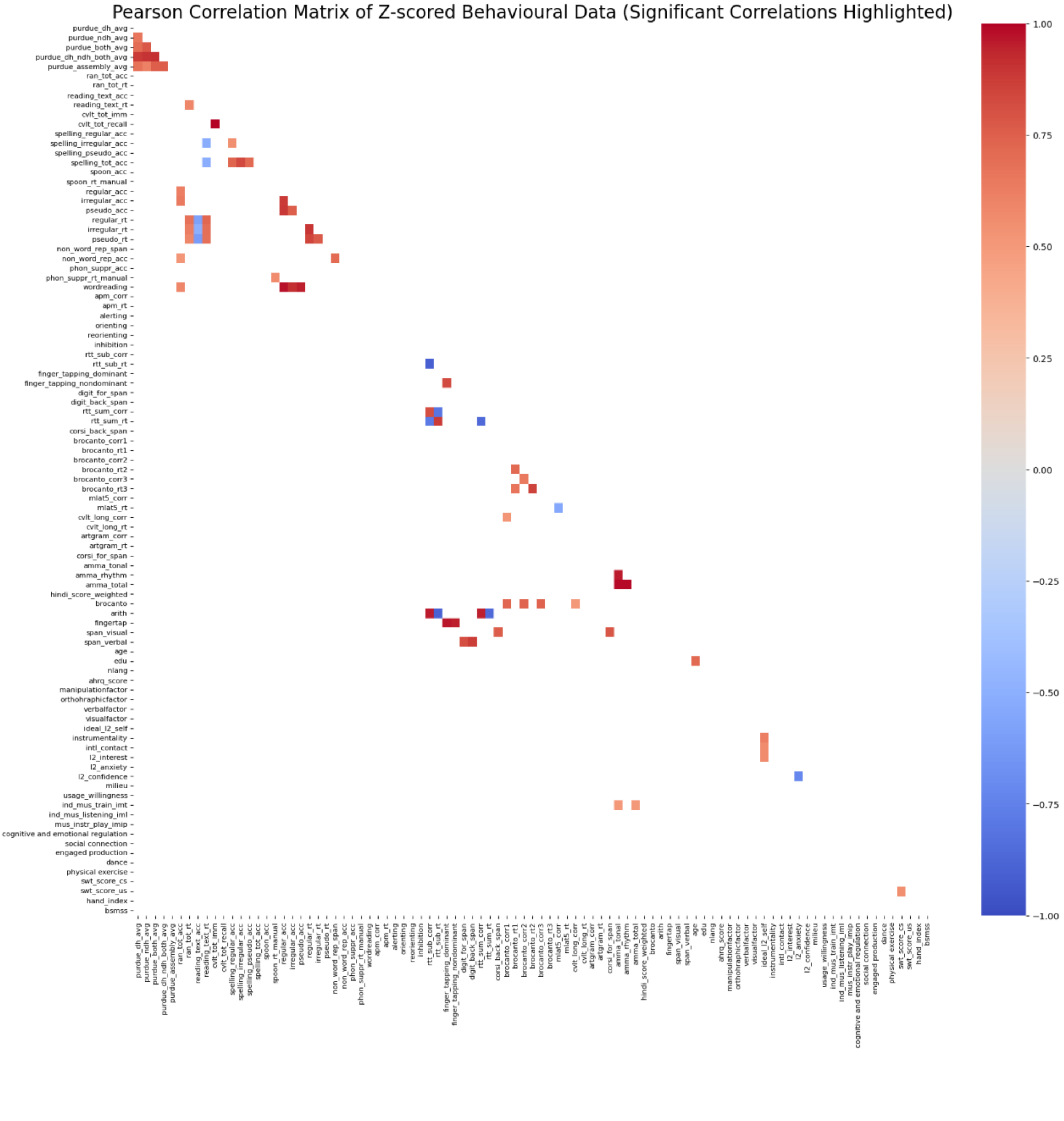

**Supplementary information v2.0 OCT-24: NEBULA101: an open dataset for the study of language aptitude in behaviour, brain structure and function**

A. Rampinini, I. Balboni, O. Kepinska, R. Berthele, N. Golestani

Figure S3
Present and missing data at a glance. Yellow squares indicate present data, purple squares indicate missing data. Subject codes are on the y axis, task names on the x axis.

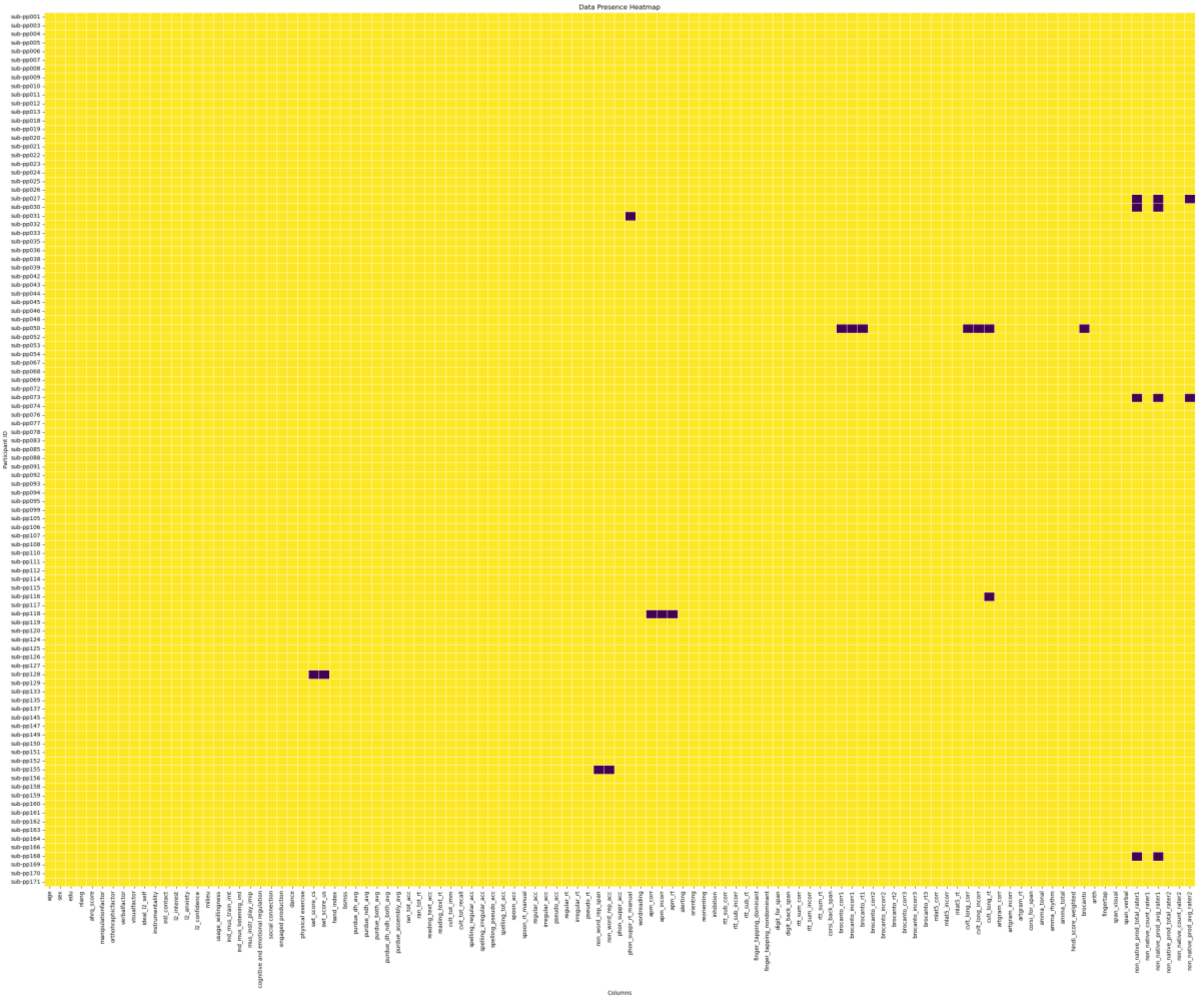

A. Rampinini, I. Balboni, O. Kepinska, R. Berthele, N. Golestani

Figure S4. Entropy violin plots: distribution shape of entropy scores.

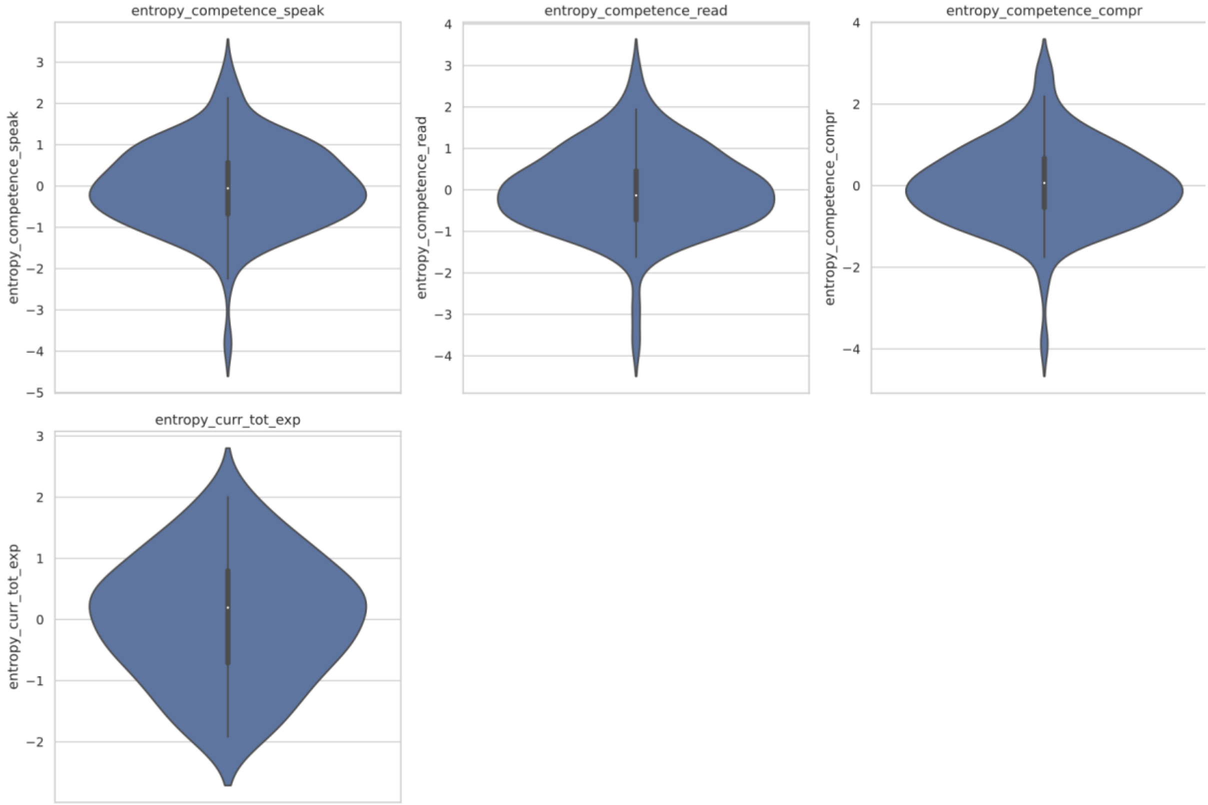

Figure S5. Questionnaire violin plots: distriution shape of questionnaire scores.

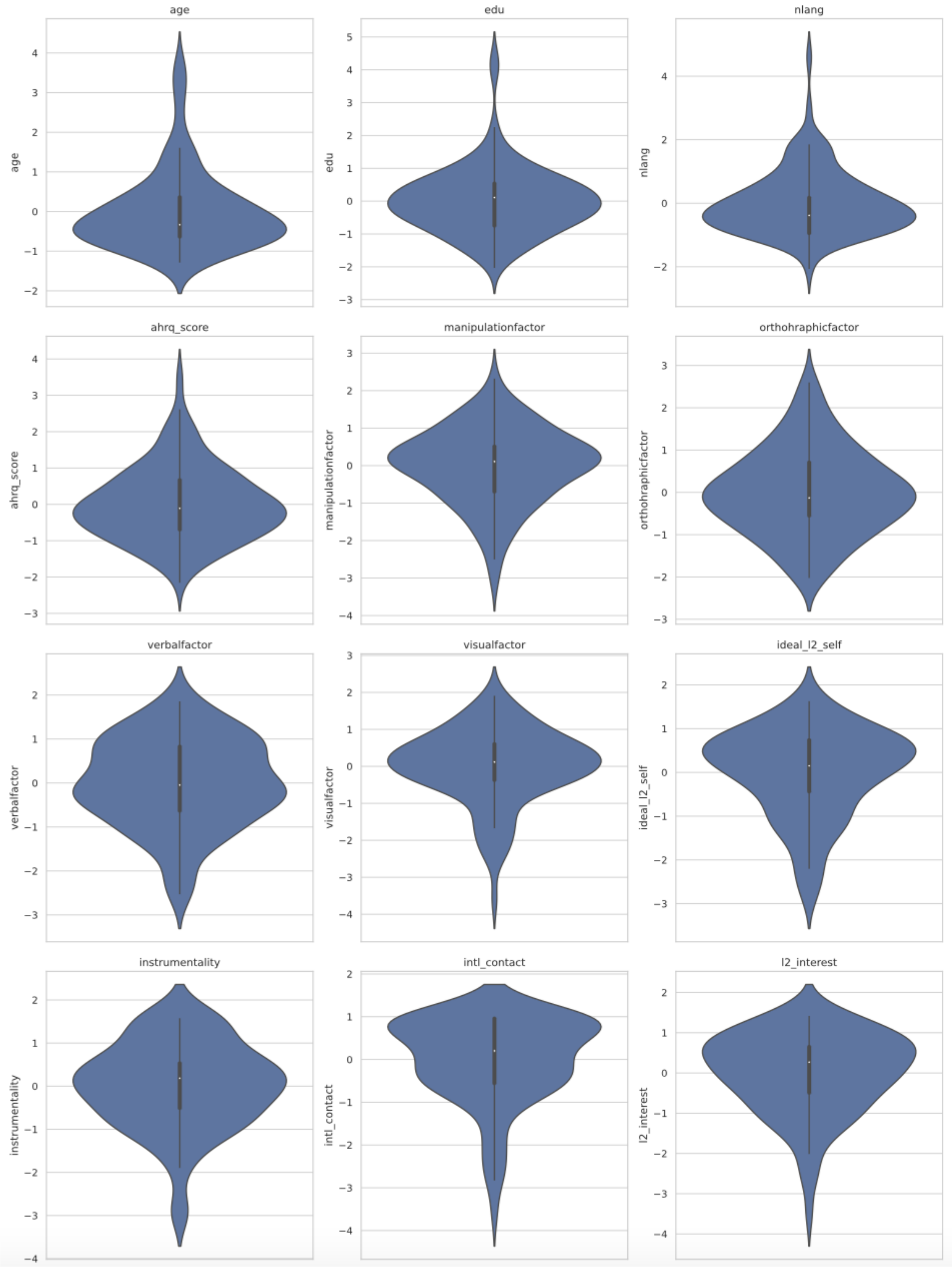

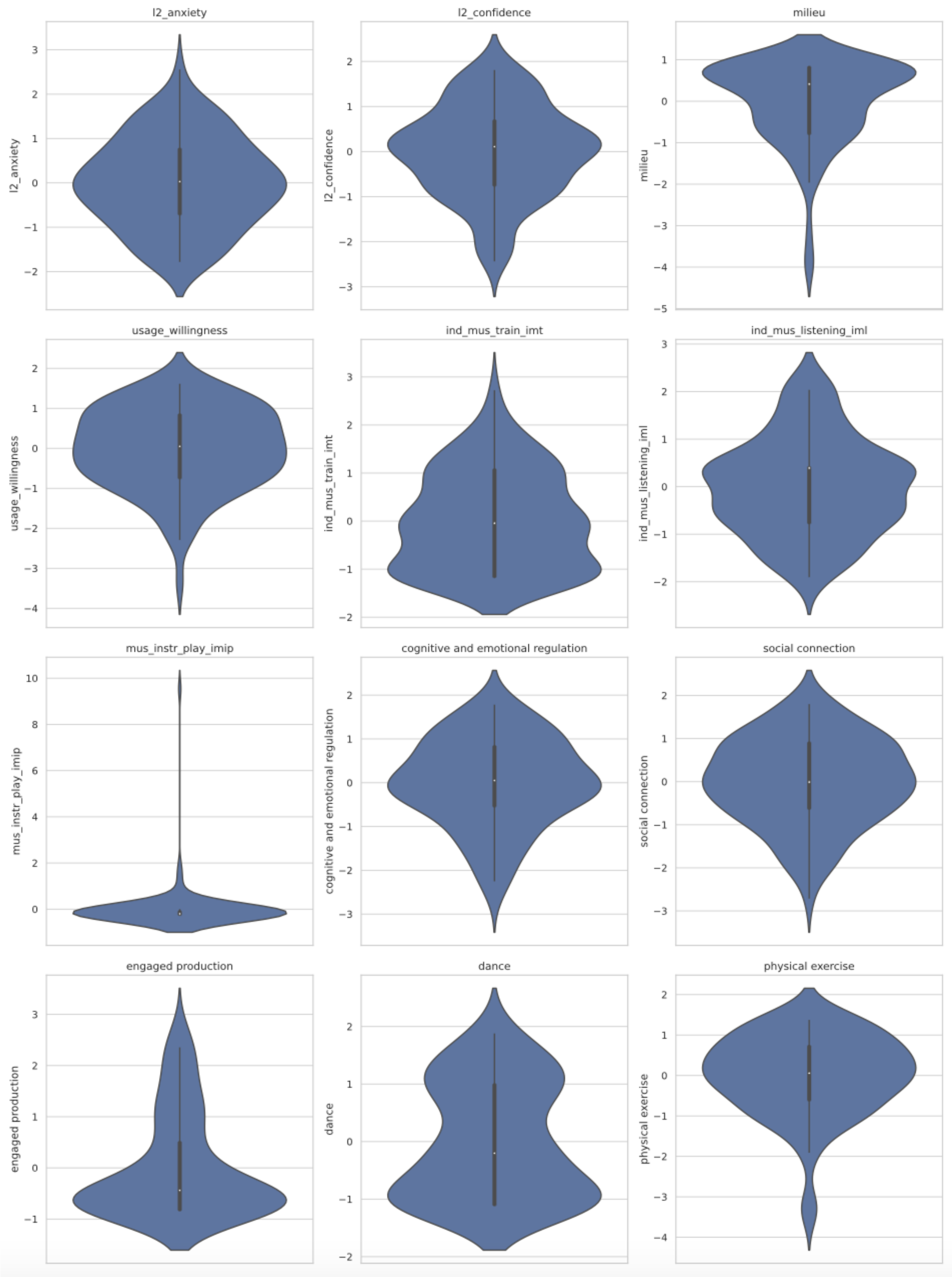
l2_anxiety
l2_confidence
milieu
usage_willingness
ind_mus_train_imt
ind_mus_listening_iml
mus_instr_play_imip
cognitive and emotional regulation
social connection
engaged production
dance
physical exercise

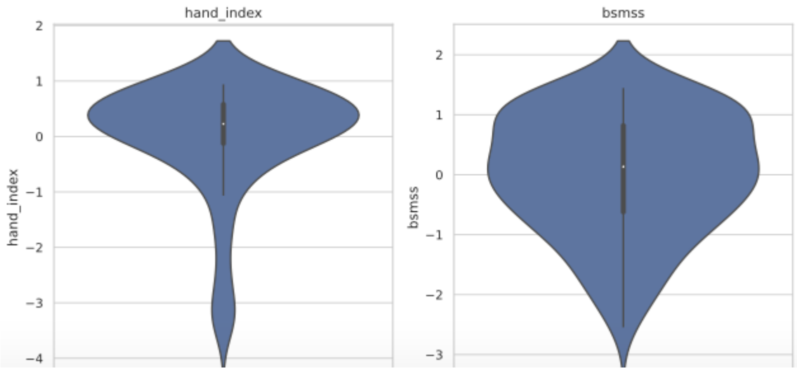
hand_index
bsmss
hand_index
bsmss
2
1
0
−1
−2
−3
−4
2
1
0
−1
−2
−3

Figure S6. Task violin plots. Distribution shape of task scores.

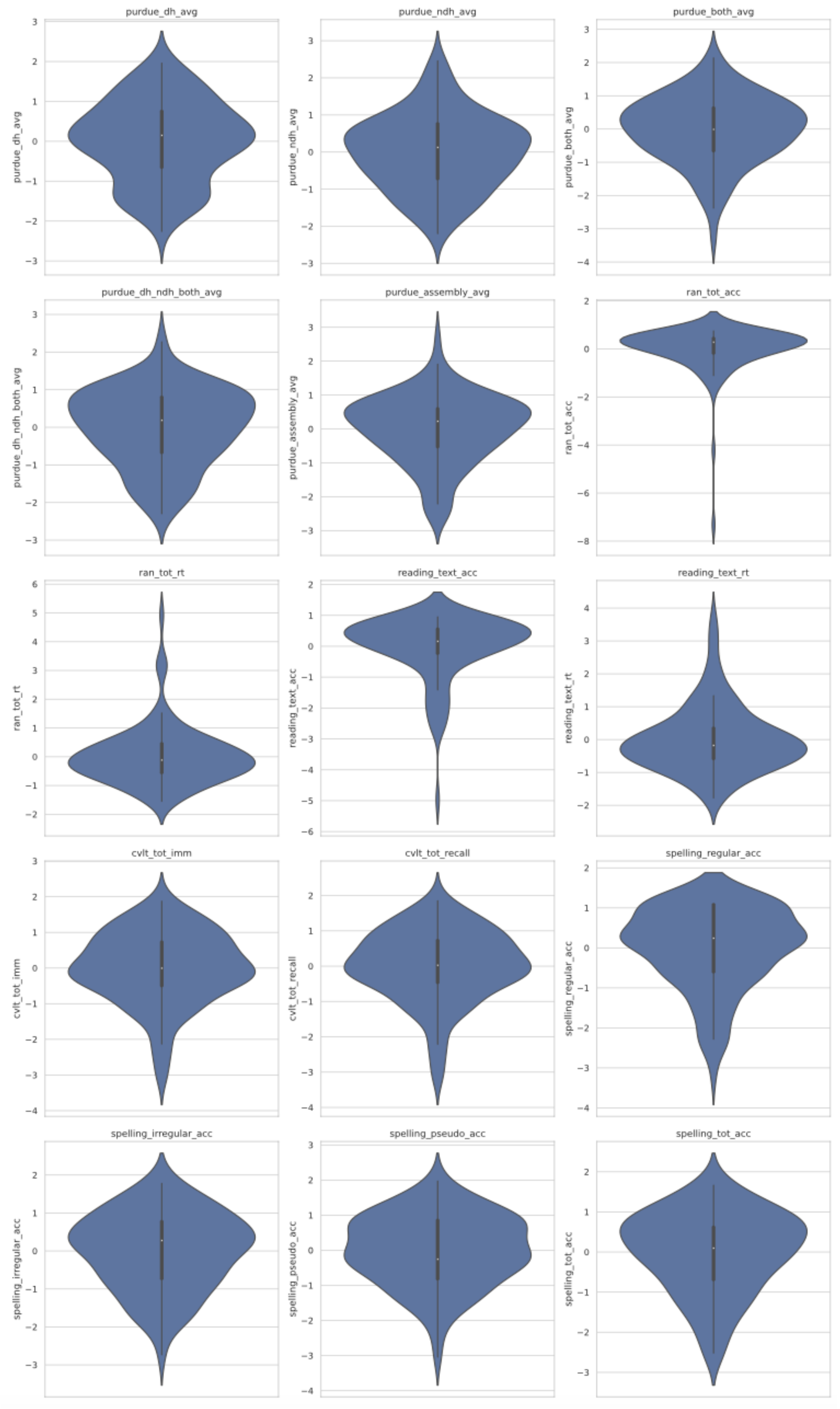

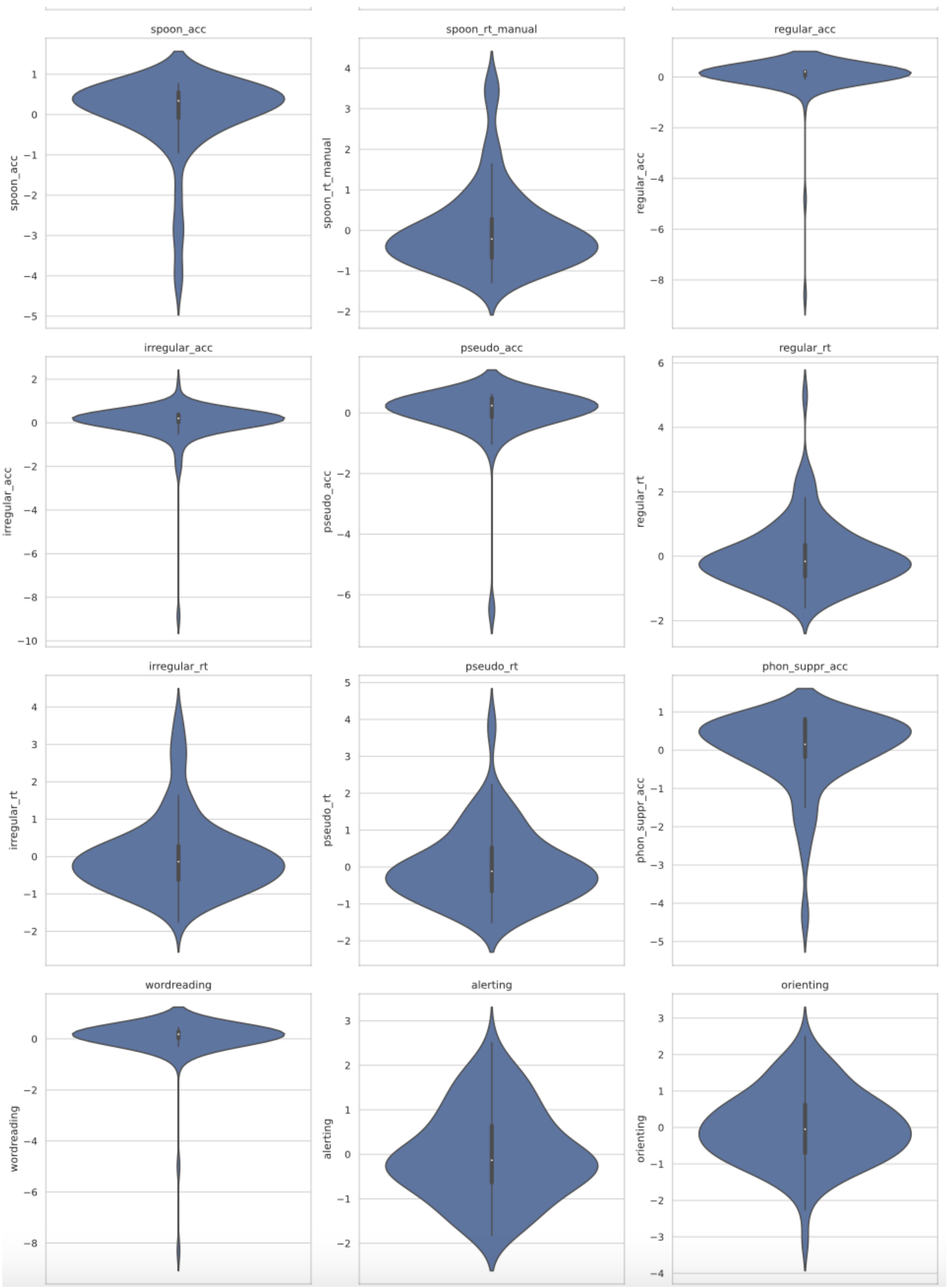
spoon_acc
spoon_rt_manual
regular_acc
irregular_acc
pseudo_acc
regular_rt
irregular_rt
pseudo_rt
phon_suppr_acc
wordreading
alerting
orienting

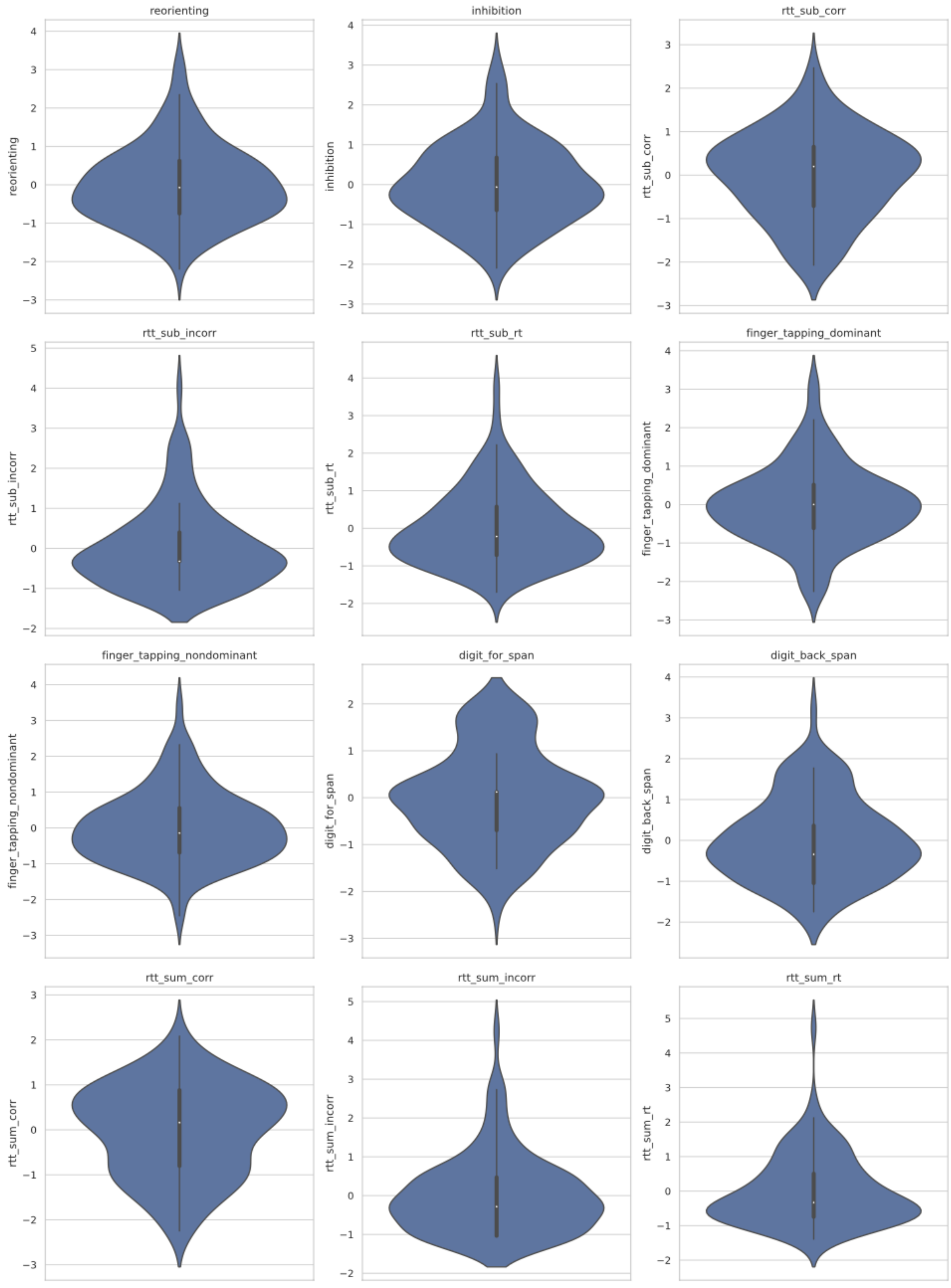
reorienting
inhibition
rtt_sub_corr
rtt_sub_incorr
rtt_sub_rt
finger_tapping_dominant
finger_tapping_nondominant
digit_for_span
digit_back_span
rtt_sum_corr
rtt_sum_incorr
rtt_sum_rt

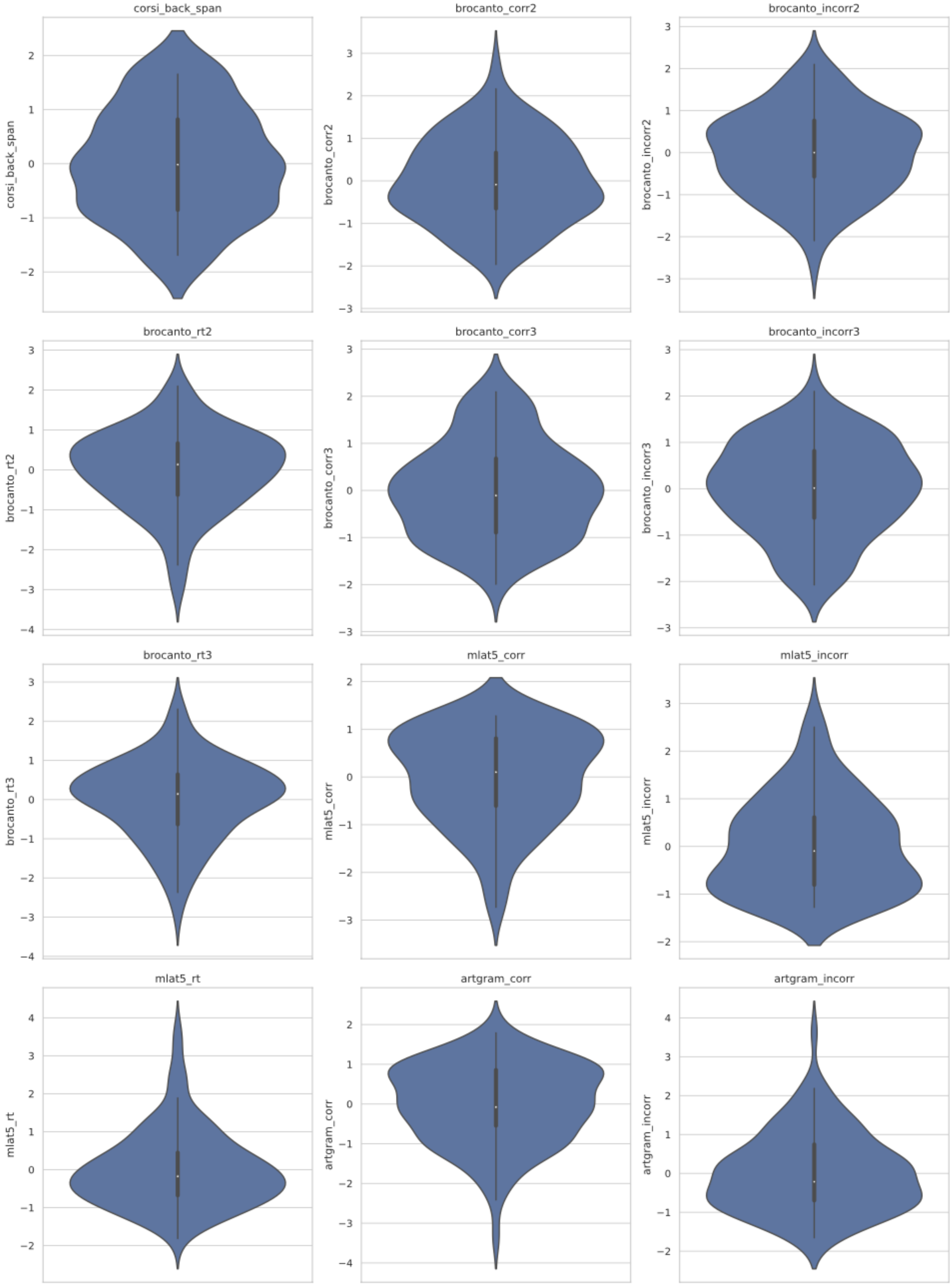
corsi_back_span
brocanto_corr2
brocanto_incorr2
brocanto_rt2
brocanto_corr3
brocanto_incorr3
brocanto_rt3
mlat5_corr
mlat5_incorr
mlat5_rt
artgram_corr
artgram_incorr

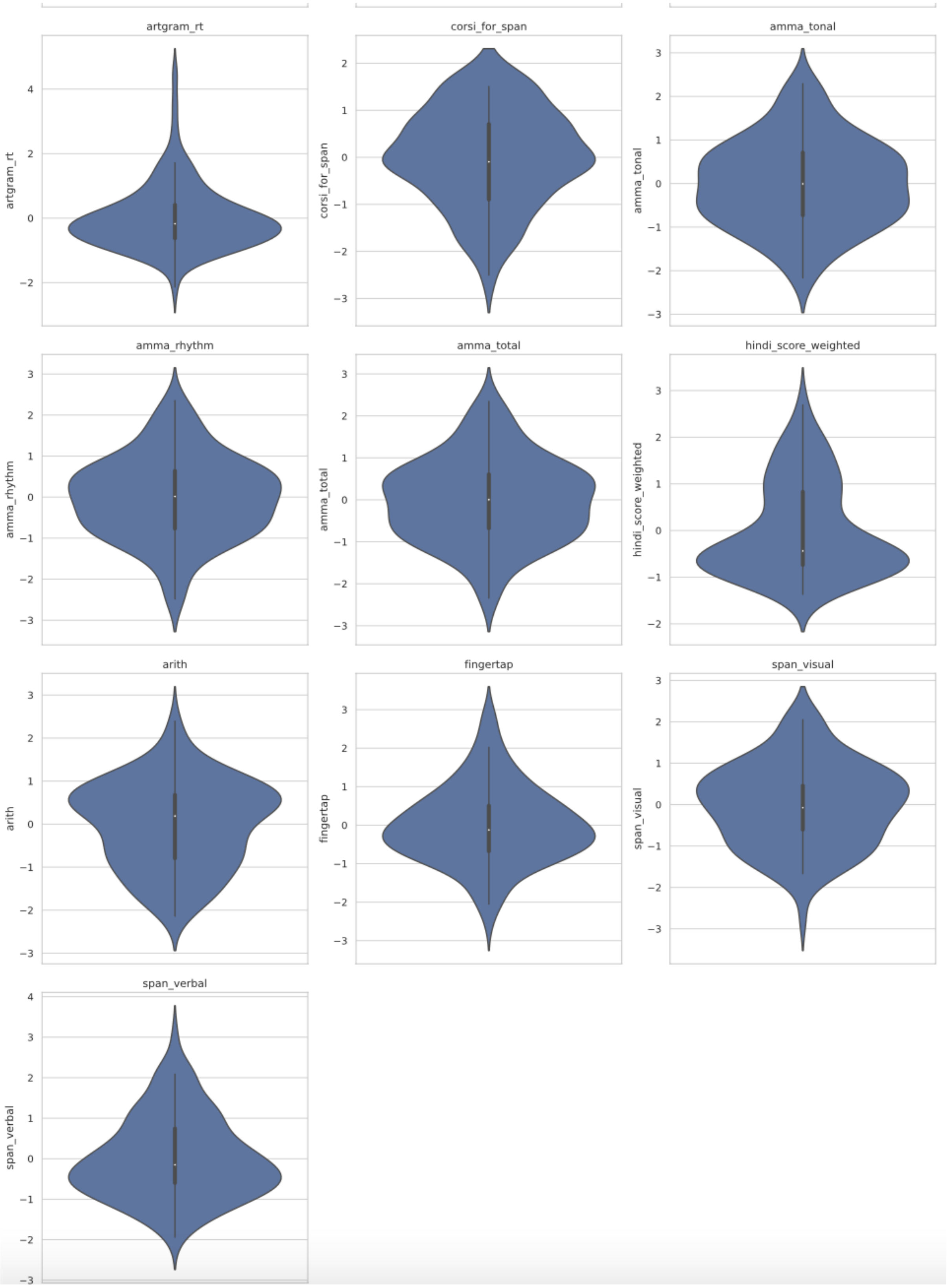
artgram_rt
corsi_for_span
amma_tonal
amma_rhythm
amma_total
hindi_score_weighted
arith
fingertap
span_visual
span_verbal

### ANTi_reliability_NEBULA101

### IB

### 5/8/2024

Reliability assessment for the task ANT-i was computed with split-half correlations using the package splithalf

### Cleaning and preprocessing

We clean and preprocess the data as we did for the scoring but this time create 1 variable containing all of the participants with cleaned scores divided by blocks to be used for the split-half correlation. Max responses per participant 432. Responses missing could be due to lack of attempt, abnormal timing, and incorrect responses are removed

select only relevant pp

```
pp_toinclude=read.delim('/data/team/Aptitude/nebula101/participants.tsv')
final_data <- final_data %>%
  filter(participant_id %in% pp_toinclude$participant_id)
```

##Split-half correlation

```
difference <- splithalf(data = final_data,
                        outcome = "RT",
                        score = "difference",
                        halftype = "random",
                        permutations = 10000,
                        var.RT = "ant_RT",
                        var.participant = "participant_id",
                        var.compare = "alerting.code",
                        compare1 = "0",
                        compare2 = "1",
                        average = "mean",
                        plot = TRUE)
```

```
|
|===================                                                    |  28%
|
|====================                                                   |  29%
|
|=====================                                                  |  30%
|
|=====================                                                  |  31%
|
|======================                                                 |  32%
|
|=======================                                                |  33%
|
|========================                                               |  34%
|
|========================                                               |  35%
|
|=========================                                              |  36%
|
|==========================                                             |  37%
|
|==========================                                             |  38%
|
|===========================                                            |  39%
|
|============================                                           |  40%
|
|============================                                           |  41%
|
|=============================                                          |  42%
|
|==============================                                         |  43%
|
|==============================                                         |  44%
|
|===============================                                        |  45%
|
|================================                                       |  46%
|
|=================================                                      |  47%
|
|=================================                                      |  48%
|
|==================================                                     |  49%
|
|===================================                                    |  50%
|
|====================================                                   |  51%
|
|=====================================                                  |  52%
|
|=====================================                                  |  53%
|
|======================================                                 |  54%
|
|=======================================                                |  55%
```

```
|
|======================================                             |  56%
|
|======================================                             |  57%
|
|=======================================                            |  58%
|
|========================================                           |  59%
|
|========================================                           |  60%
|
|=========================================                          |  61%
|
|==========================================                         |  62%
|
|==========================================                         |  63%
|
|===========================================                        |  64%
|
|============================================                       |  65%
|
|============================================                       |  66%
|
|=============================================                      |  67%
|
|==============================================                     |  68%
|
|===============================================                    |  69%
|
|===============================================                    |  70%
|
|================================================                   |  71%
|
|=================================================                  |  72%
|
|=================================================                  |  73%
|
|==================================================                 |  74%
|
|===================================================                |  75%
|
|===================================================                |  76%
|
|====================================================               |  77%
|
|=====================================================              |  78%
|
|=====================================================              |  79%
|
|======================================================             |  80%
|
|=======================================================            |  81%
|
|========================================================           |  82%
|
|========================================================           |  83%
```

```
  |
  |===========================================================           |  84%
  |
  |============================================================          |  85%
  |
  |============================================================          |  86%
  |
  |=============================================================         |  87%
  |
  |==============================================================        |  88%
  |
  |==============================================================        |  89%
  |
  |===============================================================       |  90%
  |
  |================================================================      |  91%
  |
  |================================================================      |  92%
  |
  |=================================================================     |  93%
  |
  |==================================================================    |  94%
  |
  |===================================================================   |  95%
  |
  |===================================================================   |  96%
  |
  |====================================================================  |  97%
  |
  |===================================================================== |  98%
  |
  |===================================================================== |  99%
  |
  |======================================================================| 100%[1] "c
ondition all complete"
## [1] "Calculating split half estimates"
## [1] "split half estimates for 10000 random splits"
##   condition   n spearmanbrown SB_low SB_high
## 1       all 101          0.51   0.32    0.66
## [1] "this could be reported as: using 10000 random splits, the spearman-brown corr
ected reliability  estimate for the all condition was 0.51, 95% CI [0.32, 0.66]"
```

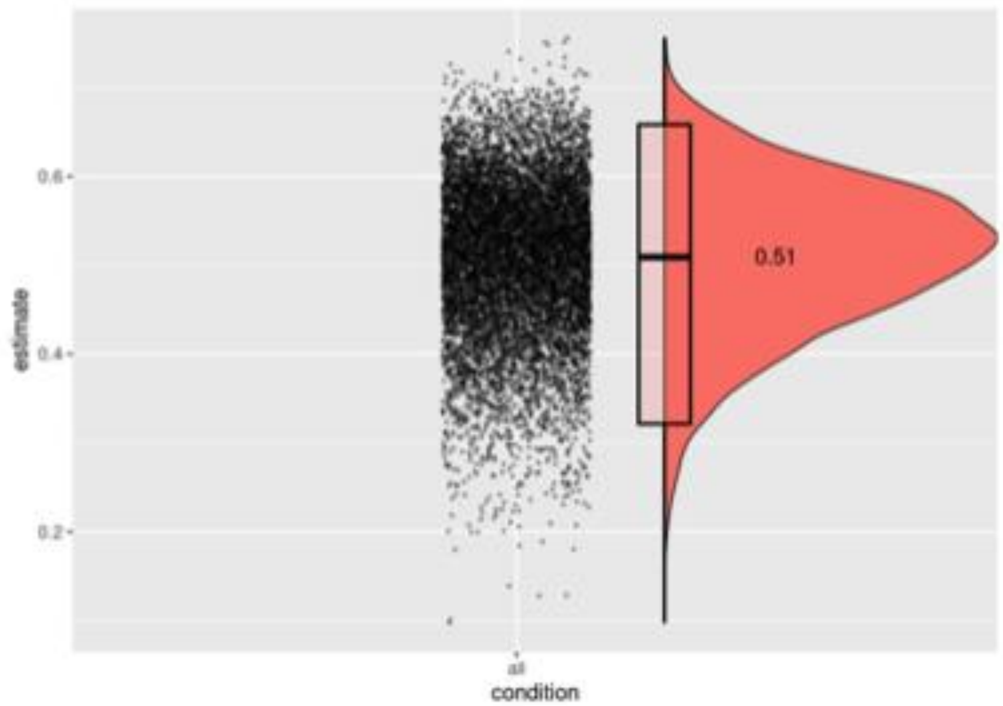


```
difference_orienting <- splithalf(data = final_data,
                        outcome = "RT",
                        score = "difference",
                        halftype = "random",
                        permutations = 10000,
                        var.RT = "ant_RT",
                        var.participant = "participant_id",
                        var.compare = "cued.code",
                        compare1 = "0",
                        compare2 = "1",
                        average = "mean",
                        plot = TRUE)
```

```
##
  |
  |                                                                      |   0%
  |
  |=                                                                     |   2%
  |
  |==                                                                    |   3%
  |
  |===                                                                   |   4%
  |
  |===                                                                   |   5%
  |
  |====                                                                  |   6%
  |
  |=====                                                                 |   7%
  |
  |======                                                                |   8%
  |
  |======                                                                |   9%
  |
  |=======                                                               |  10%
  |
  |========                                                              |  11%
  |
  |========                                                              |  12%
  |
  |=========                                                             |  13%
  |
  |==========                                                            |  14%
  |
  |==========                                                            |  15%
  |
  |===========                                                           |  16%
  |
  |============                                                          |  17%
  |
  |============                                                          |  18%
  |
  |=============                                                         |  19%
  |
  |==============                                                        |  20%
  |
  |===============                                                       |  21%
  |
  |===============                                                       |  22%
  |
  |================                                                      |  23%
  |
  |=================                                                     |  24%
  |
  |=================                                                     |  25%
  |
  |==================                                                    |  26%
  |
  |===================                                                   |  27%
```

```
|
|===================                                                   |  28%
|
|====================                                                  |  29%
|
|=====================                                                 |  30%
|
|=====================                                                 |  31%
|
|======================                                                |  32%
|
|=======================                                               |  33%
|
|========================                                              |  34%
|
|========================                                              |  35%
|
|=========================                                             |  36%
|
|==========================                                            |  37%
|
|==========================                                            |  38%
|
|===========================                                           |  39%
|
|============================                                          |  40%
|
|============================                                          |  41%
|
|=============================                                         |  42%
|
|==============================                                        |  43%
|
|==============================                                        |  44%
|
|===============================                                       |  45%
|
|================================                                      |  46%
|
|=================================                                     |  47%
|
|=================================                                     |  48%
|
|==================================                                    |  49%
|
|===================================                                   |  50%
|
|====================================                                  |  51%
|
|=====================================                                 |  52%
|
|=====================================                                 |  53%
|
|======================================                                |  54%
|
|=======================================                               |  55%
```

```
|
|========================================                              |  56%
|
|========================================                              |  57%
|
|=========================================                             |  58%
|
|==========================================                            |  59%
|
|==========================================                            |  60%
|
|===========================================                           |  61%
|
|============================================                          |  62%
|
|============================================                          |  63%
|
|=============================================                         |  64%
|
|==============================================                        |  65%
|
|==============================================                        |  66%
|
|===============================================                       |  67%
|
|================================================                      |  68%
|
|=================================================                     |  69%
|
|=================================================                     |  70%
|
|==================================================                    |  71%
|
|===================================================                   |  72%
|
|===================================================                   |  73%
|
|====================================================                  |  74%
|
|=====================================================                 |  75%
|
|=====================================================                 |  76%
|
|======================================================                |  77%
|
|=======================================================               |  78%
|
|=======================================================               |  79%
|
|========================================================              |  80%
|
|=========================================================             |  81%
|
|==========================================================            |  82%
|
|==========================================================            |  83%
```

```
  |                                                                      
  |===========================================================           |  84%
  |                                                                      
  |============================================================          |  85%
  |                                                                      
  |============================================================          |  86%
  |                                                                      
  |=============================================================         |  87%
  |                                                                      
  |==============================================================        |  88%
  |                                                                      
  |==============================================================        |  89%
  |                                                                      
  |===============================================================       |  90%
  |                                                                      
  |================================================================      |  91%
  |                                                                      
  |================================================================      |  92%
  |                                                                      
  |=================================================================     |  93%
  |                                                                      
  |==================================================================    |  94%
  |                                                                      
  |===================================================================   |  95%
  |                                                                      
  |===================================================================   |  96%
  |                                                                      
  |====================================================================  |  97%
  |                                                                      
  |===================================================================== |  98%
  |                                                                      
  |===================================================================== |  99%
  |                                                                      
  |======================================================================| 100%[1] "c
ondition all complete"
## [1] "Calculating split half estimates"
## [1] "split half estimates for 10000 random splits"
##   condition   n spearmanbrown SB_low SB_high
## 1       all 101          0.61   0.46    0.73
## [1] "this could be reported as: using 10000 random splits, the spearman-brown corr
ected reliability  estimate for the all condition was 0.61, 95% CI [0.46, 0.73]"
```

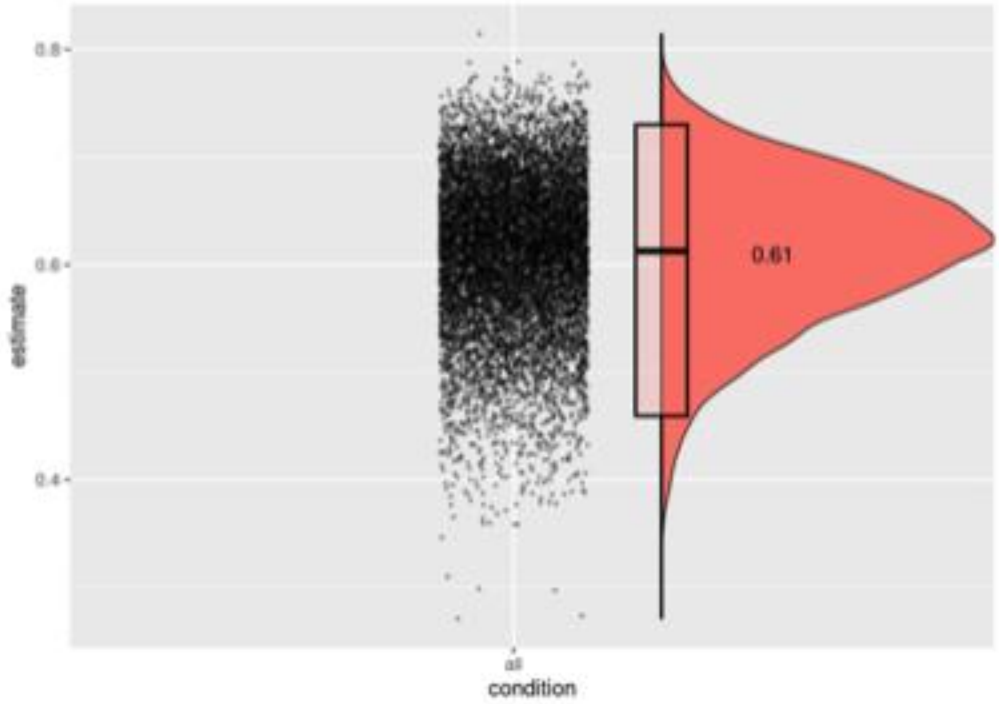


```
difference_reorienting <- splithalf(data = final_data,
                          outcome = "RT",
                          score = "difference",
                          halftype = "random",
                          permutations = 10000,
                          var.RT = "ant_RT",
                          var.participant = "participant_id",
                          var.compare = "cued.code",
                          compare1 = "2",
                          compare2 = "1",
                          average = "mean",
                          plot = TRUE)
```

```
##
  |
  |                                                                      |   0%
  |
  |=                                                                     |   2%
  |
  |==                                                                    |   3%
  |
  |===                                                                   |   4%
  |
  |===                                                                   |   5%
  |
  |====                                                                  |   6%
  |
  |=====                                                                 |   7%
  |
  |======                                                                |   8%
  |
  |======                                                                |   9%
  |
  |=======                                                               |  10%
  |
  |========                                                              |  11%
  |
  |========                                                              |  12%
  |
  |=========                                                             |  13%
  |
  |==========                                                            |  14%
  |
  |==========                                                            |  15%
  |
  |===========                                                           |  16%
  |
  |============                                                          |  17%
  |
  |============                                                          |  18%
  |
  |=============                                                         |  19%
  |
  |==============                                                        |  20%
  |
  |===============                                                       |  21%
  |
  |===============                                                       |  22%
  |
  |================                                                      |  23%
  |
  |=================                                                     |  24%
  |
  |=================                                                     |  25%
  |
  |==================                                                    |  26%
  |
  |===================                                                   |  27%
```

```
|
|===================                                                   |  28%
|
|====================                                                  |  29%
|
|=====================                                                 |  30%
|
|=====================                                                 |  31%
|
|======================                                                |  32%
|
|=======================                                               |  33%
|
|========================                                              |  34%
|
|========================                                              |  35%
|
|=========================                                             |  36%
|
|==========================                                            |  37%
|
|==========================                                            |  38%
|
|===========================                                           |  39%
|
|============================                                          |  40%
|
|============================                                          |  41%
|
|=============================                                         |  42%
|
|==============================                                        |  43%
|
|==============================                                        |  44%
|
|===============================                                       |  45%
|
|================================                                      |  46%
|
|=================================                                     |  47%
|
|=================================                                     |  48%
|
|==================================                                    |  49%
|
|===================================                                   |  50%
|
|====================================                                  |  51%
|
|=====================================                                 |  52%
|
|=====================================                                 |  53%
|
|======================================                                |  54%
|
|=======================================                               |  55%
```

```
|
|========================================                              |  56%
|
|========================================                              |  57%
|
|=========================================                             |  58%
|
|==========================================                            |  59%
|
|==========================================                            |  60%
|
|===========================================                           |  61%
|
|============================================                          |  62%
|
|============================================                          |  63%
|
|=============================================                         |  64%
|
|==============================================                        |  65%
|
|==============================================                        |  66%
|
|===============================================                       |  67%
|
|================================================                      |  68%
|
|=================================================                     |  69%
|
|=================================================                     |  70%
|
|==================================================                    |  71%
|
|===================================================                   |  72%
|
|===================================================                   |  73%
|
|====================================================                  |  74%
|
|=====================================================                 |  75%
|
|=====================================================                 |  76%
|
|======================================================                |  77%
|
|=======================================================               |  78%
|
|=======================================================               |  79%
|
|========================================================              |  80%
|
|=========================================================             |  81%
|
|==========================================================            |  82%
|
|==========================================================            |  83%
```

```
  |
  |==========================================================           |  84%
  |
  |===========================================================          |  85%
  |
  |===========================================================          |  86%
  |
  |============================================================         |  87%
  |
  |=============================================================        |  88%
  |
  |=============================================================        |  89%
  |
  |==============================================================       |  90%
  |
  |===============================================================      |  91%
  |
  |===============================================================      |  92%
  |
  |================================================================     |  93%
  |
  |=================================================================    |  94%
  |
  |==================================================================   |  95%
  |
  |==================================================================   |  96%
  |
  |===================================================================  |  97%
  |
  |==================================================================== |  98%
  |
  |==================================================================== |  99%
  |
  |=====================================================================| 100%[1] "c
ondition all complete"
## [1] "Calculating split half estimates"
## [1] "split half estimates for 10000 random splits"
##   condition   n spearmanbrown SB_low SB_high
## 1       all 101          0.65   0.51    0.76
## [1] "this could be reported as: using 10000 random splits, the spearman-brown corr
ected reliability  estimate for the all condition was 0.65, 95% CI [0.51, 0.76]"
```

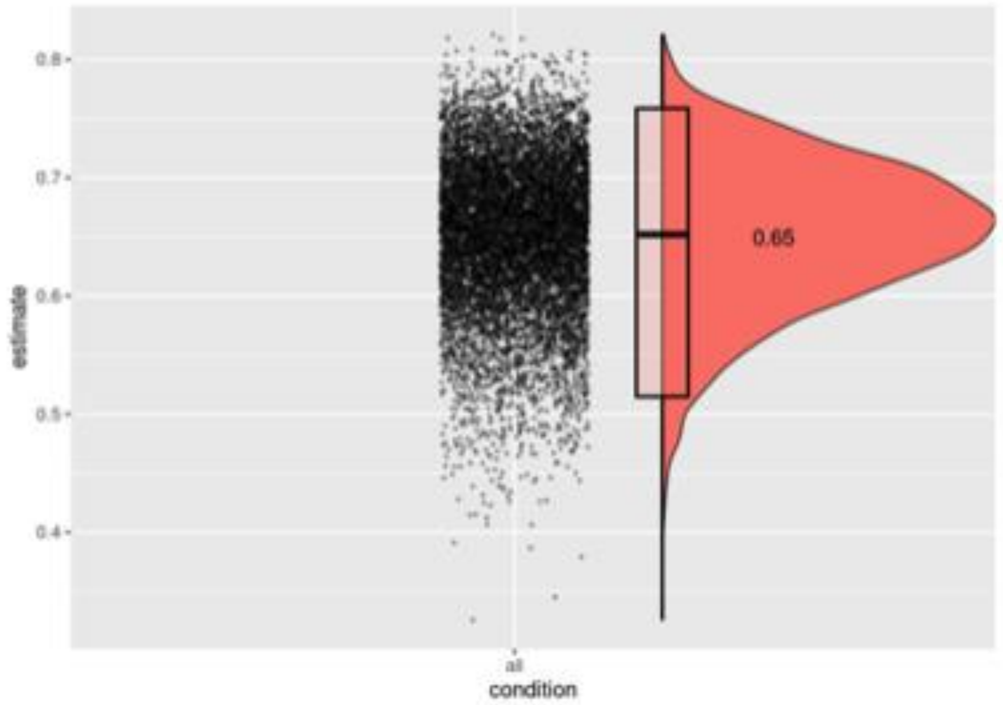


```
difference_executive <- splithalf(data = final_data,
                        outcome = "RT",
                        score = "difference",
                        halftype = "random",
                        permutations = 10000,
                        var.RT = "ant_RT",
                        var.participant = "participant_id",
                        var.compare = "congruency.code",
                        compare1 = "0",
                        compare2 = "1",
                        average = "mean",
                        plot = TRUE)
```

```
##
  |                                                                      
  |                                                                |   0%
  |                                                                      
  |=                                                               |   2%
  |                                                                      
  |==                                                              |   3%
  |                                                                      
  |===                                                             |   4%
  |                                                                      
  |===                                                             |   5%
  |                                                                      
  |====                                                            |   6%
  |                                                                      
  |=====                                                           |   7%
  |                                                                      
  |======                                                          |   8%
  |                                                                      
  |======                                                          |   9%
  |                                                                      
  |=======                                                         |  10%
  |                                                                      
  |========                                                        |  11%
  |                                                                      
  |========                                                        |  12%
  |                                                                      
  |=========                                                       |  13%
  |                                                                      
  |==========                                                      |  14%
  |                                                                      
  |==========                                                      |  15%
  |                                                                      
  |===========                                                     |  16%
  |                                                                      
  |============                                                    |  17%
  |                                                                      
  |============                                                    |  18%
  |                                                                      
  |=============                                                   |  19%
  |                                                                      
  |==============                                                  |  20%
  |                                                                      
  |===============                                                 |  21%
  |                                                                      
  |===============                                                 |  22%
  |                                                                      
  |================                                                |  23%
  |                                                                      
  |=================                                               |  24%
  |                                                                      
  |=================                                               |  25%
  |                                                                      
  |==================                                              |  26%
  |                                                                      
  |===================                                             |  27%
```

```
  |
  |===================                                                   |  28%
  |
  |====================                                                  |  29%
  |
  |=====================                                                 |  30%
  |
  |=====================                                                 |  31%
  |
  |======================                                                |  32%
  |
  |=======================                                               |  33%
  |
  |========================                                              |  34%
  |
  |========================                                              |  35%
  |
  |=========================                                             |  36%
  |
  |==========================                                            |  37%
  |
  |==========================                                            |  38%
  |
  |===========================                                           |  39%
  |
  |============================                                          |  40%
  |
  |============================                                          |  41%
  |
  |=============================                                         |  42%
  |
  |==============================                                        |  43%
  |
  |==============================                                        |  44%
  |
  |===============================                                       |  45%
  |
  |================================                                      |  46%
  |
  |=================================                                     |  47%
  |
  |=================================                                     |  48%
  |
  |==================================                                    |  49%
  |
  |===================================                                   |  50%
  |
  |====================================                                  |  51%
  |
  |=====================================                                 |  52%
  |
  |=====================================                                 |  53%
  |
  |======================================                                |  54%
  |
  |=======================================                               |  55%
```

```
|
|========================================                              |  56%
|
|========================================                              |  57%
|
|=========================================                             |  58%
|
|==========================================                            |  59%
|
|==========================================                            |  60%
|
|===========================================                           |  61%
|
|============================================                          |  62%
|
|============================================                          |  63%
|
|=============================================                         |  64%
|
|==============================================                        |  65%
|
|==============================================                        |  66%
|
|===============================================                       |  67%
|
|================================================                      |  68%
|
|=================================================                     |  69%
|
|=================================================                     |  70%
|
|==================================================                    |  71%
|
|===================================================                   |  72%
|
|===================================================                   |  73%
|
|====================================================                  |  74%
|
|=====================================================                 |  75%
|
|=====================================================                 |  76%
|
|======================================================                |  77%
|
|=======================================================               |  78%
|
|=======================================================               |  79%
|
|========================================================              |  80%
|
|=========================================================             |  81%
|
|==========================================================            |  82%
|
|==========================================================            |  83%
```

```
  |
  |===========================================================           |  84%
  |
  |============================================================          |  85%
  |
  |============================================================          |  86%
  |
  |=============================================================         |  87%
  |
  |==============================================================        |  88%
  |
  |==============================================================        |  89%
  |
  |===============================================================       |  90%
  |
  |================================================================      |  91%
  |
  |================================================================      |  92%
  |
  |=================================================================     |  93%
  |
  |==================================================================    |  94%
  |
  |===================================================================   |  95%
  |
  |===================================================================   |  96%
  |
  |====================================================================  |  97%
  |
  |===================================================================== |  98%
  |
  |===================================================================== |  99%
  |
  |======================================================================| 100%[1] "c
ondition all complete"
## [1] "Calculating split half estimates"
## [1] "split half estimates for 10000 random splits"
##   condition   n spearmanbrown SB_low SB_high
## 1       all 101          0.67   0.55    0.77
## [1] "this could be reported as: using 10000 random splits, the spearman-brown corr
ected reliability  estimate for the all condition was 0.67, 95% CI [0.55, 0.77]"
```

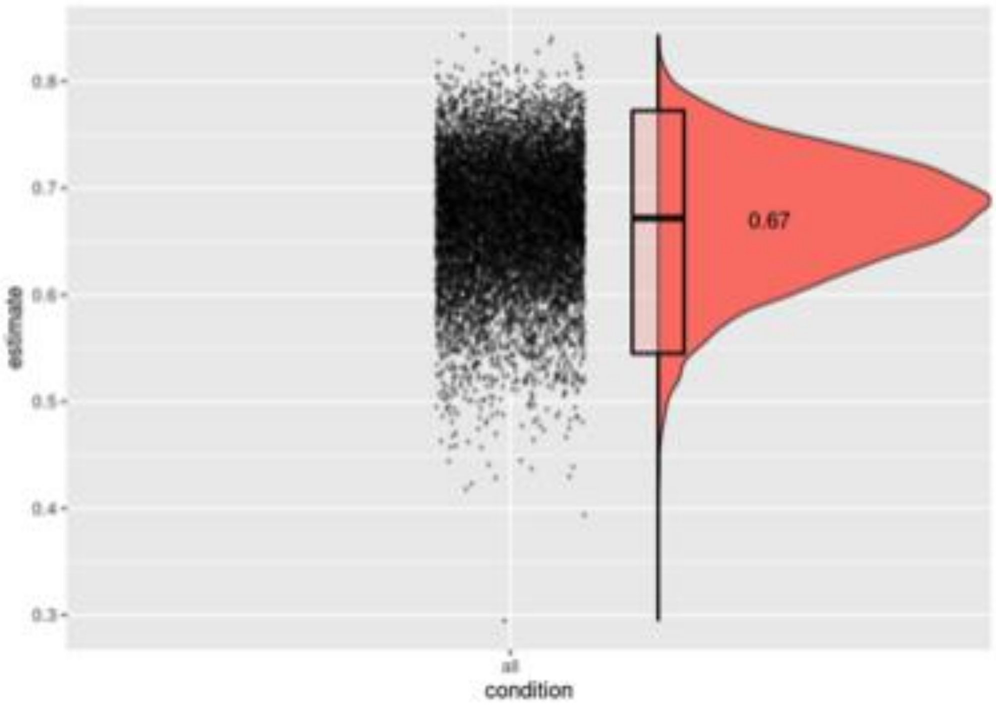
0.8
0.7
0.6
0.5
0.4
0.3
estimate
0.67
all
condition
